# Comparing mental imagery experiences across visual, auditory, and other sensory modalities

**DOI:** 10.1101/2023.05.15.540306

**Authors:** Alexander A Sulfaro, Amanda K Robinson, Thomas A Carlson

## Abstract

Although mental imagery is often studied as a visual phenomenon, it can occur in any sensory modality. Given that mental images may recruit similar modality-specific neural systems to those which support veridical perception, the properties of mental images may be constrained by the modality in which they are experienced. Yet, little is known about how mental images are experienced at all, let alone how such experiences may vary depending on the modality in which they occur. Here we explored how mental images are experienced in different modalities using an extensive questionnaire. Mainly focusing on visual and auditory mental imagery, we surveyed participants on if and how they experienced their thought content in a sensory way when thinking about the appearance or sound of the letter “O”. Specifically, we investigated temporal properties of imagined content (e.g. onset latency, duration), as well as spatial properties (e.g. apparent location), effort (e.g. ease, spontaneity, control), dependence on body movements (e.g. eye movements), interactions between real and imagined content (e.g. inner speech during reading), the perceived normality of imagery experiences, and how participants labeled their own experiences. Participants also ranked their mental imagery experiences in the five traditional sensory modalities and reported on the involvement of each modality during their thoughts, imagination, and dreams. Confidence ratings were taken for every answer recorded. Overall, visual and auditory experiences tended to dominate mental events relative to other sensory modalities. However, most people reported that auditory mental imagery was superior to visual mental imagery on almost every metric tested, except with respect to spatial properties. Our findings suggest that mental images are restrained in a similar matter to other modality-specific sensory processes in the brain. Broadly, our work also provides a wealth of insights and observations into how mental images are experienced by individuals, acting as a useful resource for future investigations.

## Introduction

Whether from gazing onto its whiskered head, patting its soft fur, or listening to its thrumming purr, each sensory modality can provide us with a stream of modality-specific information from which we can deduce the same modality-independent conclusion: we have encountered a cat. Yet, long after our cat has left the scene, we might still be able to see, hear, and pat it again in our thoughts. This phenomenon, where thoughts involve a sensory experience of what we are thinking about, even when it is not there, is known as *mental imagery*. Though mental imagery can hypothetically occur in any sensory modality, it is frequently studied in the visual and auditory modalities (including as a consequence of *inner* or *covert speech* when imagining speech sounds specifically). However, it is not clear how the subjective experience of mental imagery differs between these different sensory modalities. This limits the inferences which can be drawn about whether mental images are subject to the same modality-specific processing idiosyncrasies as their veridical counterparts, hindering our understanding of the mechanisms behind mental imagery. Consequently, here we sought to determine and compare the perceptual characteristics of imagined stimuli in multiple modalities, mainly focusing on visual and auditory experiences.

While the subjective experience of mental imagery in the visual and auditory modalities has been compared previously, comparisons are usually specific to one or two factors, or have been carried out with respect to comparing imagery *vividness* (e.g. Andrade et al., 2014; Gissurarson, 1992; Schifferstein, 2009; Sheehan, 1967; Switras, 1978; Talamini et al., 2022). However, as a construct, mental image vividness is infrequently defined or disambiguated such that ratings of imagery vividness have quite limited explanatory power (Dean & Morris, 2003; Richardson, 1988). It is therefore difficult to draw interpretable conclusions from comparing imagery vividness in different modalities. More interpretable measures of how mental images are experienced do exist, such as for visual imagery (see Pearson et al., 2013 for an extensive list) and auditory imagery (see Alderson-Day & Fernyhough, 2015; Hubbard, 2010 for reviews). Yet, these measures tend to be tailored to a single modality rather than for comparisons between modalities. Hence, it is still largely unknown how the quality of mental imagery is affected by the modality in which it occurs.

Here, we sought to surmount these existing challenges in an online questionnaire asking participants to report on how they experience their thoughts, focusing on visual and auditory imagery in detail. Crucially, we compared visual and auditory mental imagery on the same set of questions, probing the same properties, using interpretable metrics, and using a single simple imagery stimulus that can be experienced both in a visual and auditory way: the letter “O”. This approach allowed us to assess a multitude of properties concerning how stimuli are imagined, including where imagined thought content seems to be located, whether imagery occurs automatically, immediately, and indefinitely, and whether imagined content fluctuates in visibility or audibility. In contrast to vividness-based investigations, many of the traits that we probe are precisely defined and can be similarly applied to measuring real-world stimulus characteristics. This allows for imagined experience to be compared in a highly-interpretable way across different sensory modalities.

We also incorporated a range of explicit comparisons to allow respondents to unambiguously report which of five sensory modalities were most dominant with respect to a number of criteria concerning mental imagery properties (e.g. which modality’s imagery started most quickly, occurred most easily, or required the least body movements). Further, we investigated the prevalence of imagination, thoughts, and dreams in each sensory modality, how people evaluate and define their own imagery experiences, and how confident respondents were about each of their responses. Ultimately, we observed modality-specific biases in mental imagery which, for most respondents, generally tended to favour more refined auditory imagery rather than visual imagery. We account for our findings by considering modality-specific differences in the brain’s capacity to process and simulate sensory information.

## Methods

### Participants

In total, 590 undergraduate psychology students from the University of Sydney participated in the study. Research was conducted with ethics approval from the University of Sydney ethics committee. Participants provided informed consent before participating. Each participant received course credit for their participation which was automatically granted to the student upon completion of all required parts of the study. Despite knowing that many would be excluded from analysis using our pre-defined criteria, our study was made broadly available to first-year research participants so that students would be able to complete components of their coursework related to research participation. 120 participants were excluded from analysis if they reported that they had a known neurological, psychological, or psychiatric condition that was likely to affect their thoughts, imagination, dreams, or perception. 111 additional participants were also excluded for failing any of the three basic attention checks present, and 1 extra individual was excluded due to an error that allowed them to avoid completing all required parts of the questionnaire. Pilot testing indicated that proper completion of the questionnaire required approximately 30 minutes and likely could not be completed properly in less than 20 minutes. Hence, 63 further participants were excluded for completing the study in less than 20 minutes.

To ensure that participants correctly interpreted our descriptions of sensory thought content as mental imagery, 80 participants were conservatively excluded from analysis if English was not their dominant language, nor the language they were most comfortable reading in. Similarly, to ensure participants did not confuse descriptions of mental imagery with hallucinations given the potential ambiguity surrounding descriptions of sensory thought content (e.g. hearing your thought content), 14 participants were excluded from most analyses because they chose the term “Hallucination” to describe the phenomenon they most commonly have when they in some way see or hear what they are thinking about. However, these individuals were still included in some specified analyses, such as when evaluating participant beliefs about their own experiences. After exclusions, responses from a total of 201 participants were analysed, or a total of 215 participants for analyses including those describing their experience as a hallucination (median age = 19, female = 140, male = 72). Although we excluded a significant number of participants from the analyses here, the full dataset is available for analysis without exclusions (Supplementary S2).

### Materials

The questionnaire was created using Qualtrics in-browser software (www.qualtrics.com). Participants accessed and completed the study within their internet browsers. Qualtrics hardware detection measures were used to prevent participants from being able to start the study on a mobile device. Analysis was conducted using custom Python code (v3.9.15) in JupyterLab (v3.4.4), with hypothesis tests calculated using the scipy (v1.9.3) and statsmodels (v0.13.2) Python packages.

### Procedure

Participants were asked to complete a questionnaire concerning how they experienced their thoughts. The full questionnaire is given in Supplementary S1.1. Participants were given a battery of questions individually probing how they experienced the appearance of thought content and the sound of thought content (e.g. “When did your thought content first seem to become visible to you in some way?”). These questions were designed to ascertain the properties of mental images in a similar way to real stimuli, probing properties such as the onset latency, duration, spatial location, and temporal consistency of sensory thought content. We also probed participant beliefs about their thought processes, such as whether they considered seeing or hearing thought content in some way to be a normal process.

Additionally, we asked questions explicitly comparing how visual and auditory thought content is experienced, as well as comparing mental imagery across the five traditional senses of sight, hearing, smell, taste, and touch (e.g. “Which is the easiest to initiate in some way?”). We further explored the involvement of the five traditional senses in thoughts, imagination, and dreams. Demographic information was collected along with information about the environmental and mental context in which participants completed the study (e.g. a description of their environment, environmental brightness and loudness levels, and time since waking).

For most questions, participants were explicitly instructed to think about either the appearance or sound of the letter “O”, with their eyes specified as opened or closed, and sometimes for a specified duration of time. Participants were instructed to respond honestly to questions based on their actual experience in each instance of thinking rather than based on what prior experiences had let them to believe. Where logical, questions were matched between visual and auditory modalities. Most questions were single-response multiple-choice questions. An option was always included to allow participants to indicate that they had no sensory experience of their thought content where appropriate. All questions throughout the questionnaire, apart from demographic and optional free-text questions, also required the participant to rate how sure of their response they were on a 6-point scale from “Completely unsure” to “Completely sure”.

Participants were encouraged to complete the questionnaire in a quiet place and were warned that attention checks would be present within the study. No actual penalty was given to the student if they failed to complete the study or any of the three attention checks distributed throughout the questionnaire. There was no time limit to complete the study.

### Assessing sensory thought content rather than “mental imagery”

Note that, despite investigating mental imagery, we specifically avoided using the term “mental image” throughout the experiment, instead opting to ask participants whether they had a visual or auditory experience of their thought content, or if they saw or heard their thought content in some way. This was done for several reasons:

Firstly, “mental images” are commonly associated with the visual modality, and naïve participants may not consider sensory thought content in other modalities to be “mental imagery.” Among other issues, using “mental image” terminology could therefore result in reduced confidence when responding in non-visual modalities purely because of a lack of familiarity with what a non-visual mental imagery experience entails. Secondly, describing mental images in terms of sensory thought content may reduce the likelihood that participants answer based on their preconceived notions of what “mental imagery” is. For example, our approach should be more sensitive to individuals who have quasi-sensory experiences of their thoughts but who may not report having mental imagery because they assume mental imagery involves an eidetic or hallucinatory (i.e. unambiguously-sensory) experience phenomenologically equivalent to veridical perception. Our approach is also sensitive to probing individuals who truly have no sensory experience of their thoughts, but still believe that they can do “mental imagery” because they assume that forming a “mental image” is merely a metaphor, figure of speech, or visual device in film and television rather than a literal sensory experience. Reports of these mix-ups extend back to the nineteenth century (Galton, 1880) and continue today such that it is now virtually a trope for one to discover that “mental imagery” is not a metaphorical term. Indeed, at the time of writing, two of the top three posts of all time on www.reddit.com/r/aphantasia, a forum discussing imagery extremes with more than 50,000 users, relate to this exact situation. One of our respondents also reported a similar predicament with regards to auditory imagery, stating *“I used to think internal monologues were only ever used in movies to help verbalise a characters* [sic] *thoughts, not a real experience”*. For these reasons, we believe that avoiding explicitly asking participants about “mental images” may provide more accurate subjective reports.

### Analysis

In general, this study investigated how visual and auditory imagery were experienced on a number of properties. Because most questions had both a visual and auditory equivalent, we were able to compare how imagery was experienced in each modality using paired responses. For paired categorical responses, such as for comparing single-response multiple-choice questions, differences in responding patterns could be tested using a Bhapkar homogeneity test. When comparing questions where only two responses were possible per modality (2×2), this test is equivalent to a McNemar’s test and differences between the modalities can be directly attributed to differences between specific pairs of response proportions. However, where more than two response options are available, this test cannot be used to ascertain which specific differences in response proportions are driving any general differences between the modalities. In these cases, response proportions were grouped for each modality into target vs. non-target responses before conducting a Bhapkar (McNemar’s) test on the collapsed 2×2 data, and then repeating this analysis for each possible response option. For instance, for a question with response options A, B, C, and D, to judge if there was a significant difference between modalities in terms of the proportion of respondents choosing option A, we would run a McNemar’s test comparing the proportion of responses for option A and the proportion of responses for options B, C, and D combined (i.e. *not-*A responses). To assess differences for each possible response option, this process was repeated for each response option (e.g. B and not-B, then C and not-C, then D and not-D responses), and a Bonferroni correction was applied based on the number of 2×2 tests conducted (in this example, 4).

Hypothesis tests were two-sided unless otherwise specified, with an alpha level of 0.05. Significance was determined according to the Bonferroni-corrected p-value, where applicable. Raw p-values are reported in the main text, however inferences and figure annotations are derived from corrected p-values. Note that our entire Results section was generated as dynamic text in a JupyterLab notebook (Supplementary S3) such that all hypothesis test outcomes can be transparently traced back to their underlying data and reanalysed if desired.

### Unimodal-vs. multimodal-imager comparisons

Importantly, we note that differences in responding between modalities can be caused by two distinct factors: differences in how mental imagery is experienced (e.g. shorter vs. longer imagery durations), and differences in the frequency with which mental imagery is experienced at all (e.g. having imagery vs. not having imagery). We must be cautious to avoid conflating differences in the incidence of mental imagery between modalities with differences in the properties of mental imagery between modalities. Therefore, when comparing the properties of imagery between modalities, we included data only from those who reported experiencing mental images in both modalities examined for that question (multimodal imagers). We excluded those who reported imagery in only one of the modalities examined (unimodal imagers), and those who reported no imagery in any modality examined (non-imagers), except where the incidence of imagery itself was relevant.

## Results

### Prevalence of mental imagery

We first sought to determine the proportion of individuals who experienced mental imagery at all in either the visual or auditory modality. We asked respondents to think about the appearance or sound of the letter “O” and report whether or not they in some way saw or heard their thought content during this specific instance of thinking. When asked explicitly, 87.6% of included respondents reported a visual experience of their thought content and 75.1% reported an auditory experience of their thought content.

However, there was considerable variation in how well individuals were able to generate mental imagery over the course of the questionnaire. During the questionnaire, respondents were frequently asked to attempt imagery and report only on their experience at that moment. Given that there were 30 questions probing the properties of imagery per visual or auditory modality, participants were therefore presented with 30 opportunities to report whether they experienced imagery at the time of answering. Across these questions, 51.8% of participants reported no visual experience of their thought content at least once, while 1.0% never reported a visual experience of their thought content. For audition, 31.9% reported no auditory experience of their thought content at least once and 1.0% never reported an auditory experience of their thought content. On average, participants reported no sensory experience on 3.6% of visual questions, and 4.6% of auditory questions. Overall, this variability reflects that mental imagery generation may be far from reliable for many individuals.

### Eyes-open vs. eyes-closed mental imagery

We investigated whether the incidence of imagery varied depending on whether a participant conducted imagery with their eyes open or closed. Our questionnaire included two opportunities for this comparison to be made per modality, with the average incidences across these two comparisons shown in Figure 1A. In the visual modality, more respondents reported imagery with their eyes closed rather than open (comparison 1: Χ²(1) = 39.2, p < .001; comparison 2: Χ²(1) = 32.4, p < .001). Yet, eye-closure did not affect the likelihood of hearing one’s thought content (comparison 1: Χ²(1) = 1.3, p = .255; comparison 2: Χ²(1) = 1.3, p = .255).

**Figure 1.**
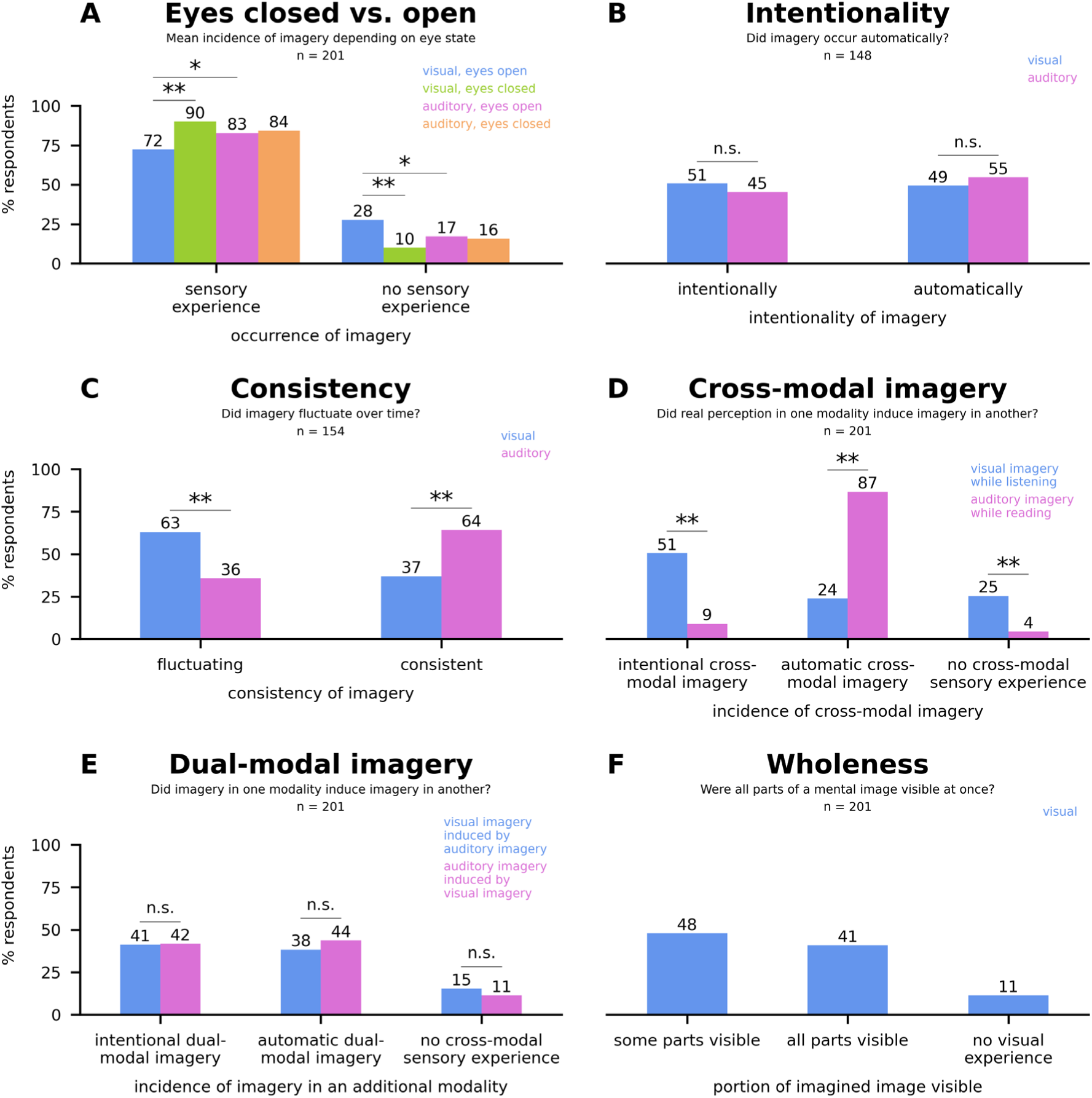
Incidence and properties of visual and auditory mental imagery. *Note.* **A-F.** Multiple-choice response distributions. Annotations indicate significant differences between modalities in the proportion of times a specific response option was chosen, after Bonferroni corrections: n.s. = p > 0.05, * = 0.001 < p ≤ 0.05, ** = p ≤ 0.001. **A.** For each modality, there were two opportunities within the questionnaire where eyes-open and eyes-closed imagery could be compared. Proportions shown here are averaged over these two opportunities. Hypothesis tests yielded the same inferences for both comparisons. For clarity, non-significant results are not shown in this plot, however main effects of modality and eye-state were tested in all combinations. **B.** Responses shown only from those who experienced imagery in both visual and auditory modalities (multimodal imagers). **C.** Responses shown from multimodal imagers only. **D.** Responses pooled from unimodal and multimodal imagers. **E.** Responses shown for participants who could at least generate each of visual and auditory imagery separately **F.** Responses shown in the visual modality only.

### Intentionality: Did mental imagery occur automatically?

We probed whether a sensory experience occurred intentionally or automatically as part of thinking. For those reporting an imagery experience in both modalities (i.e. multimodal imagers; Figure 1B), there was no significant difference in whether imagery occurred automatically or intentionally between modalities (Χ²(1) = 0.8, p = .370). However, for those reporting only either visual imagery or auditory imagery (unimodal imagers, n = 53), responding differed between modalities (Χ²(2) = 23.8, p < .001), with visual imagery 10.4x more frequently reported as intentionally-constructed compared to auditory imagery (39.6% vs. 3.8% respectively; Χ²(1) = 22.3, p < .001). Yet, there was no significant difference between modalities in the frequency with which unimodal imagers reported visual or auditory imagery to be automatic when thinking (17.0% vs. 24.5% respectively; Χ²(1) = 0.7, p = .390). Hence, for those who experienced imagery in both modalities, imagery was just as likely to be generated in the same way in either modality. Yet, if visual imagery was the only form of imagery experienced, it was most likely to be intentionally constructed, rather than automatically. If only auditory imagery was experienced, it was most frequently automatic. However, differences between modalities for unimodal imagers may be at least partially attributed to differences in the proportion of individuals who failed to experience visual and auditory imagery at all for this question (43.4% vs. 71.7% for each modality, respectively; Χ²(1) = 5.5, p = .019).

### Consistency: Did mental imagery fluctuate over time?

Participants also reported on whether their thought content was consistently visible or audible during a 5-second period, or whether it fluctuated in how visible or audible it was (Figure 1C). Responding differed between modalities, with imagined sounds more frequently reported as consistent and imagined images more often as fluctuating (Χ²(1) = 27.3, p < .001).

### Cross-modal imagery: Did real perception in one modality induce mental imagery in another?

We asked participants whether the real perception of environmental stimuli using vision or audition (i.e. reading written words or listening to spoken words) would induce mental imagery of the same thought content in the other modality (i.e. mentally hearing or seeing the words). We refer to this experience as cross-modal mental imagery. Across all respondents, there were significantly more participants who reported that reading induced auditory imagery (cross-modal auditory imagery) than there were reporting that listening induced visual imagery (cross-modal visual imagery; Χ²(1) = 40.8, p < .001). The distribution of responses regarding cross-modal imagery for all respondents is shown in Figure 1D. To determine whether one form of cross-modal imagery was more or less automatic than the other, we assessed responses only from participants who experienced cross-modal mental imagery in both modalities (i.e. multimodal imagers). For these respondents, cross-modal auditory imagery was far more likely to be automatic than cross-modal visual imagery, with 2.8x more respondents reporting that they heard words that they were reading automatically compared to automatically seeing words that they were hearing (Χ²(1) = 172.0, p < .001). Participants could picture words that they had heard, but generally only if they intended to do so.

### Dual-modal imagery: Did mental imagery in one modality induce mental imagery in another?

We asked participants whether visual or auditory mental imagery in one modality also induced mental imagery in the other modality at the same time. We call this experience dual-modal mental imagery. For respondents who could generate imagery in both modalities separately, imagining in another modality simultaneously was also possible, if not automatic (Figure 1E). Responding did not significantly differ depending on whether auditory imagery was added to visual imagery or whether visual imagery was added to auditory imagery, both for participants who could construct imagery in both modalities separately (Χ²(2) = 5.8, p = .055), or only in one of the two modalities separately (Χ²(2) = 5.3, p = .070).

### Wholeness: Were all parts of a mental image visible at once? (visual only)

We also asked respondents to report whether all parts of their thought content were visible at any given time, or if only some parts were visible (Figure 1F). Partial and whole imagery were reported roughly equally, with slightly more respondents reporting only partial visibility. We did not ask an equivalent question regarding the imagined auditory stimulus as it involved only a single phoneme.

### Peak: When was mental imagery most sensory within a thought period?

Participants reported when their thought content was most visible or audible during a given 5-second period where they actively attempted imagery. For multimodal imagers (Figure 2A), the point when imagery was most sensory significantly differed depending on imagery modality (Χ²(3) = 71.6, p < .001). Thought content was most frequently reported as being most sensory during the middle of the imagery period for visual imagery, significantly more frequently than for auditory imagery (Χ²(1) = 20.7, p < .001). Auditory imagery, however, was most frequently reported as equally audible throughout the imagery period, by 2.6x as many respondents as for visual imagery (Χ²(1) = 49.7, p < .001).

**Figure 2.**
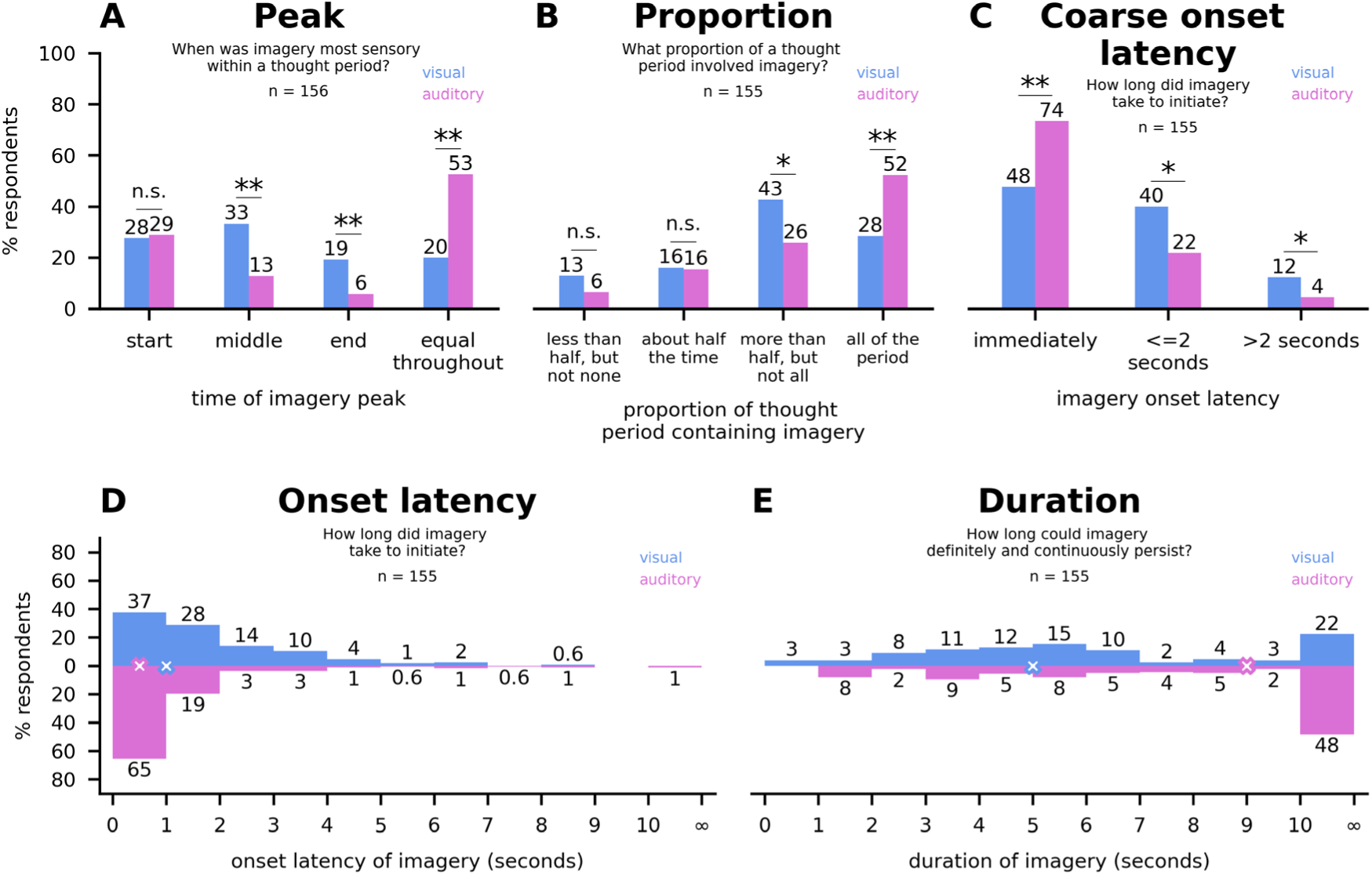
Temporal properties of visual and auditory mental imagery. *Note.* **A-C.** Multiple-choice response distributions. Annotations indicate significant differences between modalities in the proportion of times a specific response option was chosen, after Bonferroni corrections: n.s. = p > 0.05, * = 0.001 < p ≤ 0.05, ** = p ≤ 0.001. Responses shown only from those who experienced imagery in both visual and auditory modalities (multimodal imagers). **D-E.** Histograms of responses from multimodal imagers taken using a sliding scale. Responses are grouped into 1s bins (lower-limit inclusive and upper-limit exclusive), except for the final bin, which shows the proportion of all responses of ≥ 10s. Crosses indicate the median response for each modality. **D.** Auditory imagery had a significantly shorter onset latency than visual imagery. **E.** Auditory imagery has a significantly longer duration than visual imagery.

### Proportion: What proportion of a thought period involved mental imagery?

We asked participants to report the proportion of time in which sensory content was visible or audible during a 5-second period where they actively attempted imagery. For multimodal imagers (Figure 2B), the proportion of the period which involved imagery significantly differed depending on imagery modality (Χ²(3) = 27.8, p < .001). Thought content was most frequently reported as visible for more than half, but not all, of the thought period for visual imagery, significantly more frequently than for auditory imagery (Χ²(1) = 10.6, p = .001). Yet, auditory imagery was most frequently reported as occurring throughout the entire thought period, by 1.8x as many respondents as for visual imagery (Χ²(1) = 25.3, p < .001).

### Onset latency: How long did mental imagery take to initiate?

We asked participants to report how long it took for thought content to become visible or audible, both using a multiple choice question, and using a sliding numerical scale. For categorical responses from multimodal imagers (Figure 2C), the onset latency of imagery significantly differed depending on modality (Χ²(2) = 23.5, p < .001). While imagery was most frequently reported as beginning immediately for both modalities, 1.5x as many respondents reported immediate auditory imagery compared to visual imagery (Χ²(1) = 21.7, p < .001). Using a sliding scale, participants could report their imagery onset latency with a numerical value from 0-10 seconds or otherwise report that their imagery took 10 or more seconds to initiate (Figure 2D). For multimodal imagers, a two-sided Wilcoxon signed-rank test showed a significant difference in onset latency between visual and auditory imagery (Z = −4.5, p < .001). Onset latencies for auditory imagery (median = 0.5s) were shorter than for visual imagery (median = 1.0s). Overall, these findings indicate auditory imagery began more quickly than visual imagery.

### Duration: How long could mental imagery definitely and continuously persist?

We asked participants to report how long they could definitely and continuously see or hear their thought content. Using a sliding scale, participants could report their imagery duration with a numerical value from 0-10 seconds or otherwise report that their imagery could last for 10 or more seconds (Figure 2E). For multimodal imagers, a two-sided Wilcoxon signed-rank test showed no significant difference in imagery duration between modalities (Z = −5.0, p < .001). However, visual imagery (median = 5.0s) tended not to last as long as auditory imagery (median = 9.0s), and there were significant differences between modalities in the proportion of multimodal imagers reporting that their imagery could last for 10 or more seconds, as opposed to below 10 seconds (Χ²(1) = 36.3, p < .001). Indeed, auditory imagery (48.4%) was 2.2x as frequently reported as lasting 10 or more seconds than visual imagery (22.3%). Overall, these findings suggest that most respondents could generate auditory imagery for a longer period than visual imagery.

### Movement-dependence: Which body movements were required for mental imagery?

Previous research indicates that mental imagery often involves motor activity related to the content being imagined. For instance, eye movements during visual imagery tend to relate to the spatial content of visual mental images (Fourtassi et al., 2017), while muscle movements related to vocalisation are detectable when imagining sounds (Pruitt et al., 2019). Here we probed whether or not participants believed that they required any such movements to perform imagery in each modality. Specifically, we asked participants to report whether they could see or hear their thought content while remaining completely still, or if they had to move some part of their body in some way for imagery to occur (Figure 3A). We also probed which specific body parts were most frequently reported as necessary for imagery in each modality (Figure 3B). The majority of multimodal imagers reported that body movements were not required for imagery in both modalities. However 2.5x more multimodal imagers reported requiring movement for visual, but not auditory, imagery (Χ²(1) = 15.8, p < .001).

**Figure 3.**
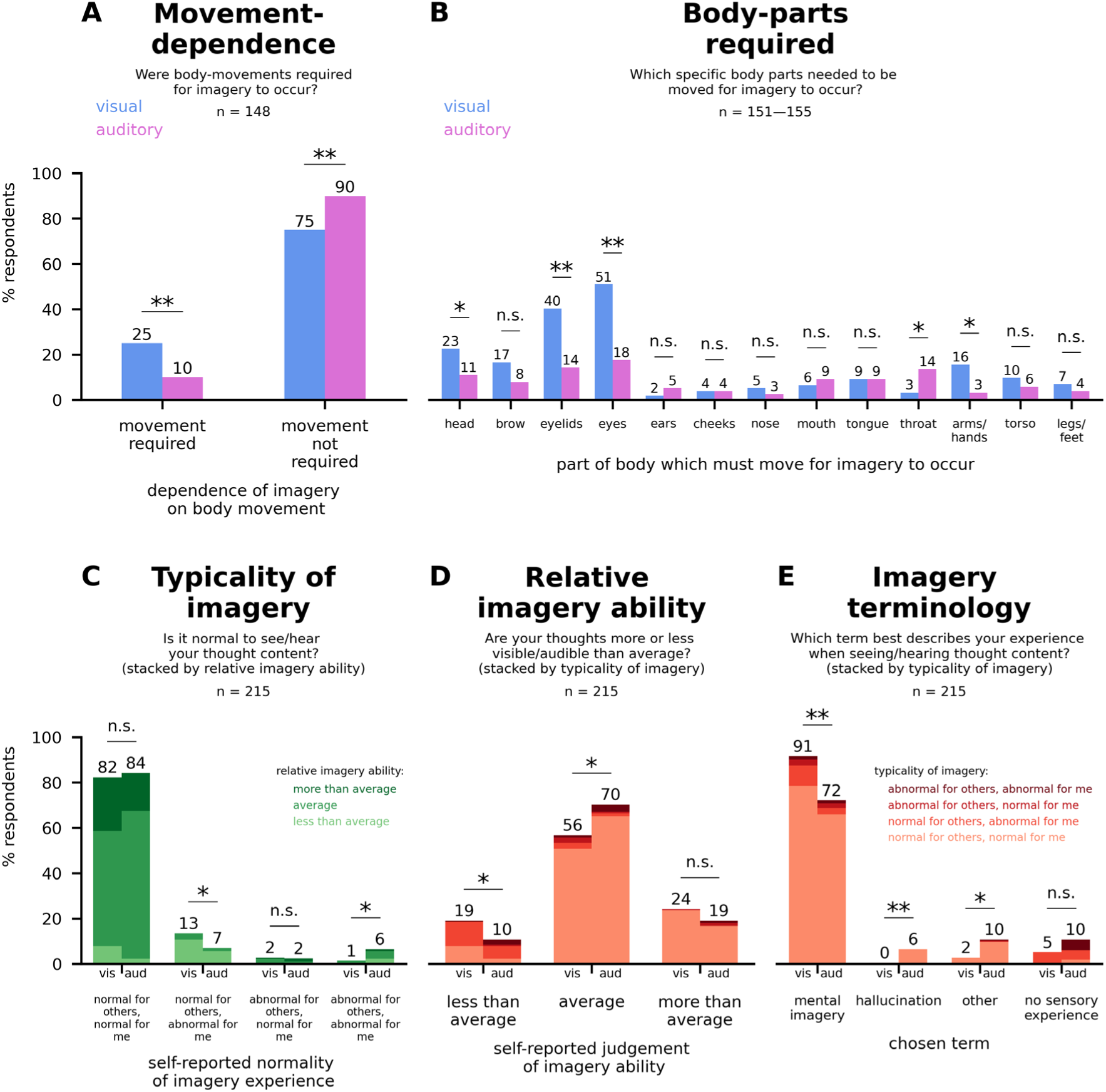
Body movement during visual and auditory mental imagery, and beliefs about mental imagery. *Note.* **A-E.** Multiple-choice response distributions. Annotations indicate significant differences between modalities in the proportion of times a specific response option was chosen, after Bonferroni corrections: n.s. = p > 0.05, * = 0.001 < p ≤ 0.05, ** = p ≤ 0.001. **A.** Responses shown only from those who experienced imagery in both visual and auditory modalities (multimodal imagers). **B.** Responses shown only for multimodal imagers. Participants answered a separate question for each body part. **C-E.** Beliefs about mental imagery, with responses in each column split by response choice in neighbouring questions. Statistics apply only to comparisons of proportions for the main question; the breakdown of each column by secondary response was not assessed statistically. Responses presented in C-F include those from respondents who reported mental imagery as a hallucination in E.

Despite most multimodal imagers reporting that movement was not required for imagery when asked broadly, many more respondents reported that body movements were required when they were asked about the involvement of individual body parts one-by-one. Figure 3B shows the proportion of multimodal imagers who reported that a particular body movement was necessary for imagery. Respondents may have had difficulty introspecting on the involvement of body parts during imagery until they were required to systematically test a list of body parts. They may also have interpreted the phrasing of the explicit question concerning whether imagery occurred “even if you were remaining completely still” to refer to body, but not head, stillness. For visual imagery, the three most frequently reported body movements, from most-reported to least-reported, were the eyes, eyelids, and head. Note that, for visual imagery, the majority of multimodal imagers reported that seeing thought content required eye movements (51.0%) when explicitly asked about eye movements, yet only a minority reported that movements of any form were required when asked about body movements generally. For auditory imagery, few participants reported that body movements were necessary when asked broadly or specifically, with the three most frequently reported body movements being those of the eyes, eyelids, and throat. We do note that, for both modalities, participants were first instructed to conduct imagery with their eyes closed and then comment on their experience at the time. However, participants were explicitly instructed not to report body movements purely associated with closing their eyes for the task.

In either modality, 3.0% or fewer multimodal imagers reported that they needed to move a body part which was not listed in Figure 3B. However, almost all open text responses (Supplementary S1.8) regarding the unlisted body part referred to body parts already listed, or related to changes indirectly related to imagery (e.g. moving to change posture or put something away, or moving in order to keep breathing). One person mentioned that they had to rub their closed eyes to produce visual imagery. Jaw movements and stomach or diaphragm movements were each mentioned once for auditory imagery.

### Coarse position and location control: Where did mental images seem to be located, and could the apparent location of mental images be controlled?

We investigated the spatial properties of imagery, namely whether the spatial location of seen or heard thought content could be controlled, and where such thought content most frequently seemed to originate from or be spatially positioned at. We investigated each property during both eyes-open and eyes-closed imagery. For respondents that experienced imagery in both modalities and eye states, a significantly larger proportion of respondents could control the apparent position of seen thought content rather than heard thought content (Figure 4A), both when eyes were open (Χ²(1) = 17.4, p < .001) and closed (Χ²(1) = 7.3, p = .007). Eye-closure did not significantly affect the proportion of these respondents reporting that they could control their visual imagery (Χ²(1) = 2.9, p = .086) or auditory imagery (Χ²(1) = 0.0, p > .999).

**Figure 4.**
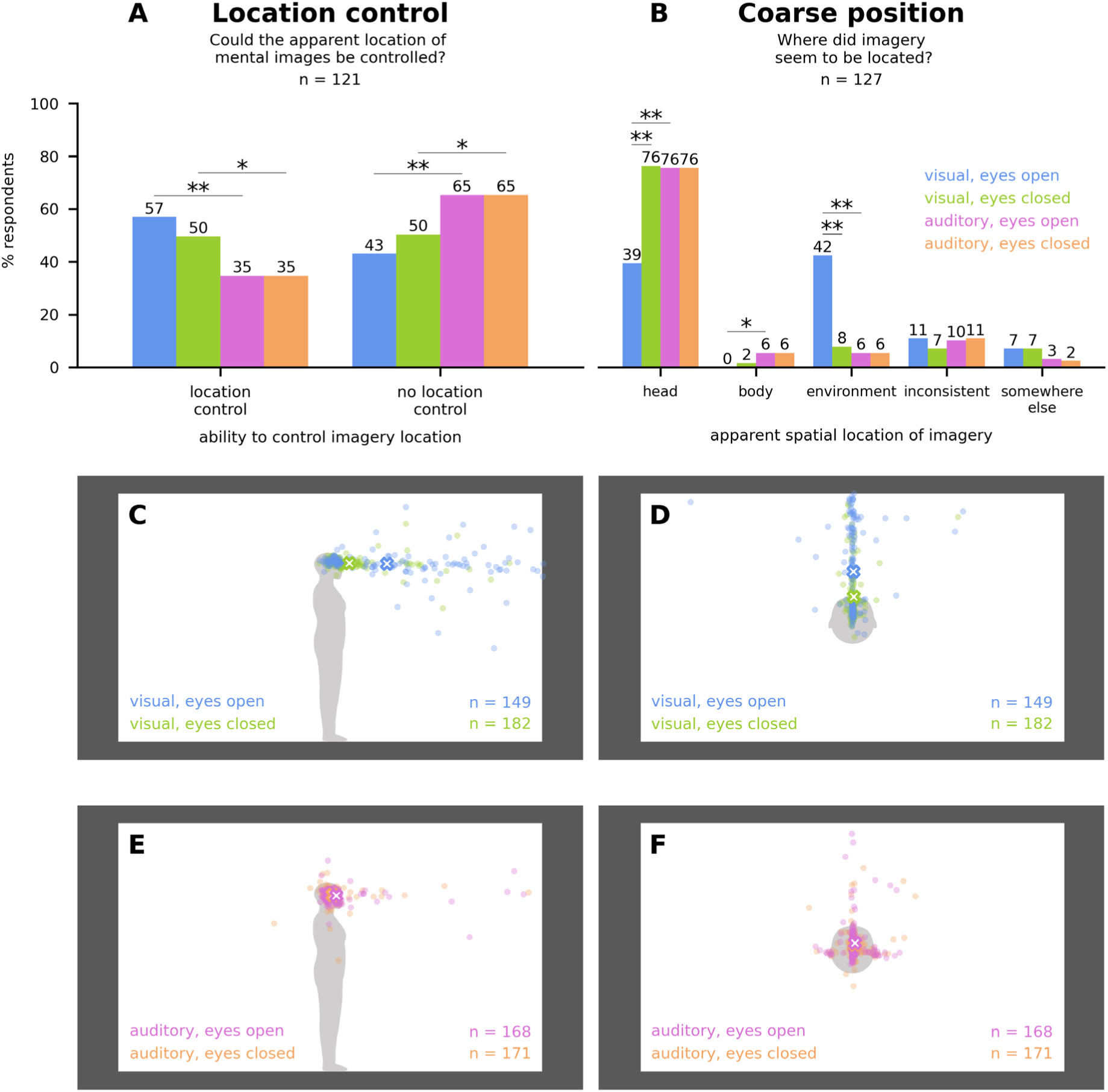
Spatial properties of visual and auditory mental imagery. *Note.* **A-B.** Multiple-choice response distributions. Annotations indicate significant differences between modalities in the proportion of times a specific response option was chosen, after Bonferroni corrections: * = 0.001 < p ≤ 0.05, ** = p ≤ 0.001. Responses shown only from those who experienced imagery in both visual and auditory modalities (multimodal imagers). For clarity, non-significant results are not shown, however main effects of modality and eye-state were tested in all combinations. **C-F.** Distribution maps of precise locations of imagery. Each dot indicates a single respondent’s chosen location for where imagery most commonly seems to be located or originate from. Crosses indicate mean positions. Participants were not given the option to place a marker if they had indicated that they did not have a sensory experience of their thoughts when previously asked about the coarse location of their imagery categorically. **C-D.** Visual imagery was projected forward, away from the head, when eyes were open, significantly more so than when eyes were closed. **E-F.** Eye state did not significantly affect the location of auditory imagery.

As for the reported position of imagined sounds and images, when asked in a multiple choice question, most individuals reported that imagined sounds and images were located within their head, except for during eyes-open visual imagery (Figure 4B). Eyes-open visual imagery was significantly less frequently reported as occurring in the head compared to eyes-closed visual imagery (Χ²(1) = 58.8, p < .001), and relative to eyes-open auditory imagery (Χ²(1) = 44.1, p < .001). Instead, images visualised with eyes open were significantly more likely to be reported as being positioned in the environment relative to eyes-closed imagined images (Χ²(1) = 64.3, p < .001), or eyes-open imagined sounds (Χ²(1) = 57.9, p < .001).

### Precise spatial position of mental imagery

If participants reported a visual or auditory experience of their thoughts for a particular eye state, we also asked them to indicate the apparent location of the thought content they were seeing or hearing by placing a single marker on an image of a human in a room, as viewed from the side (Figure 4C & E) or above (Figure 4D & F). This allowed us to more precisely visualise where in space imagined images and sounds were perceived as originating from or being located at. In general, imagined content was perceived either in the head, directly in front or beside the head, or projected forward from the head. When eyes were open, the content of visual mental images appeared to shift forward compared to when eyes were closed, projecting away from the head and into the environment. We were able to estimate this distance in centimetres by assuming an average height of 171.0cm and an average head circumference of 56.0cm for the figures portrayed in our images. Two-tailed Wilcoxon signed-rank tests confirmed a statistically significant forward projection of visual imagery when eyes were open rather than closed, with a mean projection of approximately 35.5cm along the horizontal axis in Figure 4C (Z = −6.2, p < .001) and 10.2cm along the vertical axis in Figure 4D (Z = −5.8, p < .001). Levene tests also showed that the reported imagery position was more variable along these axes when eyes were open relative to closed, with standard deviation increases of 29.0cm (W = 69.7, p < .001) and 7.0cm (W = 91.5, p < .001) respectively. Up-down position variance also increased with eye opening along the vertical axis of Figure 4C, with a standard deviation increase of 7.0cm (W = 10.5, p = .001). All other comparisons of variance or position between an eyes-open and eyes-closed state within a modality were non-significant after Bonferroni correction (p > .013).

### Typicality, ability, and terminology: Is mental imagery normal, how do individuals compare to others, and which term best describes their experience?

We asked participants to report on whether they considered seeing or hearing their thoughts in some way to be a normal experience for themselves and for others (typicality; Figure 3C), whether they considered their thoughts to be more or less sensory than average (ability; Figure 3D), and which term they would use to best describe the experience they most commonly have when seeing or hearing their thought content (terminology; Figure 3E). Note that participants who described their experience as a hallucination were included in the analyses of these three questions despite being excluded from analyses in previous questions. Although most respondents thought that seeing or hearing their thought content was a normal experience for others and themselves, there was a slight bias towards reporting that seeing thought content was normal, even if it was not a normal event for the participant, while hearing thought content was slightly more likely to be considered an abnormal process for all. Most participants felt that their imagery abilities were average, with slightly more participants suspecting that their visual imagery, rather than auditory imagery, was less sensory than average. Note that many of these individuals still recognised visual imagery as a normal process for others even if they had a low self-reported visibility of their own thought content. Significant differences are shown in Figure 3.

Throughout this study, we deliberately avoided asking participants to report on “mental imagery” for reasons previously detailed (see Methods). Instead, we asked participants to report on whether they in some way saw or heard their thought content, or had a visual or auditory experience of their thought content. We confirm that most participants understood this phenomenon to be synonymous with mental imagery (Figure 3E), although a substantial proportion of respondents described hearing thought content as a hallucination, or using another term, despite using “mental imagery” to describe seeing thought content. This is not unexpected given that “mental imagery” is traditionally associated with the visual modality. Alternative terms provided by respondents for the common experience of hearing one’s thoughts are listed in Supplementary S1.8, with most still describing the phenomenon of auditory mental imagery despite not using the term explicitly, or using an auditory-specific term instead e.g. an inner or internal voice or monologue, hearing one’s thoughts, or hearing a voice in their head.

### Explicit comparisons of mental imagery in different modalities

Until now, we have compared imagery modalities by asking participants to make judgements about their thought content in each modality separately and then comparing their judgements after the fact. However, we also asked participants to make explicit judgements comparing imagery properties between modalities. In Figure 5A, we show the distribution of responses from when participants were explicitly asked to compare their experiences and judge which of visual or auditory thought content most met a given criteria. Participants could also report that sensory thoughts in both modalities were experienced in the same way with respect to the criterion being inquired about, or whether they had no sensory experience at all (e.g. “Which is the easiest to do in some way?” “Seeing my thought content in some way”, “Hearing my thought content in some way”, “There is no difference”, or “I do not see or hear my thought content in any way”). We also asked participants to make judgements about not just visual and auditory thought content, but sensory thought content in the five traditional senses (sight, sound, smell, taste, and touch). The distribution of these judgements is shown in Figure 5B. Note that, for these latter judgements, participants could choose multiple sensory modalities if they deemed that the criteria could apply to multiple modalities equally, or they could report that they had no experience in any modality.

**Figure 5.**
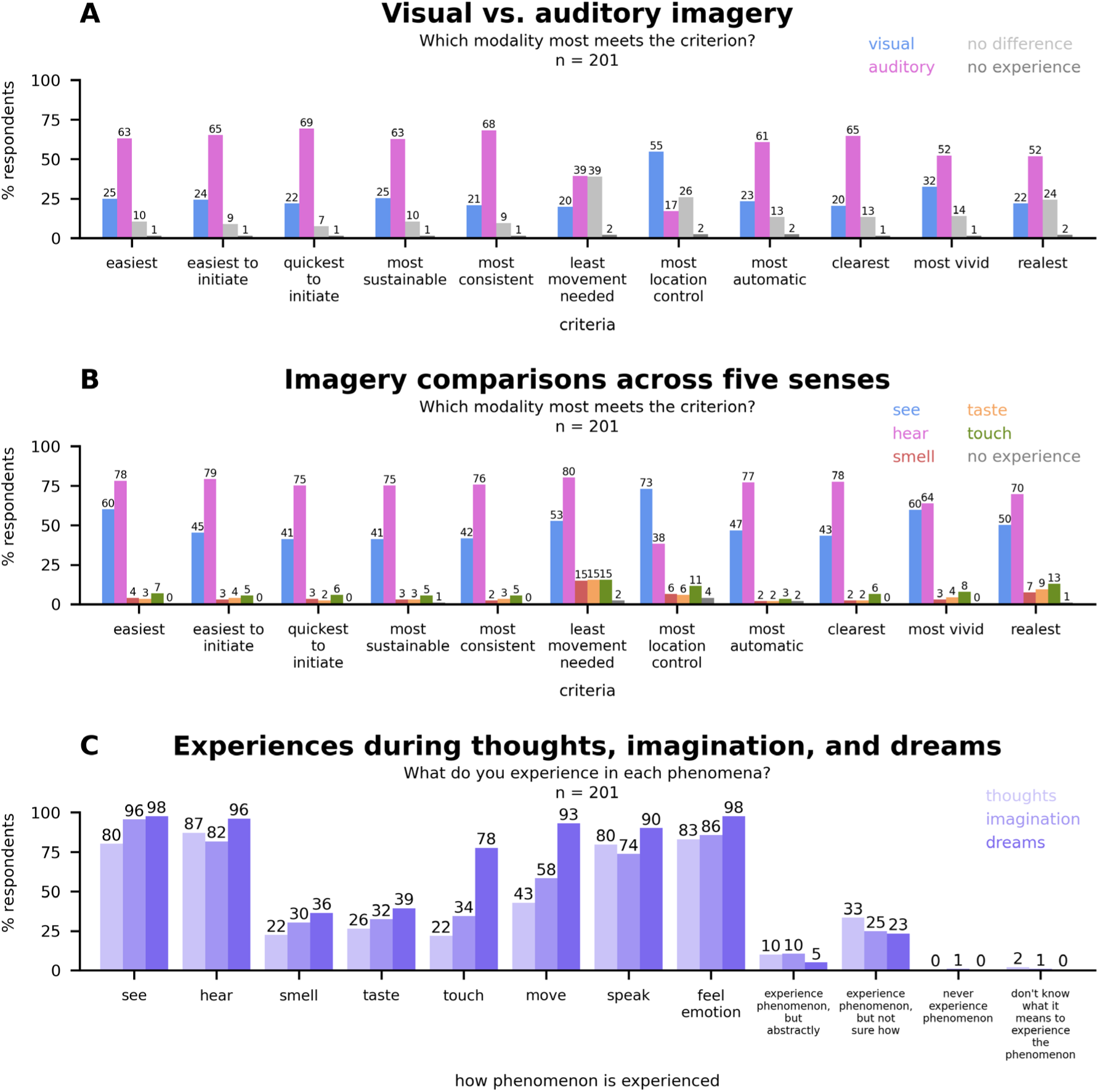
Explicit comparisons of mental imagery modalities, thoughts, imagination, and dreams. *Note.* **A.** Participants made a single choice for each criterion. **B.** Participants could choose multiple responses if each applied equally. **C.** Participants could select each experience that applied in each phenomenon. Statistical comparisons are detailed in the supplementary materials (Supplementary S1.2 & S1.3)

Overall, auditory thought content tended to be most frequently ranked as the best match on each criterion, except for location control. Most participants judged that it was easier to control the apparent location of visual thought content rather than auditory thought content. Although visual imagery was generally less dominant than auditory imagery with respect to most criteria, both of these modalities still tended to eclipse the other sensory modes of thinking for each criterion probed.

### Sensory and non-sensory experiences in thoughts, imagination, and dreams

Given that mental images can be defined as sensory thoughts, it should follow that seeing or hearing one’s thought content is, by definition, mental imagery. However, despite sharing a common root word, “mental imagery” need not necessarily be considered equivalent to, or a part of “imagining”, nor need “thinking” in a sensory way be necessarily considered equivalent to “imagining”. To explore public intuition concerning these phenomena, we asked participants to provide a binary response as to whether they considered imagining the appearance or sound of a letter to be the same process as thinking about the appearance or sound of the same letter. We also probed which forms of sensory and non-sensory experience participants could recall occurring at least sometimes and in some way during their imagination and thoughts (Figure 5C). To compare to another form of internally-generated phenomenon, we also asked participants to specify their experiences during dreaming.

Across 215 respondents, 71.2% reported imagining and thinking to be the same process regarding visual appearance, and 77.2% reported imagining and thinking to be the same regarding sound (Χ²(1) = 4.0, p = .045; Supplementary S1.1, Q9.12 & Q9.13). We also provided an optional free-text field for participants to report on any general differences they perceived between imagining and thinking (i.e. not specifically in reference to thinking or imagining sensory content; Supplementary S1.1, Q9.14). Although many reported no difference even for this general comparison, recurring comments (Supplementary S1.8) regarded that imagination tended to be more associated with sensory content than thinking, and generally more so with visual content. Many also differentiated imagination and thinking as a difference in content, but not process: imagination was frequently regarded as involving more speculative or creative content which was less grounded in immediate reality compared to thinking.

The incidence of particular sensory or non-sensory experiences during thoughts, imagination, and dreams is summarised in Figure 5C. Detailed statistical comparisons within and between phenomena are listed in Supplementary S1.2 & S1.3. Care must be taken to avoid confusing the proportion of individuals reporting a particular experience with the intensity or degree to which individuals believe that experience to occur. Seeing and hearing tend to be the dominant sensory modalities in each experience. Although most individuals consider the three phenomena to involve largely the same experiences, we also observed that there is a small proportion of respondents who experience more sensory, motor, and emotional content moving from thoughts to imagination to dreams, except concerning the auditory experiences of hearing and speaking, which appear slightly more frequently associated with thinking rather than imagining. However, many of these comparisons were not significant (Supplementary S1.3). Interestingly, touch and movement are significantly more frequently reported in dreams relative to other phenomena (Supplementary S1.3). However, our probe prompt for movement, “I feel like I’m moving as I am in my thoughts/imagination/dreams” may not clearly disambiguate between local movements of body parts and global movements of body position.

### Confidence: visual vs. auditory imagery

For each imagery question, participants were asked to indicate how sure they were about their response on a six-point scale from “Completely unsure” (0) to “Completely sure” (5). Here we analysed the confidence of participants when reporting on visual imagery relative to when reporting on auditory imagery for all 34 questions with visual and auditory equivalents. 215 participants were analysed (including non-imagers and those classifying sensory thoughts as hallucinations). For both visual and auditory questions, the modal confidence rating was “Completely sure” and median rating was “Mostly sure” in each modality. To numerically compare confidence between modalities, ratings were converted to a 0-5 score, and the difference in confidence between modalities (visual confidence score - auditory confidence score) was calculated for each of the 34 questions. Hence, there were 7310 possible confidence differences between modalities in total (215 participants X 34 questions). However, as the availability of some questions was conditional on previous answers, some participants did not complete each question for each modality. Hence, only 7068 differences were analysed. A two-tailed Wilcoxon signed-rank test showed that participant confidence was slightly higher when reporting on auditory imagery compared to visual imagery (mean difference = −0.1, standard deviation = 1.1; Z = - 8.9, p < .001). Note that this comparison is only a coarse measure of the confidence difference between modalities. Statistical comparisons of the average paired difference in confidence between modalities for each underlying modality-specific question asked are shown in Supplementary S1.4.

### Confidence: imagination vs. thoughts vs. dreaming

As per visual and auditory imagery questions, we also recorded and analysed the confidence ratings of participants when reporting on what type of sensory or non-sensory experience they had during their thoughts, imagination, and dreams. There were 2580 confidence ratings in total for each phenomenon (215 participants X 12 different types of experience to rate per phenomenon). For thoughts and imagination, the modal rating was “Completely sure” and median rating was “Mostly sure” for each modality. For dreams, the modal and median confidence ratings were both “Completely sure”. Scoring confidence ratings from 0 (Completely unsure) to 5 (Completely sure), we investigated whether confidence was higher in one phenomenon relative to another with two-tailed Wilcoxon signed-rank tests for the 2580 paired differences in confidence between each pair of phenomena. Confidence was significantly higher when reporting on dreams relative to imagination (mean difference = 0.2; Z = −12.2, p < .001, and dreams relative to thoughts (mean difference = 0.3; Z = −14.6, p < .001). Confidence was significantly higher when reporting on imagination relative to thoughts (mean difference = 0.1; Z = −5.2, p < .001). Hence, respondents were most confident about the experiences of dreaming, followed by imagination, then thinking. Mean confidences for each individual aspect of each phenomenon are shown in Supplementary S1.5. Detailed comparisons of confidence within each phenomenon, and between phenomena, are reported in Supplementary S1.6 & S1.7

## Discussion

### Visual vs. auditory mental imagery

Our investigation was able to document how visual and auditory mental imagery are experienced with respect to a vast number of properties. Because we asked participants to imagine the same stimulus in each modality (the letter “O”), and because we probed well-defined stimulus properties (e.g. location, duration, onset latency), we were able to compare how visual and auditory mental images are experienced in a clear and interpretable way. We were also careful to frame our questions with precise and unassuming language, reducing the likelihood of imagery judgements being contaminated by preconceived beliefs about what mental imagery is.

Previous comparisons of mental imagery across modalities have tended to rely on a singular umbrella metric, such as mental image vividness ratings, producing mixed results (e.g. Andrade et al., 2014; Gissurarson, 1992; Schifferstein, 2009; Sheehan, 1967; Switras, 1978; Talamini et al., 2022). By deconvoluting judgements about the experience of imagery into many separate characteristics and properties, our study may be able to provide a more precise, accurate, and interpretable estimate as to how mental imagery differs between visual and auditory modalities, if not other modalities as well. Overall, we found that most individuals experienced audible thought content in a superior manner relative to visual thought content. However, imagery was generally experienced in a quite diverse way, with large groups of respondents still reporting visual mental imagery as dominant in many cases. Yet, auditory thought content was repeatedly reported as more consistent on multiple measures, was more automatically induced by real perception (e.g. reading words and hearing them in your head), started more quickly, lasted longer, and required less body-movements to carry out. Explicit comparisons between all five traditional sensory modalities also showed that, in addition to these factors, more respondents considered auditory thought content to occur most automatically and was the easiest to conduct and initiate. Audition was even dominant with respect to more ambiguous metrics such as clarity, similarity to real perception, and vividness, albeit by smaller margins.

In contrast, visualising thought content tended to be a more deliberate, sluggish, fleeting, and inconsistent process. However, visual imagery did outperform auditory imagery in terms of spatial control. Respondents were also more varied as to where they reported that their visual imagery seemed to be located, probably relating to the fact that the location of visual thought content was more easily manipulated than the location of auditory thought content. Modality-specific differences in mental imagery therefore largely aligned with differences between the modalities during veridical perception. Vision is generally a more precise tool with respect to spatial information, while the auditory system is tuned towards precision in the temporal domain (Ortega et al., 2014).

It is more difficult to account for the dominance of auditory imagery on metrics which are not clearly related to temporal or spatial properties, such as ease, clarity, and the spontaneity of cross-modal imagery. However, we have previously predicted that auditory imagery should likely be a more compelling mental imagery experience if mental images can utilise sensory information generated from overt or covert motor activity related to imagined content (Sulfaro et al., 2022). Mental imagery is likely to be supported by information from real and simulated muscle movements in both visual (Brandt & Stark, 1997; Fourtassi et al., 2017; Laeng & Teodorescu, 2002) and auditory modalities (Pruitt et al., 2019; Tian et al., 2016; Tian & Poeppel, 2010; Whitford et al., 2017). This may help inform imagined content by anchoring it to precise information from real or simulated eye movements in visual imagery and real or simulated speech in auditory imagery. Yet, while eye movements can provide information about the location of an imagined object or the structure of an imagined scene (Brandt & Stark, 1997; Fourtassi et al., 2017; Laeng & Teodorescu, 2002), they should provide much more ambiguous information about image content (e.g. hue, contrast, spatial frequency). However, muscle movements related to synthesising speech of an imagined sound, either overtly or covertly, have the potential to provide information not only about the exact timing of when sound content should be mentally heard, but also about the content of what should be heard as well. If auditory imagery can piggyback off speech simulations, then imagining speech sounds, or even imagined onomatopoeia of non-speech sounds, could let auditory imagery benefit from the rich sensory information created when simulating speech.

The involvement of modality-specific motor information in mental imagery can also logically account for why auditory imagery of speech (i.e. inner speech, internal voices, or internal monologues) are colloquially referred to as “the voice in your head”. If simulated speech is used to inform imagined sound, then imagined sounds would tend to seem like they were originating from the same location that normal speech originates from: inside the head. Indeed, we found that imagined sounds are almost invariably reported as being located inside the head or at the ears, aligning with the location of where real self-produced speech sounds would be expected to be heard. Note however that our study was limited to allowing participants to place only a single marker on a map to indicate the dominant location of imagined content. Therefore, we could not detect if imagery tended to move through a region of space, nor could we discriminate between imagery that was symmetrically yet peripherally distributed (e.g. heard at both ears) from that which was simply positioned asymmetrically (e.g. heard only in the left ear). Future studies may seek to explore the exact spatial distribution of imagined content with more nuance.

While our current investigation cannot comment on the impact of covert movements during imagery, we did record whether participants believed that they required any overtly-detectable body movements to conduct imagery. While eye movements were, expectedly, the most frequently reported body movement required for visual mental imagery, they were quite unexpectedly also the most common movement reported during auditory mental imagery. However, far fewer respondents reported requiring movements of any kind for auditory imagery overall. This is somewhat surprising given the aforementioned suggestion that speech-related movements could improve auditory mental imagery. To explain this pattern of results, consider the findings of Pruitt et al. (2019) who measured speech-related muscle activity during a musical imagery task. Muscle activity was detected during auditory imagery, yet the researchers found that the accuracy of overt singing negatively correlated with laryngeal muscle activity during musical imagery. If real-world singing accuracy were to correlate with imagined singing accuracy, this would imply that motor processes could be recruited as a crutch to support poor imagery. Applying this idea across modalities, as visual imagery was generally poorer than auditory imagery for most respondents in our study, it would be unsurprising that visual imagery also required physical movements for more respondents. Indeed, allowing participants to make eye movements during visual imagery improves the recall of imagined content (Laeng & Teodorescu, 2002).

### The prevalence of mental imagery

While previous work notes that 1-4% of individuals report experiencing poor or absent visual mental imagery (Keogh et al., 2021) our findings show that the incidence of imagery is far from easy to measure consistently. Almost a third of participants reported that they did not experience auditory imagery at least once on the questionnaire and more than half of participants reported that they did not experience visual imagery at least once. Given the major problems inherent to measuring individual differences in the occurrence of mental imagery using self-report measures (Dean & Morris, 2003; Richardson, 1988), we do not consider these reports (nor any currently known reports) to be a firm indicator of the true rate of imagery in the population. Further, mental imagery may encompass a spectrum of experiences with no clear threshold demarcating its presence or absence. However, we do take our findings to indicate that many people have quite an inconsistent moment-to-moment experience of imagery even if they identify as someone who may experience imagery generally.

### Eyes-open vs. eyes-closed mental imagery

Eye-state had a marked impact on reports of visual, but not auditory, mental imagery within our study. Opening one’s eyes caused visual imagery to be less likely to be reported as occurring at all, which we have previously speculated could relate to interference between real and imagined images reducing the signal-to-noise ratio of imagined images (Sulfaro et al., 2022). Opening one’s eyes also prompted many respondents to project the image they were thinking of into the environment rather than forcing it to be restrained close to their eyes or in their head. It is plausible that many respondents tend to project their mental image by default but can only do so onto the back of their eyelids when their eyes are closed, although future studies could explore the exact nature of this difference. Unfortunately, we did not test the effect of an auditory equivalent to eye-closure. Subsequent questionnaires could ask participants to answer questions with or without concurrent noise, or while blocking their ears.

### Modality-specific biases in questioning and terminology

Previous studies tend to investigate the properties of mental imagery either using modality-specific measures (as documented in Alderson-Day & Fernyhough, 2015; Hubbard, 2010; Pearson et al., 2013) or by comparing mental imagery across modalities using ambiguously-defined, yet modality-neutral, metrics such as mental image vividness ratings (e.g. Andrade et al., 2014; Gissurarson, 1992; Schifferstein, 2009; Sheehan, 1967; Switras, 1978; Talamini et al., 2022). However, we specifically tried to create a questionnaire which could investigate clearly-defined aspects of mental imagery in a way that generalised across modalities. Although we believe that we have largely achieved this goal, at least for visual and auditory mental imagery, biases favouring particular modalities may still persist within our methodology. Here, we discuss such considerations, both within our own research as well as mental imagery research generally.

Perhaps the most apparent modality-specific bias present within our study is that our questionnaire was delivered using text such that participants frequently saw the stimulus they were asked to imagine, but never heard it. This could perhaps facilitate visual mental imagery given that the imagined stimulus was usually recently seen, although it could also inhibit imagination by causing adaptation to the stimulus. Some may also argue that our study includes more questions with a temporal focus, as opposed to a spatial focus, favouring auditory imagery. However, we note that we neglected to include temporal analogues of some spatial questions, such as if participants could control how quickly imagery begins, or control for how long it persists, so an intrinsic temporal bias favouring auditory imagery may not be present. It could also be argued that order effects drove a difference between modalities given that participants responded here concerning visual imagery before auditory imagery, however this may not have made a meaningful difference given that auditory imagery still tended to dominate comparisons even when participants were asked to make an explicit judgement about which modality was superior with respect to some criterion.

Criticisms may also be targeted towards our modality-neutral choice of stimulus, the letter “O”. The most obvious critique is its lack of neutrality towards more than two sensory modalities. This meant that we lacked a particular stimulus to imagine during comparisons of the five senses such that respondents were only able to reflect on their imagery experiences generally during such questions. Our disproportionate focus on visual and auditory imagery throughout the rest of our questionnaire may have also biased participant responses towards these modalities during comparisons of the five senses. Ideally, future studies can conceive of an equally-simple stimulus which is effectively matched across even more modalities, but this may be quite a feat. Even here, our stimulus choice may be contentious, and some of our participants reported that they would usually be able to visualise or make audible real objects but had difficulty with our stimulus specifically (Supplementary S1.8).

Additionally, the letter “O” could be interpreted as a letter, simple shape, tone, or phoneme. Future studies may wish to investigate how comparisons between visual and auditory mental imagery may be affected by these different manifestations. For instance, imagining the stimulus as a phoneme could potentially induce the involvement of articulatory motor processes related to synthesising speech sounds. As previously discussed, this may provide additional support to auditory mental images such that imagined speech may be more likely to dominate over visual mental images relative to non-speech auditory imagery. However, most of our respondents reported requiring eye movements for visual mental imagery, while few reported requiring movements of any kind for auditory mental imagery. This suggests that motor processes (or at least overt movements) may be more involved in visual, not auditory, mental imagery. Ultimately, it is unclear whether mental images can be entirely and unambiguously dissociated from motor activity related to the sensory content of imagery, and we have speculated that imagining all sounds as speech sounds may even be the default manner in which auditory imagery occurs given the potential benefits this may entail (Sulfaro et al., 2022).

We also sought to assess whether modality-specific biases exist in the terminology researchers use to investigate mental imagery. Our questionnaire asked participants about how their thought content was experienced in a sensory way rather than explicitly asking participants about their “mental images”. We previously detailed our rationale behind this approach in the Methods section, with one benefit being that such an approach may avoid the preconceived association of the term “mental imagery” with the phenomenon of visual mental imagery specifically. Here, we found that the term “mental imagery” was indeed more often associated with visual rather than auditory imagery, supporting our approach. We also confirmed that most participants understood the act of seeing or hearing thought content to be a form of mental imagery even if many specifically avoided labelling such a process as “mental imagery” in the auditory modality. This indicates both that participants successfully understood our thought-based terminology while simultaneously demonstrating that they did not consider “mental imagery” to be an appropriate descriptor of their audible thought experience, further validating the use of thought-based questioning styles. We did note that some individuals understood the experience of hearing their thought content to refer to a “hallucination” rather than an experience more commonly understood to be mental imagery. However, by surveying participants on how they interpreted our terminology, we were able to filter these respondents accordingly. Overall, while subjectivity inevitably exists in the language used to probe mental imagery, we believe that probing participants using thought-based terminology results in a net bias reduction relative to imagery-based terminology. Given that thought-based terminology also relies on fewer assumptions generally (as previously discussed), future studies should strongly consider adopting a similar approach to that used here, or else be cautious when using “mental imagery” terminology in studies of mental imagery across modalities.

### Novel insights and outstanding questions

Some outstanding questions relate to why vision and audition seem to be far more frequently reported as the most compelling modalities for mental imagery relative to taste, smell, and touch. Of further note is that the senses of taste and smell were infrequently reported as occurring during thoughts, imagination, and dreaming, yet the sense of touch was uniquely reported by the majority of respondents during dreams. It is unclear why this may be the case, although it is possible that respondents may remember seeing themselves physically interacting with the environment during their dreams and therefore conclude that they must have experienced tactile sensations as part of those interactions. The same logic could be applied to explaining the relatively high incidence of movement during dreams relative to imagination and thoughts. The memorability of a particular sensory experience could also be a major factor determining whether that sensory modality is reported to occur during dreams, but not thoughts and imagination, given that it may be difficult to inspect the types of sensory content present in a dream while one is inside a dream.

We also note that internal experiences of speaking may be convolved with experiences of hearing. As one respondent put it, *“I feel like I’m speaking, moreso than hearing. But you can hear yourself speak and that is what I ‘hear’ when thinking.. I think”*. It may be valuable for future studies to try and disambiguate whether participants experience imagined speech sounds as auditory and motor (i.e. proprioceptive) mental imagery simultaneously, or only as auditory or motor imagery individually.

We also noticed commonalities in open-text responses from participants (Supplementary S1.8). Multiple respondents reported incorporating motion or change into their mental images, such as by “tracing” the letter “O” in their mind or having it “flash” on or off. Some of these respondents specified that such dynamics made it easier to visualise their thought content or allowed their thought content to be visualised for longer. It could be that movement reduces the likelihood of adaptation occurring for imagined visual content in any particular region of the visual field, allowing the net period of time over which some form of imagery occurs to be extended. Additionally, if eye-movements are recruited as part of mobilising a mental image, then real or simulated motor movements could be used to anchor mental images to a stream of temporally-precise information. With respect to visual mental imagery, some participants also noted that while they typically considered themselves people who experienced mental imagery, mental images were more likely to occur when they were not trying to actively construct them, in contrast to what was asked of them during our study. While this makes sense given the association between mental imagery, daydreaming, and mind-wandering, it encourages questions as to how neural sensory representations recruited during attention might be distinct from those of unintentional mental imagery.

Finally, despite generally high levels of surety reported across the questionnaire, several respondents touched upon the extreme difficulty of describing their experience. One respondent noted that *“I can’t pinpoint whether I am actually seeing the object or if in my head I just know what it is and am thinking about it … I know what it looks like and I am imagining it in my head but am I actually seeing it?”* Likewise, another commented that *“It’s hard to tell whether I’m actually visualising the ‘O’, or if I’m just imagining myself visualising the ‘O’, with the ‘O’ stored more conceptually than visually.”* Apart from emphasising how difficult it is to quantify the manner in which mental imagery is experienced, these descriptions are indicative of a peculiarity of mental imagery which we have previously predicted elsewhere: individuals should still *feel* like they are conducting mental imagery even if they do not actually *see* their mental images (Sulfaro et al., 2022).

This phenomenon can be explained by considering mental imagery as a retrohierarchical perceptual process (Breedlove et al., 2020; Dentico et al., 2014; Dijkstra et al., 2020; Linde-Domingo et al., 2019). If mental imagery involves propagating a thought from high-level associative cortices towards low-level primary sensory cortices, then thoughts should always at least involve an abstract or semantic experience regardless of how many or how few sensory properties that thought may gain. Indeed, associative cortices, not primary sensory cortices, seem to be most consistently involved in mental imagery (Spagna et al., 2021). Hence, even if someone’s thinking experience never involves any significant sensory experience (that is, a true mental *image*), respondents may still seem to *feel* or *know* that they are imagining what they are thinking about because they are activating high-level neural representations usually involved in the recognition of sensory content. Such an experience may approximate what Watkins described as “invisible imagery”, where one has *“a sensation of having an image that one feels to be there but one can’t see”* (Watkins, 2018, p. 44). Given that mental imagery is virtually unaffected by low-level damage, such as in cortical blindness (Bridge et al., 2012; Chatterjee & Southwood, 1995; de Gelder et al., 2015; Zago et al., 2010), or by mid-level damage, such as in visual agnosia (Behrmann et al., 1994), it may even be that normal imagery is a largely high-to-mid level phenomena with little, if any, actual image content being experienced at all (Sulfaro et al., 2022).

This presents an extra layer of complexity for mental imagery research. Discriminating between truly sensory imagery and merely an abstract impression of imagery using self-report may require considerable introspection from respondents. This may be a high expectation for naïve participants given that the experience of mental imagery is still largely inexplicable even to many researchers in the field, including ourselves. Using rigorous theoretical frameworks and metrics anchored to externally-verifiably standards may be necessary to guide empirical studies around these limitations, and we must take care to avoid our own personal experiences driving our beliefs about the norms of such matters.

## Conclusion

Interpretable empirical findings are essential for informing theoretical explanations of the unique phenomenology of mental imagery. Overall, we show that self-report can be used to make interpretable comparisons of the subjective experience of mental imagery in different sensory modalities. Although we observed a general tendency for imagined sounds to be experienced in a more compelling way than for other imagined content with respect to most metrics tested, we note that participant responses were generally quite diverse. Yet, our findings match theoretically-grounded predictions about how visual and auditory mental imagery should be distinct. Our dataset, analysis code, and questionnaire have all been made available and we encourage the use of our data for answering future research questions which we did not explicitly test here. However, we caution readers to consider the numerous aforementioned biases and challenges plaguing mental imagery research when interpreting current, past, and future studies in the field.

## Data availability

The questionnaire used to conduct this survey, along with additional findings, are provided in Supplementary S1. The full dataset used in our analyses, including data from respondents which we excluded, is provided in Supplementary S2. The code used to conduct our analyses, and to dynamically generate the Results section text, is provided as a Jupyter Source File, Supplementary S3. Additional files, Supplementary S4-7, are images used during the collection and analysis of responses regarding the precise spatial position of imagined content. These images are also used by the code in Supplementary S3.

## Acknowledgements

Alexander is supported by an Australian Government Research Training Program Scholarship. Amanda is supported by an Australian Research Council Discovery Early Career Researcher Award (DE200101159). Thomas is supported by an Australian Research Council Discovery Project (DP200101787).

## References

Alderson-Day, B., & Fernyhough, C. (2015). Inner Speech: Development, Cognitive Functions, Phenomenology, and Neurobiology. Psychological Bulletin, 141(5), 931– 965. https://doi.org/10.1037/bul0000021

Andrade, J., May, J., Deeprose, C., Baugh, S.-J., & Ganis, G. (2014). Assessing vividness of mental imagery: The Plymouth Sensory Imagery Questionnaire. British Journal of Psychology, 105(4), 547–563. https://doi.org/10.1111/bjop.12050

Behrmann, M., Moscovitch, M., & Winocur, G. (1994). Intact visual imagery and impaired visual perception in a patient with visual agnosia. Journal of Experimental Psychology: Human Perception and Performance, 20, 1068–1087. https://doi.org/10.1037/0096-1523.20.5.1068

Brandt, S. A., & Stark, L. W. (1997). Spontaneous eye movements during visual imagery reflect the content of the visual scene. Journal of Cognitive Neuroscience, 9(1), 27–38. https://doi.org/10.1162/jocn.1997.9.1.27

Breedlove, J. L., St-Yves, G., Olman, C. A., & Naselaris, T. (2020). Generative Feedback Explains Distinct Brain Activity Codes for Seen and Mental Images. Current Biology. https://doi.org/10.1016/j.cub.2020.04.014

Bridge, H., Harrold, S., Holmes, E. A., Stokes, M., & Kennard, C. (2012). Vivid visual mental imagery in the absence of the primary visual cortex. Journal of Neurology, 259(6), 1062–1070. https://doi.org/10.1007/s00415-011-6299-z

Chatterjee, A., & Southwood, M. H. (1995). Cortical blindness and visual imagery. Neurology, 45(12), 2189–2195. https://doi.org/10.1212/WNL.45.12.2189

de Gelder, B., Tamietto, M., Pegna, A. J., & Van den Stock, J. (2015). Visual imagery influences brain responses to visual stimulation in bilateral cortical blindness. Cortex, 72, 15–26. https://doi.org/10.1016/j.cortex.2014.11.009

Dean, G. M., & Morris, P. E. (2003). The relationship between self-reports of imagery and spatial ability. British Journal of Psychology, 94(2), 245–273. https://doi.org/10.1348/000712603321661912

Dentico, D., Cheung, B. L., Chang, J.-Y., Guokas, J., Boly, M., Tononi, G., & Van Veen, B. (2014). Reversal of cortical information flow during visual imagery as compared to visual perception. NeuroImage, 100, 237–243. https://doi.org/10.1016/j.neuroimage.2014.05.081

Dijkstra, N., Ambrogioni, L., Vidaurre, D., & van Gerven, M. (2020). Neural dynamics of perceptual inference and its reversal during imagery. ELife, 9, e53588. https://doi.org/10.7554/eLife.53588

Fourtassi, M., Rode, G., & Pisella, L. (2017). Using eye movements to explore mental representations of space. Annals of Physical and Rehabilitation Medicine, 60(3), 160–163. https://doi.org/10.1016/j.rehab.2016.03.001

Galton, F. (1880). Statistics of Mental Imagery. Mind, 5(19), 301–318.

Gissurarson, L. R. (1992). Reported auditory imagery and its relationship with visual imagery. Journal of Mental Imagery, 16(3–4), 117–122.

Hubbard, T. L. (2010). Auditory imagery: Empirical findings. Psychological Bulletin, 136(2), 302–329. https://doi.org/10.1037/a0018436

Keogh, R., Pearson, J., & Zeman, A. (2021). Chapter 15 - Aphantasia: The science of visual imagery extremes. In J. J. S. Barton & A. Leff (Eds.), Handbook of Clinical Neurology (Vol. 178, pp. 277–296). Elsevier. https://doi.org/10.1016/B978-0-12-821377-3.00012-X

Laeng, B., & Teodorescu, D.-S. (2002). Eye scanpaths during visual imagery reenact those of perception of the same visual scene. Cognitive Science, 26(2), 207–231. https://doi.org/10.1207/s15516709cog2602_3

Linde-Domingo, J., Treder, M. S., Kerrén, C., & Wimber, M. (2019). Evidence that neural information flow is reversed between object perception and object reconstruction from memory. Nature Communications, 10(1), Article 1. https://doi.org/10.1038/s41467-018-08080-2

Ortega, L., Guzman-Martinez, E., Grabowecky, M., & Suzuki, S. (2014). Audition dominates vision in duration perception irrespective of salience, attention, and temporal discriminability. Attention, Perception & Psychophysics, 76(5), 1485–1502. https://doi.org/10.3758/s13414-014-0663-x

Pearson, D. G., Deeprose, C., Wallace-Hadrill, S. M. A., Heyes, S. B., & Holmes, E. A. (2013). Assessing mental imagery in clinical psychology: A review of imagery measures and a guiding framework. Clinical Psychology Review, 33(1), 1–23. https://doi.org/10.1016/j.cpr.2012.09.001

Pruitt, T. A., Halpern, A. R., & Pfordresher, P. Q. (2019). Covert singing in anticipatory auditory imagery. Psychophysiology, 56(3), e13297. https://doi.org/10.1111/psyp.13297

Richardson, J. E. (1988). Vividness and unvividness: Reliability, consistency, and validity of subjective imagery ratings. Journal of Mental Imagery, 12(3–4), 115–122.

Schifferstein, H. N. J. (2009). Comparing Mental Imagery across the Sensory Modalities. Imagination, Cognition and Personality, 28(4), 371–388. https://doi.org/10.2190/IC.28.4.g

Sheehan, P. W. (1967). A shortened form of Betts’ questionnaire upon mental imagery. Journal of Clinical Psychology, 23(3), 386–389. https://doi.org/10.1002/1097-4679(196707)23:3<386::AID-JCLP2270230328>3.0.CO;2-S

Spagna, A., Hajhajate, D., Liu, J., & Bartolomeo, P. (2021). Visual mental imagery engages the left fusiform gyrus, but not the early visual cortex: A meta-analysis of neuroimaging evidence. Neuroscience & Biobehavioral Reviews, 122, 201–217. https://doi.org/10.1016/j.neubiorev.2020.12.029

Sulfaro, A. A., Robinson, A. K., & Carlson, T. A. (2022). *Perception as a hierarchical competition: A model that differentiates imagined, veridical, and hallucinated percepts* (p. 2022.09.02.506121). bioRxiv. https://doi.org/10.1101/2022.09.02.506121

Switras, J. E. (1978). An Alternate-Form Instrument to Assess Vividness and Controllability of Mental Imagery in Seven Modalities. Perceptual and Motor Skills, 46(2), 379–384. https://doi.org/10.2466/pms.1978.46.2.379

Talamini, F., Vigl, J., Doerr, E., Grassi, M., & Carretti, B. (2022). Auditory and visual mental imagery in musicians and non-musicians. Musicae Scientiae, 10298649211062724. https://doi.org/10.1177/10298649211062724

Tian, X., & Poeppel, D. (2010). Mental imagery of speech and movement implicates the dynamics of internal forward models. Frontiers in Psychology, 1. https://www.frontiersin.org/articles/10.3389/fpsyg.2010.00166

Tian, X., Zarate, J. M., & Poeppel, D. (2016). Mental imagery of speech implicates two mechanisms of perceptual reactivation. Cortex, 77, 1–12. https://doi.org/10.1016/j.cortex.2016.01.002

Watkins, N. W. (2018). (A)phantasia and severely deficient autobiographical memory: Scientific and personal perspectives. Cortex, 105, 41–52. https://doi.org/10.1016/j.cortex.2017.10.010

Whitford, T. J., Jack, B. N., Pearson, D., Griffiths, O., Luque, D., Harris, A. W., Spencer, K. M., & Le Pelley, M. E. (2017). Neurophysiological evidence of efference copies to inner speech. ELife, 6, e28197. https://doi.org/10.7554/eLife.28197

Zago, S., Corti, S., Bersano, A., Baron, P., Conti, G., Ballabio, E., Lanfranconi, S., Cinnante, C., Costa, A., Cappellari, A., & Bresolin, N. (2010). A Cortically Blind Patient With Preserved Visual Imagery. Cognitive and Behavioral Neurology, 23(1), 44–48. https://doi.org/10.1097/WNN.0b013e3181bf2e6e

