## Supplementary S1-7 for "Comparing mental imagery experiences across visual, auditory, and other sensory modalities": Supplementary S1.docx

**Supplementary 1.1**

*Questionnaire*

Notes on the questionnaire:

1. All questions presented to participants required a mandatory response, unless specified
2. [Square brackets] indicate researcher notes or description
3. The analysis code provided (Supplementary S3) shows which questions are used for each analysis and section of the Results

| **Section** | **Question ID** | **Question/Instruction** |
| --- | --- | --- |
| Preface | Q1.1 | Before you begin this study, make sure you're in a quiet place and using a computer (rather than a mobile device). |
|  | Q2.1^[1]^ | What service linked you to this questionnaire? If you are a USYD student here to earn research participation credits, select "SONA (student)”   - SONA (student) - SONA (paid) - MTurk - Prolific - Other |
|  | Q2.2, Q2.3, Q2.4 | [Participant information sheet] |
|  | Q2.5 | [Participant consent acknowledgement] |
|  | Q3.1 | This questionnaire will ask you to think about specific images or sounds and then report on how you experienced your thoughts. Try to respond honestly based on your actual experience in the moment rather than based on what your experiences prior to this study have led you to believe. |
|  | Q3.2^[2]^ | To check that you're paying attention, some questions may ask for one specific answer. Answering these questions correctly is necessary to ensure your credit for participating in the study is not negatively affected. |
|  | Q3.3 | Note that if you no longer wish to participate, you can close the study window at any point. If you have any queries, feel free to email Alex at.  Your participation today is much appreciated!  Make sure you're in a quiet place, then proceed to the next page when you're ready to begin. |
| Visual imagery | Q4.1 | **Close your eyes and think about the *appearance* of the letter 'O', then respond to the following questions regarding your experience.** |
|  | Q4.2 | Did your thought involve a visual experience? That is, you felt like you could in some way see what you were thinking about, even though your eyes were closed.   - Yes - No |
|  | Q4.3^[3]^ | How sure are you about the accuracy of your answer to the previous question?   - Completely unsure - Mostly unsure - Somewhat unsure - Somewhat sure - Mostly sure - Completely sure |
|  | Q4.4 | **Close your eyes and think about the *appearance* of the letter 'O', *continuing for at least 5 seconds*, then respond to the following questions regarding your experience.** |
|  | Q4.5 | Did your thought automatically seem to involve seeing what you were thinking about in some way?   - No, my thought content was not visible in any way - Yes, automatically, even if I wasn't trying to make it do so - Yes, but only because I specifically tried to make it do so |
|  | Q4.6 | How sure are you about the accuracy of your answer to the previous question?   - Completely unsure - Mostly unsure - Somewhat unsure - Somewhat sure - Mostly sure - Completely sure |
|  | Q4.7 | When did your thought content first seem to become visible to you in some way?   - My thought content was never visible in any way - As soon as I started thinking - Within a second or two after I started thinking - Several or more seconds after I started thinking |
|  | Q4.8 | How sure are you about the accuracy of your answer to the previous question?   - Completely unsure - Mostly unsure - Somewhat unsure - Somewhat sure - Mostly sure - Completely sure |
|  | Q4.9 | Was your thought content visible in some way consistently during the period you were thinking about it?   - My thought content was not visible in any way - My thought content fluctuated in how visible it was during the period - My thought content was consistently visible for the entire period |
|  | Q4.10 | How sure are you about the accuracy of your answer to the previous question?   - Completely unsure - Mostly unsure - Somewhat unsure - Somewhat sure - Mostly sure - Completely sure |
|  | Q4.11 | What proportion of your thinking period involved seeing your thought content in some way?   - My thought content was not visible in any way - Less than half, but not none - About half the time - More than half, but not all All of the period |
|  | Q4.12 | How sure are you about the accuracy of your answer to the previous question?   - Completely unsure - Mostly unsure - Somewhat unsure - Somewhat sure - Mostly sure - Completely sure |
|  | Q4.13 | At what point was your thought content most visible in some way?   - My thought content was not visible in any way - Towards the start of my thought - Towards the middle of my thought - Towards the end of my thought - It was equally visible throughout my thought |
|  | Q4.14 | How sure are you about the accuracy of your answer to the previous question?   - Completely unsure - Mostly unsure - Somewhat unsure - Somewhat sure - Mostly sure - Completely sure |
|  | Q4.15 | **Close your eyes and think about the *appearance* of the letter 'O', then respond to the following questions regarding your experience.** |
|  | Q4.16 | Approximately how many seconds does it take before your thought content becomes visible in some way? *If your duration exceeds the maximum value on the scale, report the maximum value (10)*   - [Slider ranging from 0-10 in increments of 0.1 to indicate Duration (in seconds)] - My thought content was never visible in any way |
|  | Q4.17 | How sure are you about the accuracy of your answer to the previous question?   - Completely unsure - Mostly unsure - Somewhat unsure - Somewhat sure - Mostly sure - Completely sure |
|  | Q4.18 | Approximately how many seconds can you **definitely and continuously** keep your thought content visible for in some way? *If your duration exceeds the maximum value on the scale, report the maximum value (10)*   - [Slider ranging from 0-10 in increments of 0.1 to indicate Duration (in seconds)] - My thought content was never visible in any way |
|  | Q4.19 | How sure are you about the accuracy of your answer to the previous question?   - Completely unsure - Mostly unsure - Somewhat unsure - Somewhat sure - Mostly sure - Completely sure |
|  | Q4.20 | **While trying to remain *completely still*, close your eyes and think about the *appearance* of the letter 'O', then respond to the following questions regarding your experience.** |
|  | Q4.21 | Was your thought content visible in some way even if you were remaining completely still (apart from closing your eyes)?   - No, regardless of whether I moved or not - No, I had to move some part of myself in some way, even if by just a very small amount - Yes, even if I was completely still and unmoving |
|  | Q4.22 | How sure are you about the accuracy of your answer to the previous question?   - Completely unsure - Mostly unsure - Somewhat unsure - Somewhat sure - Mostly sure - Completely sure |
|  | Q4.23^[4]^ | Respond to the following statements regarding which parts of your body needed to move (apart from closing your eyes), even by just a very small amount, in order for your thought content to become visible in some way  My thought was visible, but only if I moved my head  My thought was visible, but only if I moved my brow  My thought was visible, but only if I moved my eyelids  My thought was visible, but only if I moved my eyes  My thought was visible, but only if I moved my ears  My thought was visible, but only if I moved my cheeks  My thought was visible, but only if I moved my nose  My thought was visible, but only if I moved my mouth  My thought was visible, but only if I moved my tongue  My thought was visible, but only if I moved my throat  My thought was visible, but only if I moved my arms or hands  My thought was visible, but only if I moved my torso  My thought was visible, but only if I moved my legs or feet  [For each statement, participants answered the two questions below]  Do you agree with the statement?   - Agree - Disagree - I did not see my thought in any way   How sure are you about the accuracy of your answer?   - Completely unsure - Mostly unsure - Somewhat unsure - Somewhat sure - Mostly sure - Completely sure |
|  | Q4.24 | Was it necessary for you to move any other parts of your body not already mentioned in order for you to in some way see your thought content?   - My thought content was not visible in any way - No - Yes (please specify) [Text response] |
|  | Q4.25 | How sure are you about the accuracy of your answer to the previous question?   - Completely unsure - Mostly unsure - Somewhat unsure - Somewhat sure - Mostly sure - Completely sure |
|  | Q4.26^[5]^ | To prove that you're reading these instructions, please select the last option below.   - Never - Rarely - Sometimes - Often - Always |
|  | Q4.27 | **With your eyes open, think about the *appearance* of the letter 'O', then respond to the following questions regarding your experience.** |
|  | Q4.28^[6]^ | If your thought content was visible in some way, could you choose where it seemed to be located when your eyes were **open**?   - No, my thought content was not visible in any way - No, my thought content was visible, but I could not choose where it seemed to be located - Yes, I could choose where it seemed to be located |
|  | Q4.29 | How sure are you about the accuracy of your answer to the previous question?   - Completely unsure - Mostly unsure - Somewhat unsure - Somewhat sure - Mostly sure - Completely sure |
|  | Q4.30 | If your thought content was visible in some way, where did it most seem to be located when your eyes were **open**, regardless of whether you chose to locate it there or not?   - My thought content was not visible in any way - It had no consistent location - In my head - In my body - In the environment around me - Not in my head, body, or the environment, but somewhere else |
|  | Q4.31 | How sure are you about the accuracy of your answer to the previous question?   - Completely unsure - Mostly unsure - Somewhat unsure - Somewhat sure - Mostly sure - Completely sure |
|  | Q4.32^[7]^ | Please indicate on the diagram where in relation to your body (pictured) your visible thought content most seemed to be located when your eyes were **open**, regardless of whether you chose to locate it there or not.  [Respondents placed a single marker on the image in Supplementary S4] |
|  | Q4.33^f^ | How sure are you about the accuracy of your answer to the previous question?   - Completely unsure - Mostly unsure - Somewhat unsure - Somewhat sure - Mostly sure - Completely sure |
|  | Q4.34^f^ | Please indicate on the diagram where in relation to your head (pictured) your visible thought content most seemed to be located when your eyes were **open**, regardless of whether you chose to locate it there or not.  [Respondents placed a single marker on the image in Supplementary S6] |
|  | Q4.35^f^ | How sure are you about the accuracy of your answer to the previous question?   - Completely unsure - Mostly unsure - Somewhat unsure - Somewhat sure - Mostly sure - Completely sure |
|  | Q4.36 | **With your eyes closed, think about the *appearance* of the letter 'O', then respond to the following questions regarding your experience.** |
|  | Q4.37^e^ | If your thought content was visible in some way, could you choose where it seemed to be located when your eyes were **closed**?   - No, my thought content was not visible in any way - No, my thought was visible, but I could not choose where it seemed to be located - Yes, I could choose where it seemed to be located |
|  | Q4.38 | How sure are you about the accuracy of your answer to the previous question?   - Completely unsure - Mostly unsure - Somewhat unsure - Somewhat sure - Mostly sure - Completely sure |
|  | Q4.39 | If your thought content was visible in some way, where did it most seem to be located when your eyes were **closed**, regardless of whether you chose to locate it there or not?   - My thought content was not visible in any way - It had no consistent location - In my head - In my body - In the environment around me - Not in my head, body, or the environment, but somewhere else |
|  | Q4.40 | How sure are you about the accuracy of your answer to the previous question?   - Completely unsure - Mostly unsure - Somewhat unsure - Somewhat sure - Mostly sure - Completely sure |
|  | Q4.41^[8]^ | Please indicate on the diagram where in relation to your body (pictured) your visible thought content most seemed to be located when your eyes were **closed**, regardless of whether you chose to locate it there or not.  [Respondents placed a single marker on the image in Supplementary S4] |
|  | Q4.42^h^ | How sure are you about the accuracy of your answer to the previous question?   - Completely unsure - Mostly unsure - Somewhat unsure - Somewhat sure - Mostly sure - Completely sure |
|  | Q4.43^h^ | Please indicate on the diagram where in relation to your head (pictured) your visible thought content most seemed to be located when your eyes were **closed**, regardless of whether you chose to locate it there or not.  [Respondents placed a single marker on the image in Supplementary S6] |
|  | Q4.44^h^ | How sure are you about the accuracy of your answer to the previous question?   - Completely unsure - Mostly unsure - Somewhat unsure - Somewhat sure - Mostly sure - Completely sure |
|  | Q4.45 | Is it normal for other people to have thoughts that sometimes seem to have visual properties? That is, do you think that it is normal for other people to feel like they can in some way see what they’re thinking about, even though it’s not there?   - It is normal for other people, and normal for me - It is normal for other people, but not normal for me - It is not normal for other people, but is normal for me - It is not normal for other people, and not normal for me |
|  | Q4.46 | How sure are you about the accuracy of your answer to the previous question?   - Completely unsure - Mostly unsure - Somewhat unsure - Somewhat sure - Mostly sure - Completely sure |
|  | Q4.47 | When thinking about the appearance of something, do you suspect that your thought content is more or less visible compared to the average person?   - More visible - Same as the average person - Less visible |
|  | Q4.48 | How sure are you about the accuracy of your answer to the previous question?   - Completely unsure - Mostly unsure - Somewhat unsure - Somewhat sure - Mostly sure - Completely sure |
|  | Q4.49 | Which of these terms best describes the phenomenon you most commonly have when you seem to in some way see what you're thinking about?   - My thought content is not visible in any way - Hallucination - Mental imagery - Other (please specify) [Text response] |
|  | Q4.50 | How sure are you about the accuracy of your answer to the previous question?   - Completely unsure - Mostly unsure - Somewhat unsure - Somewhat sure - Mostly sure - Completely sure |
|  | Q4.51 | **Close your eyes and think about the appearance of the letter 'O', continuing for at least 5 seconds, then respond to the following questions regarding your experience.** |
|  | Q4.52 | If your thought content was visible in some way, were all parts of it visible at the same time?   - My thought content was not visible in any way - When my thought content was visible, only some parts of it were visible at the same time - When my thought content was visible, all parts of it were visible at the same time |
|  | Q4.53 | How sure are you about the accuracy of your answer to the previous question?   - Completely unsure - Mostly unsure - Somewhat unsure - Somewhat sure - Mostly sure - Completely sure |
|  | Q4.54 | Do you have any extra comments regarding how you experience your thought content in a visual way? *Leave this field blank if not.*   - [Text response] |
| Auditory imagery | Q5.1 | **Close your eyes and think about the *sound* of the letter 'O', then respond to the following questions regarding your experience.** |
|  | Q5.2 | Did your thought involve an auditory experience? That is, you felt like you could in some way hear what you were thinking about, even though you were in a silent place.   - Yes - No |
|  | Q5.3 | How sure are you about the accuracy of your answer to the previous question?   - Completely unsure - Mostly unsure - Somewhat unsure - Somewhat sure - Mostly sure - Completely sure |
|  | Q5.4 | **Close your eyes and think about the *sound* of the letter 'O', *continuing for at least 5 seconds*, then respond to the following questions regarding your experience.** |
|  | Q5.5 | Did your thought automatically seem to involve hearing what you were thinking about in some way?   - No, my thought content was not audible in any way - Yes, automatically, even if I wasn't trying to make it do so - Yes, but only because I specifically tried to make it do so |
|  | Q5.6 | How sure are you about the accuracy of your answer to the previous question?   - Completely unsure - Mostly unsure - Somewhat unsure - Somewhat sure - Mostly sure - Completely sure |
|  | Q5.7 | When did your thought content first seem to become audible to you in some way?   - My thought content was never audible in any way - As soon as I started thinking - Within a second or two after I started thinking - Several or more seconds after I started thinking |
|  | Q5.8 | How sure are you about the accuracy of your answer to the previous question?   - Completely unsure - Mostly unsure - Somewhat unsure - Somewhat sure - Mostly sure - Completely sure |
|  | Q5.9 | Was your thought content audible in some way consistently during the period you were thinking about it?   - My thought content was not audible in any way - My thought content fluctuated in how audible it was during the period - My thought content was consistently audible for the entire period |
|  | Q5.10 | How sure are you about the accuracy of your answer to the previous question?   - Completely unsure - Mostly unsure - Somewhat unsure - Somewhat sure - Mostly sure - Completely sure |
|  | Q5.11 | What proportion of your thinking period involved hearing your thought content in some way?   - My thought content was not audible in any way - Less than half, but not none - About half the time - More than half, but not all - All of the period |
|  | Q5.12 | How sure are you about the accuracy of your answer to the previous question?   - Completely unsure - Mostly unsure - Somewhat unsure - Somewhat sure - Mostly sure - Completely sure |
|  | Q5.13 | At what point was your thought content most audible in some way?   - My thought content was not audible in any way - Towards the start of my thought - Towards the middle of my thought - Towards the end of my thought - It was equally audible throughout my thought |
|  | Q5.14 | How sure are you about the accuracy of your answer to the previous question?   - Completely unsure - Mostly unsure - Somewhat unsure - Somewhat sure - Mostly sure - Completely sure |
|  | Q5.15 | **Close your eyes and think about the *sound* of the letter 'O', then respond to the following questions regarding your experience.** |
|  | Q5.16 | Approximately how many seconds does it take before your thought content becomes audible in some way? *If your duration exceeds the maximum value on the scale, report the maximum value (10)*   - [Slider ranging from 0-10 in increments of 0.1 to indicate Duration (in seconds)] - My thought content was never audible in any way |
|  | Q5.17 | How sure are you about the accuracy of your answer to the previous question?   - Completely unsure - Mostly unsure - Somewhat unsure - Somewhat sure - Mostly sure - Completely sure |
|  | Q5.18 | Approximately how many seconds can you **definitely and continuously** keep your thought content audible for in some way? *If your duration exceeds the maximum value on the scale, report the maximum value (10)* |
|  | Q5.19 | How sure are you about the accuracy of your answer to the previous question?   - Completely unsure - Mostly unsure - Somewhat unsure - Somewhat sure - Mostly sure - Completely sure |
|  | Q5.20 | **While trying to remain *completely still*, close your eyes and think about the *sound* of the letter 'O', then respond to the following questions regarding your experience.** |
|  | Q5.21 | Was your thought content audible in some way even if you were remaining completely still (apart from closing your eyes)?   - No, regardless of whether I moved or not - No, I had to move some part of myself in some way, even if by just a very small amount - Yes, even if I was completely still and unmoving |
|  | Q5.22 | How sure are you about the accuracy of your answer to the previous question?   - Completely unsure - Mostly unsure - Somewhat unsure - Somewhat sure - Mostly sure - Completely sure |
|  | Q5.23^c^ | Respond to the following statements regarding which parts of your body needed to move (apart from closing your eyes), even by just a very small amount, in order for your thought content to become audible in some way  My thought was audible, but only if I moved my head  My thought was audible, but only if I moved my brow  My thought was audible, but only if I moved my eyelids  My thought was audible, but only if I moved my eyes  My thought was audible, but only if I moved my ears  My thought was audible, but only if I moved my cheeks  My thought was audible, but only if I moved my nose  My thought was audible, but only if I moved my mouth  My thought was audible, but only if I moved my tongue  My thought was audible, but only if I moved my throat  My thought was audible, but only if I moved my arms or hands  My thought was audible, but only if I moved my torso  My thought was audible, but only if I moved my legs or feet  [For each statement, participants answered the two questions below]  Do you agree with the statement?   - Agree - Disagree - My thought content was not audible in any way   How sure are you about the accuracy of your answer?   - Completely unsure - Mostly unsure - Somewhat unsure - Somewhat sure - Mostly sure - Completely sure |
|  | Q5.24 | Was it necessary for you to move any other parts of your body not already mentioned in order for you to in some way hear your thought content?   - My thought content was not audible in any way - No - Yes (please specify) [Text response] |
|  | Q5.25 | How sure are you about the accuracy of your answer to the previous question?   - Completely unsure - Mostly unsure - Somewhat unsure - Somewhat sure - Mostly sure - Completely sure |
|  | Q5.26^[9]^ | To show that you're paying attention, select the option below which starts with the first letter of the alphabet.   - Louder than usual - About the same - Quieter than usual |
|  | Q5.27 | **With your eyes open, think about the *sound* of the letter 'O', then respond to the following questions regarding your experience.** |
|  | Q5.28^[10]^ | If your thought content was audible in some way, could you choose where it seemed to be located when your eyes were **open**?   - No, my thought content was not audible in any way - No, my thought content was audible, but I could not choose where it seemed to be located - Yes, I could choose where it seemed to be located |
|  | Q5.29 | How sure are you about the accuracy of your answer to the previous question?   - Completely unsure - Mostly unsure - Somewhat unsure - Somewhat sure - Mostly sure - Completely sure |
|  | Q5.30 | If your thought content was audible in some way, where did it most seem to be located when your eyes were **open**, regardless of whether you chose to locate it there or not?   - My thought content was not audible in any way - It had no consistent location - In my head - In my body - In the environment around me - Not in my head, body, or the environment, but somewhere else |
|  | Q5.31 | How sure are you about the accuracy of your answer to the previous question?   - Completely unsure - Mostly unsure - Somewhat unsure - Somewhat sure - Mostly sure - Completely sure |
|  | Q5.32^[11]^ | Please indicate on the diagram where in relation to your body (pictured) your audible thought content most seemed to be located when your eyes were **open**, regardless of whether you chose to locate it there or not.  [Respondents placed a single marker on the image in Supplementary S4] |
|  | Q5.33^l^ | How sure are you about the accuracy of your answer to the previous question?   - Completely unsure - Mostly unsure - Somewhat unsure - Somewhat sure - Mostly sure - Completely sure |
|  | Q5.34^l^ | Please indicate on the diagram where in relation to your head (pictured) your audible thought content most seemed to be located when your eyes were **open**, regardless of whether you chose to locate it there or not.  [Respondents placed a single marker on the image in Supplementary S6] |
|  | Q5.35^l^ | How sure are you about the accuracy of your answer to the previous question?   - Completely unsure - Mostly unsure - Somewhat unsure - Somewhat sure - Mostly sure - Completely sure |
|  | Q5.36 | **With your eyes closed, think about the *sound* of the letter 'O', then respond to the following questions regarding your experience.** |
|  | Q5.37^i^ | If your thought content was audible in some way, could you choose where it seemed to be located when your eyes were **closed**?   - No, my thought content was not audible in any way - No, my thought was audible, but I could not choose where it seemed to be located - Yes, I could choose where it seemed to be located |
|  | Q5.38 | How sure are you about the accuracy of your answer to the previous question?   - Completely unsure - Mostly unsure - Somewhat unsure - Somewhat sure - Mostly sure - Completely sure |
|  | Q5.39 | If your thought content was audible in some way, where did it most seem to be located when your eyes were **closed**, regardless of whether you chose to locate it there or not?   - My thought content was not audible in any way - It had no consistent location - In my head - In my body - In the environment around me - Not in my head, body, or the environment, but somewhere else |
|  | Q5.40 | How sure are you about the accuracy of your answer to the previous question?   - Completely unsure - Mostly unsure - Somewhat unsure - Somewhat sure - Mostly sure - Completely sure |
|  | Q5.41^[12]^ | Please indicate on the diagram where in relation to your body (pictured) your audible thought content most seemed to be located when your eyes were **closed**, regardless of whether you chose to locate it there or not.  [Respondents placed a single marker on the image in Supplementary S4] |
|  | Q5.42^n^ | How sure are you about the accuracy of your answer to the previous question?   - Completely unsure - Mostly unsure - Somewhat unsure - Somewhat sure - Mostly sure - Completely sure |
|  | Q5.43^n^ | Please indicate on the diagram where in relation to your head (pictured) your audible thought content most seemed to be located when your eyes were **closed**, regardless of whether you chose to locate it there or not.  [Respondents placed a single marker on the image in Supplementary S6] |
|  | Q5.44^n^ | How sure are you about the accuracy of your answer to the previous question?   - Completely unsure - Mostly unsure - Somewhat unsure - Somewhat sure - Mostly sure - Completely sure |
|  | Q5.45 | Is it normal for other people to have thoughts that sometimes seem to have auditory properties? That is, do you think that it is normal for other people to feel like they can in some way hear what they’re thinking about, even though it’s not there?   - It is normal for other people, and normal for me - It is normal for other people, but not normal for me - It is not normal for other people, but is normal for me - It is not normal for other people, and not normal for me |
|  | Q5.46 | How sure are you about the accuracy of your answer to the previous question?   - Completely unsure - Mostly unsure - Somewhat unsure - Somewhat sure - Mostly sure - Completely sure |
|  | Q5.47 | When thinking about the sound of something, do you suspect that your thought content is more or less audible compared to the average person?   - More audible - Same as the average person - Less audible |
|  | Q5.48 | How sure are you about the accuracy of your answer to the previous question?   - Completely unsure - Mostly unsure - Somewhat unsure - Somewhat sure - Mostly sure - Completely sure |
|  | Q5.49 | Which of these terms best describes the phenomenon you most commonly have when you seem to in some way hear what you're thinking about?   - My thought content is not audible in any way - Hallucination - Mental imagery - Other (please specify) [Text response] |
|  | Q5.50 | How sure are you about the accuracy of your answer to the previous question?   - Completely unsure - Mostly unsure - Somewhat unsure - Somewhat sure - Mostly sure - Completely sure |
|  | Q5.51 | Do you have any extra comments regarding how you experience your thought content in an auditory way? *Leave this field blank if not.*   - [Text response] |
| Visuo-audio | Q6.1^[13]^ | When reading this sentence, do you in some way hear at least some of the words you're reading as you see them, even though you're in no way speaking the words aloud?   - Yes, automatically - Yes, but only if I specifically try/tried to - No |
|  | Q6.2 | How sure are you about the accuracy of your answer to the previous question?   - Completely unsure - Mostly unsure - Somewhat unsure - Somewhat sure - Mostly sure - Completely sure |
|  | Q6.3^[14]^ | If someone were to read this sentence aloud to you, do you suspect that you would in some way see at least some of the words that you heard, even if you weren't looking at them?   - Yes, automatically - Yes, but only if I specifically try/tried to - No |
|  | Q6.4 | How sure are you about the accuracy of your answer to the previous question?   - Completely unsure - Mostly unsure - Somewhat unsure - Somewhat sure - Mostly sure - Completely sure |
|  | Q6.5^[15]^ | If you close your eyes and think about the appearance of the letter 'O', do you **also** seem to hear the sound of the letter in some way **in addition to** seeming to see an image of the letter in some way?   - Yes, automatically - Yes, but only if I specifically try/tried to - No, I only see what I'm thinking about in some way - No, I do not see or hear what I'm thinking about in any way |
|  | Q6.6 | How sure are you about the accuracy of your answer to the previous question?   - Completely unsure - Mostly unsure - Somewhat unsure - Somewhat sure - Mostly sure - Completely sure |
|  | Q6.7^[16]^ | If you close your eyes and think about the sound of the letter 'O', do you **also** seem to see an image of the letter in some way **in addition to** seeming to hear the sound of the letter in some way also?   - Yes, automatically - Yes, but only if I specifically try/tried to - No, I only hear what I'm thinking about in some way - No, I do not see or hear what I'm thinking about in any way |
|  | Q6.8 | How sure are you about the accuracy of your answer to the previous question?   - Completely unsure - Mostly unsure - Somewhat unsure - Somewhat sure - Mostly sure - Completely sure |
|  | Q6.9 | Respond to the following questions concerning your general experience when thinking about the appearance or sound of the letter 'O' throughout this questionnaire.  Which is the easiest to do in some way?  Which is the easiest to initiate?  Which takes the least time to initiate?  Which can you definitely and continuously experience for the longest after it's initiated?  Which is experienced with the most consistency over a 5-second period?  Which requires the least body or head movement to do?  Which are you most able to control the apparent spatial location or origin of?  Which occurs most automatically (i.e. without you specifically trying to do it)?  Which is clearest?  Which is most vivid?  Which is most similar to the experience you have when you perceive real things in your environment using the same sense?  [For each statement, participants chose a response from below and then answered a confidence question]  Responses   - Seeing my thought content in some way - Hearing my thought content in some way - There is no difference - I do not see or hear my thought content in any way   How sure are you about your answer?   - Completely unsure - Mostly unsure - Somewhat unsure - Somewhat sure - Mostly sure - Completely sure |
|  | Q6.10 | Do you have any extra comments regarding how you experience your thought content in a visual or auditory way? *Leave this field blank if not.*   - [Text response] |
| Multimodal | Q7.1 | While awake, if you have a **thought**, indicate which of the following phenomena you experience at least sometimes and in some way  I feel like I’m seeing what I’m thinking about  I feel like I’m hearing what I’m thinking about  I feel like I’m touching what I’m thinking about  I feel like I’m smelling what I’m thinking about  I feel like I’m tasting what I’m thinking about  I feel like I’m moving as I am in my thought  I feel like I’m speaking as I am in my thought  I feel the emotion I’m thinking about  I have thoughts, but do not feel like I’m seeing, hearing, touching, smelling, tasting, moving, speaking, or feeling emotions during my thoughts  I have thoughts, but am not sure what I experience during my thoughts  I never have thoughts  I do not know what it means to have thoughts  [For each statement, participants answered the two questions below]  Do you agree with the statement?   - Agree - Disagree   How sure are you about the accuracy of your answer?   - Completely unsure - Mostly unsure - Somewhat unsure - Somewhat sure - Mostly sure - Completely sure |
|  | Q7.2^[17]^ | While awake, if you **imagine** something, indicate which of the following phenomena you experience at least sometimes and in some way  I will agree with this statement with complete surety to prove that I'm paying attention  I feel like I’m seeing what I’m imagining  I feel like I’m hearing what I’m imagining  I feel like I’m touching what I’m imagining  I feel like I’m smelling what I’m imagining  I feel like I’m tasting what I’m imagining  I feel like I’m moving as I am in my imagination  I feel like I’m speaking as I am in my imagination  I feel the emotion I’m imagining  I imagine things, but do not feel like I’m seeing, hearing, touching, smelling, tasting, moving, speaking, or feeling emotions while imagining  I imagine things, but am not sure what I experience while imagining  I never imagine things  I do not know what it means to imagine  [For each statement, participants answered the two questions below]  Do you agree with the statement?   - Agree - Disagree   How sure are you about the accuracy of your answer?   - Completely unsure - Mostly unsure - Somewhat unsure - Somewhat sure - Mostly sure - Completely sure |
|  | Q7.3 | While sleeping, if you have a **dream**, indicate which of the following phenomena you experience at least sometimes and in some way  I feel like I see things in my dreams  I feel like I hear things in my dreams  I feel like I touch things in my dreams  I feel like I smell things in my dreams  I feel like I taste things in my dreams  I feel like I move while in my dreams  I feel like I speak while in my dreams  I feel emotions while in my dreams  I dream, but do not feel like I’m seeing, hearing, touching, smelling, tasting, moving, speaking, or feeling emotions during my dreams  I dream, but am not sure what I experience during my dreams  I never dream  I do not know what it means to dream  [For each statement, participants answered the two questions below]  Do you agree with the statement?   - Agree - Disagree   How sure are you about the accuracy of your answer?   - Completely unsure - Mostly unsure - Somewhat unsure - Somewhat sure - Mostly sure - Completely sure |
|  | Q7.4 | Do you have any extra comments regarding how you experience your thoughts, imaginings, or dreams? *Leave this field blank if not.*   - [Text response] |
| Modality rankings | Q8.1 | **Respond to the following questions regarding how you experience your thoughts *in general*.** |
|  | Q8.2 | Which is the easiest to do in some way? *Multiple options may be selected if they are experienced equivalently.*   - Seeing my thought content in some way - Hearing my thought content in some way - Tasting my thought content in some way - Smelling my thought content in some way - Feeling (touching) my thought content in some way - I do not experience my thought content in any sensory way |
|  | Q8.3 | How sure are you about the accuracy of your answer to the previous question?   - Completely unsure - Mostly unsure - Somewhat unsure - Somewhat sure - Mostly sure - Completely sure |
|  | Q8.4 | Which is the easiest to initiate in some way? *Multiple options may be selected if they are experienced equivalently.*   - Seeing my thought content in some way - Hearing my thought content in some way - Tasting my thought content in some way - Smelling my thought content in some way - Feeling (touching) my thought content in some way - I do not experience my thought content in any sensory way |
|  | Q8.5 | How sure are you about the accuracy of your answer to the previous question?   - Completely unsure - Mostly unsure - Somewhat unsure - Somewhat sure - Mostly sure - Completely sure |
|  | Q8.6 | Which takes the least time to initiate? *Multiple options may be selected if they are experienced equivalently.*   - Seeing my thought content in some way - Hearing my thought content in some way - Tasting my thought content in some way - Smelling my thought content in some way - Feeling (touching) my thought content in some way - I do not experience my thought content in any sensory way |
|  | Q8.7 | How sure are you about the accuracy of your answer to the previous question?   - Completely unsure - Mostly unsure - Somewhat unsure - Somewhat sure - Mostly sure - Completely sure |
|  | Q8.8 | Which can you definitely and continuously experience for the longest after it's initiated? *Multiple options may be selected if they are experienced equivalently.*   - Seeing my thought content in some way - Hearing my thought content in some way - Tasting my thought content in some way - Smelling my thought content in some way - Feeling (touching) my thought content in some way - I do not experience my thought content in any sensory way |
|  | Q8.9 | How sure are you about the accuracy of your answer to the previous question?   - Completely unsure - Mostly unsure - Somewhat unsure - Somewhat sure - Mostly sure - Completely sure |
|  | Q8.10 | Which is experienced with the most consistency over a 5-second period? *Multiple options may be selected if they are experienced equivalently.*   - Seeing my thought content in some way - Hearing my thought content in some way - Tasting my thought content in some way - Smelling my thought content in some way - Feeling (touching) my thought content in some way - I do not experience my thought content in any sensory way - Seeing my thought content in some way - Hearing my thought content in some way - Tasting my thought content in some way - Smelling my thought content in some way - Feeling (touching) my thought content in some way - I do not experience my thought content in any sensory way |
|  | Q8.11 | How sure are you about the accuracy of your answer to the previous question?   - Completely unsure - Mostly unsure - Somewhat unsure - Somewhat sure - Mostly sure - Completely sure |
|  | Q8.12 | Which requires the least body or head movement to do? *Multiple options may be selected if they are experienced equivalently.*   - Seeing my thought content in some way - Hearing my thought content in some way - Tasting my thought content in some way - Smelling my thought content in some way - Feeling (touching) my thought content in some way - I do not experience my thought content in any sensory way |
|  | Q8.13 | How sure are you about the accuracy of your answer to the previous question?   - Completely unsure - Mostly unsure - Somewhat unsure - Somewhat sure - Mostly sure - Completely sure |
|  | Q8.14 | Which are you most able to control the apparent spatial location or origin of? *Multiple options may be selected if they are experienced equivalently.*   - Seeing my thought content in some way - Hearing my thought content in some way - Tasting my thought content in some way - Smelling my thought content in some way - Feeling (touching) my thought content in some way - I do not experience my thought content in any sensory way |
|  | Q8.15 | How sure are you about the accuracy of your answer to the previous question?   - Completely unsure - Mostly unsure - Somewhat unsure - Somewhat sure - Mostly sure - Completely sure |
|  | Q8.16 | Which occurs most automatically (i.e. without you specifically trying to do it)? *Multiple options may be selected if they are experienced equivalently.*   - Seeing my thought content in some way - Hearing my thought content in some way - Tasting my thought content in some way - Smelling my thought content in some way - Feeling (touching) my thought content in some way - I do not experience my thought content in any sensory way |
|  | Q8.17 | How sure are you about the accuracy of your answer to the previous question?   - Completely unsure - Mostly unsure - Somewhat unsure - Somewhat sure - Mostly sure - Completely sure |
|  | Q8.18 | Which is clearest? *Multiple options may be selected if they are experienced equivalently.*   - Seeing my thought content in some way - Hearing my thought content in some way - Tasting my thought content in some way - Smelling my thought content in some way - Feeling (touching) my thought content in some way - I do not experience my thought content in any sensory way |
|  | Q8.19 | How sure are you about the accuracy of your answer to the previous question?   - Completely unsure - Mostly unsure - Somewhat unsure - Somewhat sure - Mostly sure - Completely sure |
|  | Q8.20 | Which is most vivid? *Multiple options may be selected if they are experienced equivalently.*   - Seeing my thought content in some way - Hearing my thought content in some way - Tasting my thought content in some way - Smelling my thought content in some way - Feeling (touching) my thought content in some way - I do not experience my thought content in any sensory way |
|  | Q8.21 | How sure are you about the accuracy of your answer to the previous question?   - Completely unsure - Mostly unsure - Somewhat unsure - Somewhat sure - Mostly sure - Completely sure |
|  | Q8.22 | Which is most similar to the experience you have when you perceive real things in your environment using the same sense? *Multiple options may be selected if they are experienced equivalently.*   - Seeing my thought content in some way - Hearing my thought content in some way - Tasting my thought content in some way - Smelling my thought content in some way - Feeling (touching) my thought content in some way - I do not experience my thought content in any sensory way |
|  | Q8.23 | How sure are you about the accuracy of your answer to the previous question?   - Completely unsure - Mostly unsure - Somewhat unsure - Somewhat sure - Mostly sure - Completely sure |
|  | Q8.24 | Do you have any extra comments regarding how you experience your thoughts in different senses? *Leave this field blank if not.*   - [Text response] |
| Demographic/Other | Q9.1 | What is your age?   - [Numerical response] |
|  | Q9.2 | What is your sex?   - Female - Male - Other - Prefer not to say |
|  | Q9.3 | Do you have any known neurological, psychological, or psychiatric conditions you're aware of which are likely to affect your thoughts, imagination, dreams, or perception?   - Yes - No - Prefer not to say |
|  | Q9.4 | Is English your dominant language?   - Yes - No |
|  | Q9.5 | Is English the language you are most comfortable reading in?   - Yes - No |
|  | Q9.6 | What country did you answer this questionnaire in?   - [Respondents select from a dropdown menu] |
|  | Q9.7 | Which country were you born in?   - [Respondents select from a dropdown menu] |
|  | Q9.8 | Approximately how many hours have passed since you last woke up (including from a nap)? Round your answer to the nearest hour.   - [Numerical response] |
|  | Q9.9 | Which term best describes the noise level of the environment you answered this questionnaire in?   - Quiet (minimal noise) - Moderate (some noise, but it was not distracting) - Noisy or loud (loud and/or distracting noise) |
|  | Q9.10 | Which term best describes the brightness level of the environment you answered this questionnaire in?   - Bright (lighting is from daylight, or approximates daylight) - Dim (lighting is from lamplight, or approximates lamplight) - Dark (lighting is mostly from this device's screen) |
|  | Q9.11 | Please describe your environment in one sentence or less (e.g. bedroom, library, backyard)   - [Text response] |
|  | Q9.12 | Is imagining the appearance of the letter 'O' and thinking about the appearance of the letter 'O' the same process to you?   - Yes - No |
|  | Q9.13 | Is imagining the sound of the letter 'O' and thinking about the sound of the letter 'O' the same process to you?   - Yes - No |
|  | Q9.14 | Do you think there is a difference between thinking about something and imagining something? If so, can you explain what you personally believe the difference is? *This question is optional.*   - [Text response] |
|  | Q9.15 | Do you have any comments regarding this questionnaire or any of your answers?  *Leave this field blank if not.*   - [Text response] |
| Finish | Q10.1 | Thank you for participating in this study! If you have finished your answers and are ready to submit, click the button below and then proceed.  *If you are participating in this study as a SONA student, proceeding will automatically provide you with your research credit.*   - I'm ready to submit |

**Supplementary S1.2**

*Within-phenomenon paired comparisons of the proportion with which each aspect was reported during thoughts, imagination, and dreams*

| **comparison** | **aspect 1** | **aspect 2** | **thoughts** | **imagination** | **dreams** |
| --- | --- | --- | --- | --- | --- |
| 1 | see | hear | p = .059127 | p = .000002 | p = .255321 |
| 2 | see | smell | p < .000001 | p < .000001 | p < .000001 |
| 3 | see | taste | p < .000001 | p < .000001 | p < .000001 |
| 4 | see | touch | p < .000001 | p < .000001 | p < .000001 |
| 5 | see | move | p < .000001 | p < .000001 | p = .011227 |
| 6 | see | speak | p = .805512 | p < .000001 | p = .000168 |
| 7 | see | feel emotion | p = .430082 | p = .000409 | p > .999999 |
| 8 | see | exp. abstract | p < .000001 | p < .000001 | p < .000001 |
| 9 | see | exp. but unsure | p < .000001 | p < .000001 | p < .000001 |
| 10 | see | never exp. | p < .000001 | p < .000001 | p < .000001 |
| 11 | see | don't know | p < .000001 | p < .000001 | p < .000001 |
| 12 | hear | smell | p < .000001 | p < .000001 | p < .000001 |
| 13 | hear | taste | p < .000001 | p < .000001 | p < .000001 |
| 14 | hear | touch | p < .000001 | p < .000001 | p < .000001 |
| 15 | hear | move | p < .000001 | p < .000001 | p = .106555 |
| 16 | hear | speak | p = .010144 | p = .020196 | p = .006287 |
| 17 | hear | feel emotion | p = .226121 | p = .192501 | p = .316110 |
| 18 | hear | exp. abstract | p < .000001 | p < .000001 | p < .000001 |
| 19 | hear | exp. but unsure | p < .000001 | p < .000001 | p < .000001 |
| 20 | hear | never exp. | p < .000001 | p < .000001 | p < .000001 |
| 21 | hear | don't know | p < .000001 | p < .000001 | p < .000001 |
| 22 | smell | taste | p = .192501 | p = .449050 | p = .155239 |
| 23 | smell | touch | p = .875889 | p = .226121 | p < .000001 |
| 24 | smell | move | p = .000002 | p < .000001 | p < .000001 |
| 25 | smell | speak | p < .000001 | p < .000001 | p < .000001 |
| 26 | smell | feel emotion | p < .000001 | p < .000001 | p < .000001 |
| 27 | smell | exp. abstract | p = .000662 | p = .000001 | p < .000001 |
| 28 | smell | exp. but unsure | p = .014876 | p = .242025 | p = .005756 |
| 29 | smell | never exp. | p < .000001 | p < .000001 | p < .000001 |
| 30 | smell | don't know | p < .000001 | p < .000001 | p < .000001 |
| 31 | taste | touch | p = .157806 | p = .526680 | p < .000001 |
| 32 | taste | move | p = .000120 | p < .000001 | p < .000001 |
| 33 | taste | speak | p < .000001 | p < .000001 | p < .000001 |
| 34 | taste | feel emotion | p < .000001 | p < .000001 | p < .000001 |
| 35 | taste | exp. abstract | p = .000019 | p < .000001 | p < .000001 |
| 36 | taste | exp. but unsure | p = .124434 | p = .125562 | p = .000791 |
| 37 | taste | never exp. | p < .000001 | p < .000001 | p < .000001 |
| 38 | taste | don't know | p < .000001 | p < .000001 | p < .000001 |
| 39 | touch | move | p < .000001 | p < .000001 | p < .000001 |
| 40 | touch | speak | p < .000001 | p < .000001 | p = .000069 |
| 41 | touch | feel emotion | p < .000001 | p < .000001 | p < .000001 |
| 42 | touch | exp. abstract | p = .001805 | p < .000001 | p < .000001 |
| 43 | touch | exp. but unsure | p = .013336 | p = .051513 | p < .000001 |
| 44 | touch | never exp. | p < .000001 | p < .000001 | p < .000001 |
| 45 | touch | don't know | p < .000001 | p < .000001 | p < .000001 |
| 46 | move | speak | p < .000001 | p = .000036 | p = .131446 |
| 47 | move | feel emotion | p < .000001 | p < .000001 | p = .011227 |
| 48 | move | exp. abstract | p < .000001 | p < .000001 | p < .000001 |
| 49 | move | exp. but unsure | p = .051513 | p < .000001 | p < .000001 |
| 50 | move | never exp. | p < .000001 | p < .000001 | p < .000001 |
| 51 | move | don't know | p < .000001 | p < .000001 | p < .000001 |
| 52 | speak | feel emotion | p = .300420 | p = .000520 | p = .000168 |
| 53 | speak | exp. abstract | p < .000001 | p < .000001 | p < .000001 |
| 54 | speak | exp. but unsure | p < .000001 | p < .000001 | p < .000001 |
| 55 | speak | never exp. | p < .000001 | p < .000001 | p < .000001 |
| 56 | speak | don't know | p < .000001 | p < .000001 | p < .000001 |
| 57 | feel emotion | exp. abstract | p < .000001 | p < .000001 | p < .000001 |
| 58 | feel emotion | exp. but unsure | p < .000001 | p < .000001 | p < .000001 |
| 59 | feel emotion | never exp. | p < .000001 | p < .000001 | p < .000001 |
| 60 | feel emotion | don't know | p < .000001 | p < .000001 | p < .000001 |
| 61 | exp. abstract | exp. but unsure | p < .000001 | p = .000003 | p < .000001 |
| 62 | exp. abstract | never exp. | p = .000015 | p = .000015 | p = .005702 |
| 63 | exp. abstract | don't know | p = .000218 | p = .000005 | p = .005702 |
| 64 | exp. but unsure | never exp. | p < .000001 | p < .000001 | p < .000001 |
| 65 | exp. but unsure | don't know | p < .000001 | p < .000001 | p < .000001 |
| 66 | never exp. | don't know | p = .177768 | p > .999999 | p > .999999 |

*Note.* Outcomes for Bhapkar (McNemar) homogeneity tests for all possible comparisons within each phenomenon (i.e. within either thoughts, imagination, or dreams). Significant comparisons before correction (alpha = 0.05) are highlighted in orange. Significant comparisons after Bonferroni correction (alpha = 0.05/234) are highlighted in red.

**Supplementary S1.3**

*Between-phenomenon paired comparisons of the proportion with which each aspect was reported during thoughts, imagination, and dreams*

| **comparison** | **aspect** | **thoughts vs. imagination** | **thoughts vs. dreams** | **imagination vs. dreams** |
| --- | --- | --- | --- | --- |
| 1 | see | p < .000001 | p < .000001 | p = .204102 |
| 2 | hear | p = .046040 | p = .000730 | p < .000001 |
| 3 | smell | p = .003900 | p = .000574 | p = .155239 |
| 4 | taste | p = .043373 | p = .001980 | p = .101371 |
| 5 | touch | p = .000010 | p < .000001 | p < .000001 |
| 6 | move | p = .000003 | p < .000001 | p < .000001 |
| 7 | speak | p = .055556 | p = .000648 | p = .000004 |
| 8 | feel emotion | p = .249801 | p < .000001 | p = .000002 |
| 9 | exp. abstract | p = .796221 | p = .016843 | p = .020119 |
| 10 | exp. but unsure | p = .004366 | p = .005555 | p = .690988 |
| 11 | never exp. | p = .563381 | p > .999999 | p = .563381 |
| 12 | don't know | p = .316110 | p = .080975 | p = .563381 |

*Note.* Outcomes for Bhapkar (McNemar) homogeneity tests between phenomena for each aspect. Significant comparisons before correction (alpha = 0.05) are highlighted in orange. Significant comparisons after Bonferroni correction (alpha = 0.05/234) are highlighted in red.

**Supplementary S1.4**

*Mean confidence ratings for modality-specific questions*

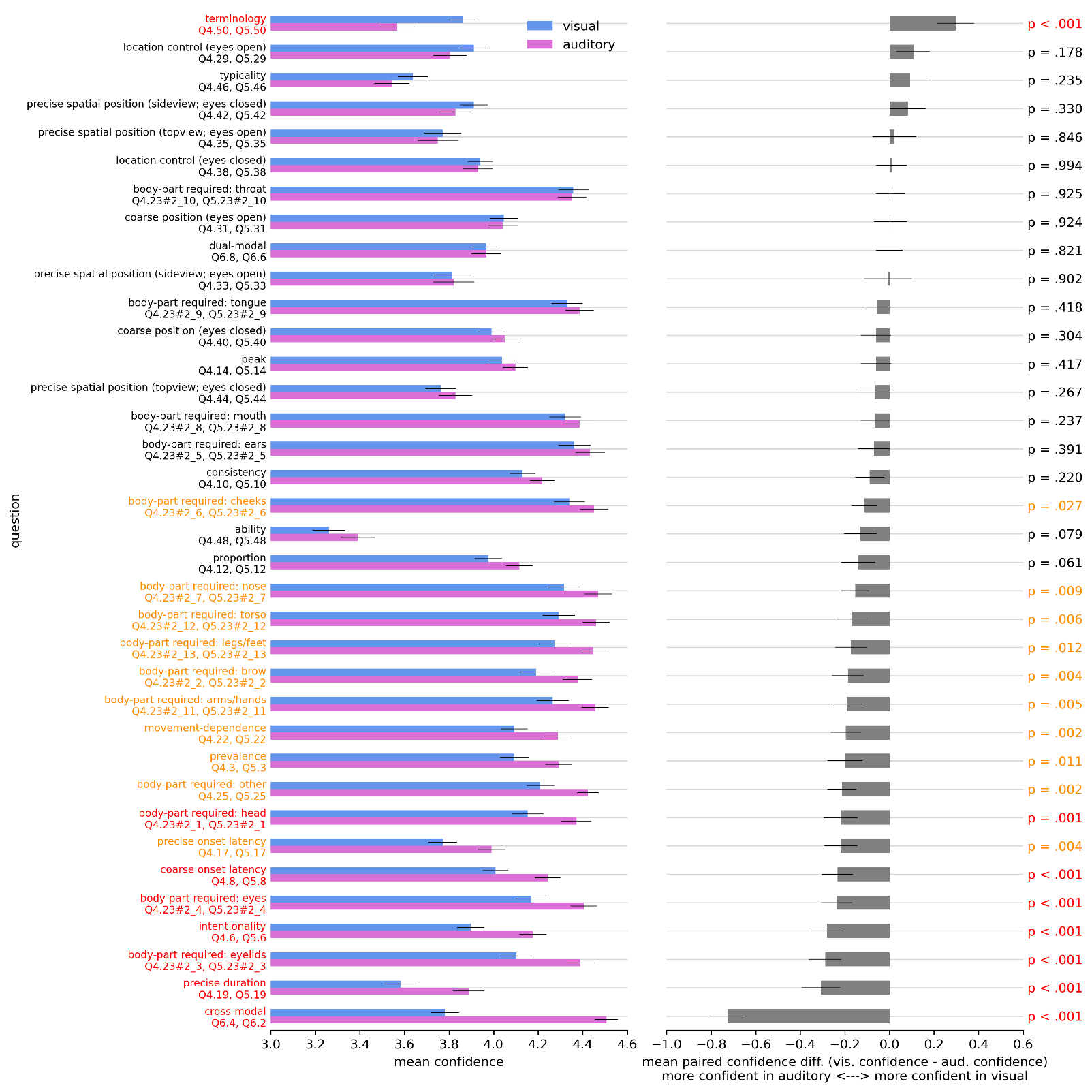

*Note.* Mean confidence ratings associated with each modality-specific question (left), and the corresponding mean paired differences in confidence between modalities (right). Uncorrected p-values shown for two-tailed Wilcoxon signed-rank tests against zero paired difference. Significant comparisons before correction (alpha = 0.05) are highlighted in orange. Significant comparisons after Bonferroni correction (alpha = 0.05/36) are highlighted in red. Error bars are the standard error of the mean. Not shown: wholeness (visual only) mean confidence = 3.88, error = 0.0040.

**Supplementary S1.5**

*Mean confidence ratings for thoughts, imagination, and dreams*

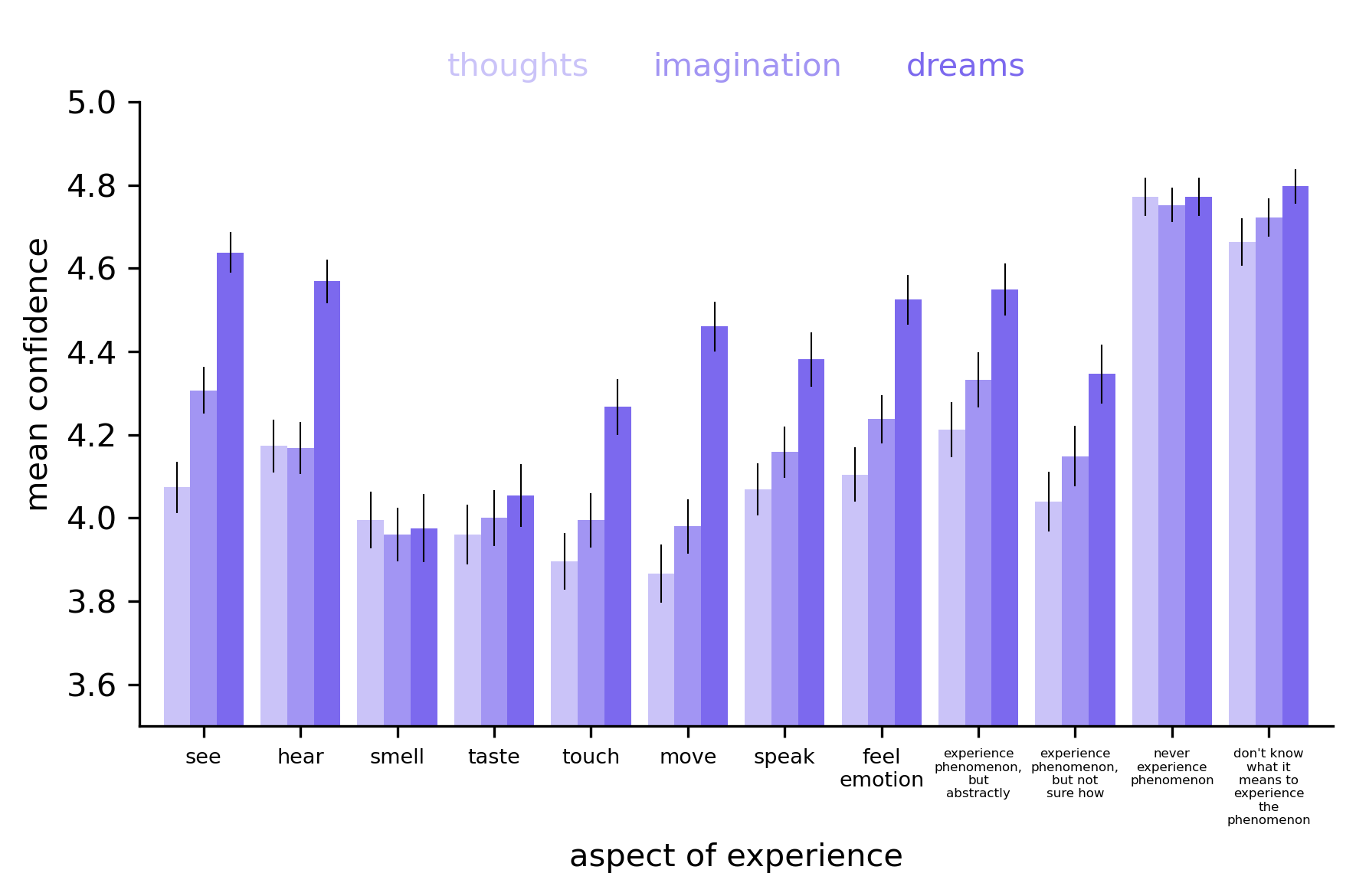

*Note.* Mean confidence ratings associated with each question concerning the presence or absence of a particular aspect of experience during thoughts, imagination, and dreams. Ratings are shown regardless of whether a respondent indicated that they did or did not experience that particular aspect. Error bars are the standard error of the mean. Statistical comparisons are detailed in S1.6 and S1.7.

**Supplementary S1.6**

*Within-phenomenon paired confidence comparisons for thoughts, imagination, and dreams*

| **comparison** | **aspect 1** | **aspect 2** | **thoughts** | **imagination** | **dreams** |
| --- | --- | --- | --- | --- | --- |
| 1 | see | hear | p = .103181 | p = .014496 | p = .014937 |
| 2 | see | smell | p = .359420 | p = .000003 | p < .000001 |
| 3 | see | taste | p = .166012 | p = .000044 | p < .000001 |
| 4 | see | touch | p = .016710 | p = .000008 | p < .000001 |
| 5 | see | move | p = .003114 | p = .000006 | p = .000139 |
| 6 | see | speak | p = .839099 | p = .023109 | p = .000004 |
| 7 | see | feel emotion | p = .525635 | p = .229256 | p = .011793 |
| 8 | see | exp. abstract | p = .077067 | p = .487258 | p = .218310 |
| 9 | see | exp. but unsure | p = .734302 | p = .067468 | p = .000028 |
| 10 | see | never exp. | p < .000001 | p < .000001 | p = .003064 |
| 11 | see | don't know | p < .000001 | p < .000001 | p = .001032 |
| 12 | hear | smell | p = .015040 | p = .002056 | p < .000001 |
| 13 | hear | taste | p = .003532 | p = .016898 | p < .000001 |
| 14 | hear | touch | p = .000121 | p = .008142 | p < .000001 |
| 15 | hear | move | p = .000032 | p = .003402 | p = .029523 |
| 16 | hear | speak | p = .133663 | p = .907462 | p = .000330 |
| 17 | hear | feel emotion | p = .321647 | p = .214504 | p = .424064 |
| 18 | hear | exp. abstract | p = .577799 | p = .005539 | p = .909771 |
| 19 | hear | exp. but unsure | p = .130261 | p = .852233 | p = .000779 |
| 20 | hear | never exp. | p < .000001 | p < .000001 | p = .000077 |
| 21 | hear | don't know | p < .000001 | p < .000001 | p = .000022 |
| 22 | smell | taste | p = .377817 | p = .292468 | p = .079934 |
| 23 | smell | touch | p = .060214 | p = .579582 | p = .000069 |
| 24 | smell | move | p = .058043 | p = .773915 | p < .000001 |
| 25 | smell | speak | p = .217864 | p = .002338 | p = .000002 |
| 26 | smell | feel emotion | p = .141646 | p = .000052 | p < .000001 |
| 27 | smell | exp. abstract | p = .009651 | p = .000004 | p < .000001 |
| 28 | smell | exp. but unsure | p = .593018 | p = .028871 | p = .000018 |
| 29 | smell | never exp. | p < .000001 | p < .000001 | p < .000001 |
| 30 | smell | don't know | p < .000001 | p < .000001 | p < .000001 |
| 31 | taste | touch | p = .164166 | p = .834442 | p = .000891 |
| 32 | taste | move | p = .142719 | p = .753945 | p < .000001 |
| 33 | taste | speak | p = .072061 | p = .018984 | p = .000007 |
| 34 | taste | feel emotion | p = .074535 | p = .000579 | p < .000001 |
| 35 | taste | exp. abstract | p = .005005 | p = .000031 | p < .000001 |
| 36 | taste | exp. but unsure | p = .382044 | p = .066321 | p = .000053 |
| 37 | taste | never exp. | p < .000001 | p < .000001 | p < .000001 |
| 38 | taste | don't know | p < .000001 | p < .000001 | p < .000001 |
| 39 | touch | move | p = .634131 | p = .732307 | p = .000819 |
| 40 | touch | speak | p = .013714 | p = .014745 | p = .037269 |
| 41 | touch | feel emotion | p = .009902 | p = .000472 | p = .000066 |
| 42 | touch | exp. abstract | p = .000400 | p = .000006 | p = .000004 |
| 43 | touch | exp. but unsure | p = .067416 | p = .066508 | p = .333638 |
| 44 | touch | never exp. | p < .000001 | p < .000001 | p < .000001 |
| 45 | touch | don't know | p < .000001 | p < .000001 | p < .000001 |
| 46 | move | speak | p = .000533 | p = .002231 | p = .072987 |
| 47 | move | feel emotion | p = .002585 | p = .000043 | p = .125735 |
| 48 | move | exp. abstract | p = .000111 | p = .000004 | p = .044851 |
| 49 | move | exp. but unsure | p = .036508 | p = .032361 | p = .064273 |
| 50 | move | never exp. | p < .000001 | p < .000001 | p = .000001 |
| 51 | move | don't know | p < .000001 | p < .000001 | p < .000001 |
| 52 | speak | feel emotion | p = .672305 | p = .166574 | p = .009435 |
| 53 | speak | exp. abstract | p = .096223 | p = .007876 | p = .006093 |
| 54 | speak | exp. but unsure | p = .802568 | p = .974659 | p = .621761 |
| 55 | speak | never exp. | p < .000001 | p < .000001 | p < .000001 |
| 56 | speak | don't know | p < .000001 | p < .000001 | p < .000001 |
| 57 | feel emotion | exp. abstract | p = .224237 | p = .134793 | p = .426843 |
| 58 | feel emotion | exp. but unsure | p = .459985 | p = .316239 | p = .012794 |
| 59 | feel emotion | never exp. | p < .000001 | p < .000001 | p = .000046 |
| 60 | feel emotion | don't know | p < .000001 | p < .000001 | p = .000017 |
| 61 | exp. abstract | exp. but unsure | p = .013059 | p = .009742 | p = .002108 |
| 62 | exp. abstract | never exp. | p < .000001 | p < .000001 | p = .000102 |
| 63 | exp. abstract | don't know | p < .000001 | p < .000001 | p = .000010 |
| 64 | exp. but unsure | never exp. | p < .000001 | p < .000001 | p < .000001 |
| 65 | exp. but unsure | don't know | p < .000001 | p < .000001 | p < .000001 |
| 66 | never exp. | don't know | p = .008165 | p = .258913 | p = .501570 |

*Note.* Outcomes for two-tailed Wilcoxon signed-rank tests against zero paired confidence difference for all possible comparisons within each phenomenon (i.e. within either thoughts, imagination, or dreams). Significant comparisons before correction (alpha = 0.05) are highlighted in orange. Significant comparisons after Bonferroni correction (alpha = 0.05/234) are highlighted in red.

**Supplementary S1.7**

*Between-phenomenon paired confidence comparisons for thoughts, imagination, and dreams*

| **comparison** | **aspect** | **thoughts vs. imagination** | **thoughts vs. dreams** | **imagination vs. dreams** |
| --- | --- | --- | --- | --- |
| 1 | see | p = .000047 | p < .000001 | p < .000001 |
| 2 | hear | p = .947901 | p < .000001 | p < .000001 |
| 3 | smell | p = .485268 | p = .793822 | p = .964429 |
| 4 | taste | p = .448061 | p = .256475 | p = .399762 |
| 5 | touch | p = .074525 | p = .000012 | p = .000651 |
| 6 | move | p = .041104 | p < .000001 | p < .000001 |
| 7 | speak | p = .105887 | p = .000025 | p = .001624 |
| 8 | feel emotion | p = .011326 | p < .000001 | p = .000010 |
| 9 | exp. abstract | p = .017465 | p < .000001 | p = .000043 |
| 10 | exp. but unsure | p = .094025 | p = .000012 | p = .003252 |
| 11 | never exp. | p = .466570 | p = .830888 | p = .596649 |
| 12 | don't know | p = .222946 | p = .026743 | p = .056249 |

*Note.* Outcomes for two-tailed Wilcoxon signed-rank tests against zero paired confidence difference between phenomena for each aspect. Significant comparisons before correction (alpha = 0.05) are highlighted in orange. Significant comparisons after Bonferroni correction (alpha = 0.05/234) are highlighted in red.

**Supplementary S1.8**

*Open-text responses to select questions (including from respondents who labelled sensory thoughts as hallucinations)*

**Other body parts required: visual** (Q4.24_3_TEXT)

1. mouth

2. I had to breath with my lungs and nose

3. My torso in order to fix my posture and my hands to put into my lap away from the computer

4. Torso

5. head, feet

6. When I closed my eyes and rubbed them with my hands I could see the O clearly.

**Other body parts required: auditory** (Q5.24_3_TEXT)

1. Jaw

2. mouth, throat, tongue

3. i had to breath

4. My torso in order to sit up, my eyes in order to close them and my hands to move them away from the computer

5. Stomach/diaphragm

**Alternative terminologies: visual** (Q4.49_4_TEXT)

1. Daydreaming (mental imagery?)

2. It is not as clear as 'mental imagery' seems to be. It is more of an idea than a picture in my mind.

3. Either imagining the letter "O" being drawn on a piece of paper in front of me, and the image of that being clear, or imagining a blank white/ black screen on which the letter "O" appears

4. imagination

5. Visualisation/ Mental Rehearsal

6. I can visualise but only for specific things. I can not see the "o" at all i dont think but I can see my outfits on my before i wear them or i can feel like i am on a beach or visualise how something will look on stage before i preform it but i can never seem to make myself see a shape, it has to be like a story almost. It is very confusing and i am not sure if this is common or not.

**Alternative terminologies: auditory** (Q5.49_4_TEXT)

1. imagination

2. imaginary sound?

3. auditory imagery

4. It is in my head and mouth. I have a constant internal monologue in my head.

5. Narrator

6. internal voice

7. I think I imagine somebody saying the thing / word and the sound they make, so some form of mental imagery maybe although I am unsure

8. Self-talk

9. Mental dialogue/stream of consciousness

10. Self-talk

11. mental phonetics

12. mental sound

13. Internal monologue

14. Inner monologue

15. auditory association or memory

16. Not too sure, may come under some form of mental imagery

17. Internal voice

18. hearing my own voice speaking

19. like a sound in my head

20. imagination

21. imagined sensory input/output

22. Hearing a voice say "o" to me. It can be me, a narrator, or anyone I want to imagine saying it to me.

**Extra comments: visual** (Q4.54)

1. if I do not give visual imagery my full attention it will disappear. in other words, the duration of the image being visible often depends upon my current concentration

2. I visualise the appearance of the letter O, most consistently, at eye level, even when my eyes are closed. I imagine the letter being drawn out similarly to how I would write it. However, I do not see a hand writing it. The letter simply comes into existence itself; like a loading circle stopping once complete.

3. The time it took for the content to become fully visible did not seem to speed up every time it was prompted

4. I'm not entirely sure what you mean by asking where the visual thought is in relation to your body, I don't particularly think of my thoughts as having a location. This was an interesting survey. :)

5. I can see 3D/2D objects in my mind usually on a black background. I was seeing the O as a white 3D letter.

6. it felt more blurred around the edges, and my eyes had to strain to focus in on it

7. When I focus solely on visualising the letter O, the image feels like it is in the center of my head and towards my forehead. When my internal monologue begins and I try to simultaneously imagine the letter O, the image moves to the right side of my head, more towards my temples.

8. It is almost like a drawing process -- creating the circle by like visually linking two ends which are sort of being circularly drawn towards each other

9. The visual image usually comes and goes, and does not stay in my mind consistently for a period of time.

10. I imagine it written on a green chalkboard

11. The 'o' forms at the bottom before the entire letter spreads out as I move my eyes behind my eyelids and I see the whole thing right in front of me in the darkness

12. I found deliberately visualising the O difficult but, some other thoughts I just naturally visualise, especially things that make me anxious

13. I would trace the letter in my head

14. when experiencing thought content visually with my eyes open i experience it more as a distortion in the environment. For example when i was thinking about the letter "o" and looking straight ahead i saw it as an indent in the wall in front of me in the shape of an "o".

15. I move my eyes to follow the outline of the letter O and it's more clearer when my eyes are on certain parts of the letter.

16. Im not sure why, but at the start my vision of the O came quite clearly. But over time, it seemed like it was harder to conjure a visual image of O in my mind.

17. when visualising the letter O I would almost involuantarily trace the shape of the letter in my mental imagery to get a better idea of the shape

18. I saw multiple different O's that seemed to be in different places

19. I can clearly see the letter 'O', its appearance is clear to me, and I can manipulate its size and appearance (i.e. colour).

20. To focus on the letter I have to think of it in the sentence of 'the letter 'O'' in the instructions, if I specifically try to focus on the O it take more concentration and I can only really focus on one aspect of it at a time

21. I'm not too sure if I had experienced a thought as all I could see was darkness.

22. when visualising the letter O with my eyes closed, it appears almost like a shadow, as opposed to having colour or light.

23. The letter 'o' often appears blue in my mental imagery, although can change. Can switch between fonts and backgrounds as well.

24. I am a second year Psychology student. I am too very interested in the internal thought processes of individual people. I, myself, do not have an internal monologue and feel that I think in a different way to many people. Sometimes I think visually and other times I think in terms of concept or objects.

25. The 'O' was transient and disjointed.

26. the images in my mind flicker between lots of different images, like the sesame street O letter block, and a magazine O, and an O being written, it flickers like a stop motion film

27. Often times I would envision drawing or writing out the letter.

28. At first, I saw the letter O in the early stages of this experiment. However, it became more difficult to see the O when I tried again.

29. 'O' was clearer depending on what 'O' looked like, some 'O's' were harder to think and visualise of and there were some 'O's' that were easy to visualise and this was due to a change in font or roundness or style of writing.

30. I felt that I needed to create an environment where O could be in from of me, I thought/imagined from a bird eye view.

31. When holding for 5 secs the image began to blink and start to fade

32. When I close my eyes I see content in either 2D or 3D almost as if on a screen projecting onto the back of my eyelids. I find it difficult to create large objects in my head without focusing on a smaller part of them, with the rest almost fading but feeling like it is still there.

33. Much more visual with nouns than abstract kind of things such as the letter 'O'

34. I'm pretty sure I just have aphantasia. I really only see black when I close my eyes and think and when I do think (I guess you can call it that) it's more like silently repeating O over and over again. Like undergoing the breathing motions for saying O, but not actually opening my mouth or making noise. So all in all, I really cannot see anything when I think about O, so apologies if my responses were redundant

35. It is more of an idea than a picture. I could describe the 'mental image' with words and detail and I could attempt to draw what I believe to see however I can't see it when I close my eyes, it is not like I have turned on a tv when my eyes shut.

36. Each time I thought about the letter O there was a slightly difference experience. Sometimes I saw the whole letter where as others it was segments at a time.

37. I'm stumped a bit on how visual of a thinker I am since I can't pinpoint whether I am actually seeing the object or if in my head I just know what it is and am thinking about it. I can create elaborate scenarios in my head and I know what things look like in it but when someone asks me to picture something and I close my eyes I struggle to visually picture it. I know what it looks like and I am imagining it in my head but am I actually seeing it? I think my visual thinking is also better when I am in the mood ie. if someone tells me to imagine something I find it harder than just naturally daydreaming.

38. I imagined the letter O in a book

39. I can use my imagination to visualize all sorts of visual scenarios. In my head, on my body, in the environment, in made up environments etc.

40. The way I visualised it with my eyes closed was as a circuit in that I traced the circle over and over, it wasn't always just fixed (sometimes it was fixed though).

41. I find it easier to visualize places, objects and people, rather than numbers and letters

42. When my eyes were closed, O looked black and grey. When my eyes were open, O looked yellow and white.

43. It seems relatively inconsistent..? I'm not quite sure, sometimes it appears very clearly as an image but sometimes it feels "just out of reach" in some way (in the same way that something can be on "the tip of your tongue", it felt like it was on the edge of my mind at times)

44. Thought content is dependent on the mood and environment.

45. I mentally draw out the O when I close my eyes

46. When asked to picture an O in my head, i pictured just merely the letter o upon a black background or letter O as it was written out in the question - not too sure if this is doing it right?

47. I feel like I was drawing it starting from the top and going round clockwise with my eyes

48. The visual is always very fuzzy and hazy

49. i can change how it looks, bold, large, small, move it around, be on it's own or on a piece of paper

50. my eyes trace out the shape of the 'o' when closed

51. Often, it's as if I was drawing an O in my head

52. I have average visualization but I am able to imagine objects better in 3D from various perspectives

53. When I close my eyes it appears right in front of my eyes, like with a black background behind it, when my eyes are open it appears on the object or wall I'm looking at

54. It's almost like there's another dimension past/in front of my eyes that I imagine the thoughts in

55. It is fragmentary and indistinct, and comes and goes.

56. At the beginning I briefly saw the letter O with my eyes closed but later in the study lost this ability. When I think about something I can see it in my head but not with my eyes...It's hard to explain.

57. It's hard to tell whether I'm actually visualising the 'O', or if I'm just imagining myself visualising the 'O', with the 'O' stored more conceptually than visually.

58. I cannot visually see my thought content, regardless of the technique used

59. I associate thought content with art. Whether it is music, visual arts, or I find that I understand better when visualising through some form of art.

60. I simply pictured the way 'O' would look typed on a word document

61. Sometimes I picture it in the dark, and the shape. Other times I think of a fields and the letter in the background. When I think of this I feel more stimulation in the back of the brain. Feel more front with picturing the shape in the dark of the letter.

62. It seems to be easier to focus on the specific visual image when I 'make it' / 'allow it' to move around (eyes closed). Making the image flash (eyes closed) helps prevent other thoughts from distracting me and causing me to lose focus (image disappears). Being still helps me maintain the image when my eyes are open.

63. It changed colour when I pictured the 'O' with a different coloured background

64. It's hard for me to think about text visually. Imagining the letter "O" is so much more difficult than imagining, say, an apple. Imagining letters and text is quite unnatural and has to be forced.

65. Like there was a black space inside my head except for my thought.

66. I can not see just an "o" by itself as tested at all. It feels so unnatural to me but when I tried to see myself wearing an "o' on a t-shirt i could see the o for the first time. I know that was not instructed so i didnt say I could see the "o" but i felt it was important to let you know this. I can visualise things but only in maybe a non abstract sense (i think). I can only visualise when it feels grounded in something that i know.

67. Sometimes I feel like I can control it, sometimes not.

68. I can't directly *see* it per se, it's a very strange thing to describe how I view my thoughts.

69. I feel like i could imagine an O but it was hazy, like it was struggling to stay complete in my mind, maybe due to my lack of focus tho.

70. i draw the image in my head e.g. a drawing of the letter O is drafted in my head but disappears

71. i could trace it out with my eyes repeatedly to keep the letter consistently visualised

72. The visual thought content in my mind sort of throbs and has movement, but remains continunously present.

**Extra comments: auditory** (Q5.51)

1. feels more like a mental sound than an audible sound. as if I am imagining what the audible sound would sound like

2. I was just hearing "O" in my head.

3. Just feels as though I can hear it....within my head -- it is the same as my thoughts right now, except that I am deliberately being aware of them

4. Scenes play out inspired by sounds

5. I could hear the thought of the O in my inner ears, but could only tick one place on the diagrams, but its both ears.

6. It wasn't audible but it felt similar to how I feel when dreaming.

7. My thought content is like an inner monolouge in which I am speaking to myself and repeting what I am reading internally

8. When I picture the sound of the letter 'O', I have complete control over it, whether it was the sound itself; could keep changing the sound of the 'O' (i.e. screeching, bouncing, static, etc.). I could completely manipulate the location of the sound in my head (i.e. I could have it spinning around, towards the front, back, sides, etc.).

9. i heard it in two spots, or like a band from ear to ear rather than a specific spot

10. I did not think the sounds have a location

11. i hear it in my own voice

12. can manipulate the 'o' sound to get different phonetic sounds

13. Unlike the visual imagery, I have previously experienced visual imagery in my life, just not in this particular task. With audible imagery, I have never ever experienced an internal monologue. In fact, I only found out people had internal monologues when I was about 16 or 17. I used to think internal monologues were only ever used in movies to help verbalise a characters thoughts, not a real experience.

14. Wasn't able to hear an 'O' sound in my head but believe I experience auditory sensation when thinking about certain things, e.g., going over a speech in my head.

15. I didn't hear the sound of 'O' or like I didn't experience 'O' having some sort of audible property, this is probably because I think a letter does not make sounds other than when it is read. I believe I would have heard an auditory property of my thoughts if it were something like the sea. So when it comes to things which actually make sounds, when I think of them I would hear the sounds that they make but when it comes to things that are like letters or numbers I don't hear their sound. I think of 'O' whilst saying 'O', but the letter 'O' does not emit a sound, so the only audible thing that is happening is me reading out the letter 'O' rather than 'O' having a unique sound of its own.

16. for me, source of sound needed to be there, I imagined an actual O letter speaking O

17. Easier to hear words and sentences rather than a letter

18. I said previously that when I am told to think of something, it's more of me saying it without actually *saying* it over and over again. I undergo the breathing motions of saying it, but I don't actually hear it, hence why I'm only mostly sure about not being able to hear anything when I think.

19. In the first section it asked if we hear anything when visualising the letter 'O' I said no because I interpreted that as an (hallucinative) external noise/voice and I do not hear that but I do have an internal monologue in my brain that is reading, saying and hearing 'O'

20. When thinking of the sound, I was hearing my voice say the name of the letter O over and over.

21. I don't think I can control how "loud" my thoughts are. I know the difference between screaming and whispering in my head but they sound the same level of loudness.

22. It feels like I am saying it to myself quietly in my head

23. I can hear my own voice and differentiate between tone but the volume stays the same

24. Its a consistent repetition of the O sound

25. When I think about the sound of the letter 'O' it sounds like someone is whispering it to me

26. I feel like I'm speaking, moreso than hearing. But you can hear yourself speak and that is what I 'hear' when thinking.. I think

27. As opposed to imagery, sound appeared to be in my head.

28. it's easier to hear things in my head with my eyes open, rather than closed. It's easier to locate where i'm hearing things with my eyes open (i hear things in the same spot with my eyes open vs. closed, but it is much easier to pinpoint with my eyes open). when i hear things in my head, it's very distant - I almost didn't recognise it was happening before. The auditory process is my own voice reading/speaking things back to me.

29. the dots in the head (near the right ear) for this page and the previous page should also be reflected on the left side. Sound seems to stem from region of both ears. When thinking of a sustained 'O' sound, it was difficult to continue the sound when I was breathing in. It was like I was making the sound in real life (requiring the expulsion of air from my mouth to sustain the sound). The sound, in my head, would sometimes pause when I inhaled.

30. Much easier to hear text than to visualise it, but it's much harder to hear "objects" than to visualise them.

31. I'm not sure if it counts as it's like my own voice is talking to me.

32. I'm not really sure what is meant by the location of my auditory thoughts.

33. I've talked to my mum about this and she says that when she reads a book there is no audible voice in her head as she's reading, in my head when i read, i can hear the text in my own voice, my friend hears in other peoples voices, like characters, as well as her own when she reads.

34. Throbbing, loud noise the more I think about it.

**Extra comments: visual-audio interactions** (Q6.10)

1. I think I started to visualize words once I imagined they were being spoken to me, but I couldn't visualize a single letter

2. i cant control the spatial location but no answer available for that

3. I am very interested in what the results of this study show, I do not know if the way I think is considered 'normal' and I have always wanted answers as to how I really think because I find it difficult to describe myself.

4. Q6.7 was confusing because it asked if I see an image of the letter with sound and I only experienced an image and I did not here any sound and this question did not have the option for me to choose, "No, I only see what I'm thinking in some way" like in Q6.6. So I ended up choosing "No, I do not see or hear what I'm thinking about in any way" for Q6.7

5. When perceiving the sound of the letter O the experience seems to mimic the real life experience of saying the sound out loud- feeling almost as a ghost or shadow experience of actually saying the letter. For visualising the letter the image exists in a much more conceptual space not as connected to the real life experience of seeing the letter O written or typed out.

6. again, I find i easier to imagine objects, places and people, rather than letters and numbers.

7. Q6.3 - to clarify when someone is speaking to me I see visual images representing the word automatically, not the word it self. I answered the question on the premise that it was asking if I saw the words spelt out in alphabets when someone was speaking it to me. n

8. Hearing things comes more naturally to me in everyday life.

9. Visual - the O was very visible at first, but it got much harder to see it the more i was asked to picture it. This did not happen with hearing; it was easy to hear the O the entire time.

10. when I recall a memory I also can hear the sounds in my head, in a much clearer way than I can see the images (although I can picture it too). I knew I had a visual memory, but now it is clear to me that it is more auditory!

11. The sustained 'O' sound seemed to muffle over time, whilst the sound of the sentences I read sounded clear throughout.

**Extra comments: thoughts, imaginings, dreams** (Q7.4)

1. I think my sensations during thoughts, imaginations and dreams change in intensity depending on the content, and how emotionally salient it is.

2. Dreams are the most vivid, followed by imaginings

3. When it comes to smelling, tasting and feeling things in my dreams it is a little difficult to picture something I am not familiar with. What I mean by that is if I were to dream/think about a certain food I have never eaten before in my life, it is difficult to picture what it smells, tastes and feels like, however, there is a sort of substitution for those missing parts as though it was a "default" choice to a question if no answer was given. What is substituted can vary based on previous experiences such as if I experienced something similar then the taste, smell and feel of it can be somewhat reminiscent of the previous experience. If I can't round something off from a previous experience the best I would describe the substitution would be 'bland' or 'plain'.

4. I do occasionally sleep talk and only recently started remembering my dreams

5. I have very vivid dreams and imaginings and so can quite often immerse my senses into them quite well.

6. i frequently dream and frequently day dream, sometimes I struggle to determine what was day dream and what was the reality around me particularly with sound

7. I have very vivid dreams. Often I remember them, but sometimes I don't.

8. I don't usually get dreams, but similarly to thoughts and my imagination, when it does happen, it's usually black and I simply just know what is going on. I don't exactly experience it in terms of my senses or emotions, but I know what is happening as it happens

9. Visuals and auditory stimuli in my dreams can be very vivid

10. i don't remember much about my dreams because i don't give it much thought, but i definitely do wake up scared or elated sometimes depending on the dreams

11. I wish I did not think, imagine and dream so vividly all the time. I am often "day dreaming" thinking about that happened, reliving the situation, and I waste a lot of hours imagining things and situation, fantasising "what ifs" like if I had to create a movie plot about something. It is annoying. I like my dreams as they are mostly fun and nice, but when they are nightmares I almost get traumatized by them. I still remember vividly dreams that I had years ago. Now I am wondering if this is common/ normal. Probably not!

12. I have more vivid dreams which I remember when I try go back to sleep at around 7am when i've had a bad/ disruptive sleep.

13. I have felt heat, wind and water in my dreams.

14. dreams are by far the most visual experience than anything else

15. The sensation of touch is the most foreign in a dream, it feels very different to a real touch.

**Extra comments: direct multimodal comparisons** (Q8.24)

1. Again, a lot of the intensity of the sensation depends on the emotional content, and often seeing and hearing thoughts occurs concurrently if it happens, the clarity of such sensations can depend on emotion too.

2. When I labelled something as "mostly sure" it refers to what I have written in the previous section (substituting in taste, smell and feeling).

3. seeing my thoughts isn't hard, it's just it's alot easier to hear my thoughts

4. I have to consciously add smell to my mental imagery

**Thinking vs. imagining: is there a difference?** (Q9.14)

1. I do not think so. The reason I believe this is because the brain is not the primary visual or auditory organ, so it would be unable to distinguish the difference between 'thinking' and 'imagining' in addition, those words are merely abstract concepts and have no real meaning aside from what we humans place on them.

2. Not really, both require as much effort as each other

3. Thinking requires more sound and imagining is more visual

4. no

5. thinking something is something immediate but imagining is more day dreaming

6. Imagining seems to focus more on the experience of conjuring an image/sound/thought, whereas thought seems to be more aligned to evaluating this imagination

7. Yes I do believe there is a difference. My experience of thinking is more guided and strict, the thoughts are less creative, the imagery is definitely based on real life examples. Imagining allows for more creativity, things look less 'real-world-like'. For instance, when asked to think about the appearance of the letter 'O', I automatically visualised exactly that same 'O' from that sentence, in the same font. Whereas when asked to Imagine the letter 'O', and image of a large light blue 'O' with black outlines came to mind.

8. I think 'imagining' has a connotation with visual imagery whereas you can think about something without necessarily imagining it (with visual imagery). I can think about my response to this question but I haven't imagined anything visually.

9. When thinking about something, you might list the characteristics of it in your mind, but when imagining it, you would think of the appearance/sound of it.

10. imagining is more subjective than thinking as brings in more personal interpretations. whereas thinking is more clear and straight to the fact

11. Thinking feels more deliberate and controllable, imagining is less deliberate but i can control stopping it, maybe it becomes thinking at that point.

12. Not too sure, but I think being told to imagine something immediately makes it a more vivid experience than thinking

13. I am not sure but when I'm told to do something (e.g. translating, think/imagine something) I can't really. But I daydream very easily and very often about something I'm thinking of.

14. Imaging is more vivid as the subject matter becomes more complex.

15. I do not.

16. Not really

17. Yes, thinking is what happens naturally and your brain processes things without the excessive need to use any of the 5 senses in order to process it. Imagining involves using one (or more) of the 5 senses to give the thought real world context.

18. Thinking is to just well think about it or for something to be on your mind whereas imagining is to conjure a sensory image of something

19. I cannot tell the differecne mos of the time

20. There could be when it comes to their definitions but I don't really defer the thought that they mean the same thing. Honestly, it would seem annoying to have to refer to what I experienced as a specific thing (i.e. saying that I "imagined" something but I'm then told I actually "thought" it). If I were to define the two terms I'd say "imaging" seems more expansive and large-scale while "thinking" were to be a simple thought that pops into your head while reading something or solving a problem.

21. Yes but more for thinking about/imagining experiences or events rather than things such as letters

22. No

23. no

24. Yes, imagining is perceiving the appearance of a certain object/picture, while thinking is to have a particular feeling/emotion to a certain object/place/environment/situation.

25. I think that for myself, they are qualitatively the same but the difference lies in the actual content of what is being thought or imagined about.

26. I would differentiate them on that thinking is from real life experience but imagining can be more abstract.

27. I believe that thinking involves a logical structure whereas imagining can be illogical.

28. Slightly, external stimulus's could trigger different sense

29. thinking would be more based in memories and imagining would be made-up scenarios/images

30. Imagining involves conjuring up an approximation of the form of the thing you're thinking about, whereas thinking doesn't have to involve that

31. I process the actions as one in the same

32. I think that imagining something requires the integration of visually recreating the object within the mind and hence is an extension of thought processes to try and understand the thoughts

33. No, I think that its the same process in the brain. Its just that 'thinking' about something may be something that has already happened and 'imagining' something ,ay be something that has not happened.

34. I think it is more the situation for each, where imagining is more personal and hopeful.

35. I think imagining and thinking are part of the same process and are not that different

36. When I am asked to think about something, I place it within certain contexts and try to think about all its features and relations. When asked to imagine something, I try to picture it physically.

37. I both think accidentally and imagine accidentally but in the imagining I making things up, even if sort of subconsciously it is some sort of story (even when its real) but thoughts are more like observations, I dont change them actively (unless I am trying to calm myself down) and I often have many at the same time both fleeting and long. This is unlike imagining which sort of happens in the background in a consistent way if its 'subconscious' or if intentional it is constant until i edit something

38. Not really

39. imagining something involves more visual imagery whereas thinking can be just thoughts and emotions etc

40. not really

41. Imagining for me requires more effort and more likely to be audible and visual.

42. There seems to be a difference but the difference is very fine that they border on each other. Like we use think when it comes to something like, thinking of an answer to the question rather than imagine and yet we can also imagine an answer to a question. I guess they have different connotations but they both stem from our use of thought

43. thinking- thinking the concept of it imagining- what it looks or sounds like

44. I'm not too sure about this but I feel like when we think about something, we are;t as emotionally involved in comparison to if we imagine something.

45. Thinking takes effort and is more literal. Whereas, imagination is more spontaneous and involves creativity.

46. I do believe there is a difference because I was unable to think of the appearance of the letter 'O' but I have been able to imagine things visually

47. imagining is thinking about something more vividly/in more detail

48. In the case for hearing, i can conjure a consistent sound of someone saying the letter O, but when actually experiencing it some people pronounce their O's with different accents which are not so familiar to me

49. I think the position of the person when thinking something is very much in the first person and imitates that experience in real life, whereas imaging something involves a much more constructed situation which isn't always first person or congruent to real life experiences.

50. Thinking about something for me involves retrieving a premade file of something, whereas, imagining something is like a developmental process which starts from scratch and gradually shapes into what i see

51. How based in reality it is and how much concentration is being put into it. When I talk, I'm usually not thinking about what I'm saying. It's kind of odd because I am still able to filter myself and know what I can and cannot say. I just say it as I need to, but when I do need to actively think about something, it automatically becomes something that I need to put effort into, because I don't usually have very active thoughts or thoughts that I notice are occuring. Imagining on the other hand is usually more fantastical. Things that currently do not exist or occur. When I get bored, I tend to actively try to think of different scenarios and things that will never occur (though this may be because I am someone more interested in fantasy rather than reality).

52. Imagining something feels like it is more of a vivid thought than just thinking about something

53. no

54. no, i think imagination is just a thorough more creative process

55. Thinking is like recall and memory based whilst imagining can be something you haven't seen before, thus making up the image in your mind.

56. Thinking about something needs to draw on a past experience, where as imagination can be on more fabricated things.

57. I honestly have no idea but to me when I think of imagining as having at least a visual aspect but thinking can be either auditory or auditory + visual? I think imagination may be border more towards dreams as well.

58. I think they are similar. However, I think imagining means putting yourself into some situation. or imagining some sort of situation and scenario. but for thinking, you might further think about something like possibilities, critically evaluate and analyse something.

59. Imagining about something is when you get lost in thoughts and let the creative process take over. Thinking something is more automatic with no effort.

60. Yes, thinking is automatic, imagining is more of an active experience

61. No, I experience it the same.

62. Yes. I think it's easier to imagine sounds due to my constant inner dialogue

63. Imagining is deeper

64. Yes, I think imagining is more unconscious

65. I believe the difference is subtle, but I feel like imagining something is more me replicating that thing in my mind, whilst thinking is more like me stepping through the process of that thing. But since the difference is so subtle, I would probably use these words interchangeably if I wasn't asked directly like I am in this question.

66. I think it's the same

67. Yes - you can imagine something that is not real.

68. Yes. Thoughts seem automatically and more auditory-based, like an internal dialogue. Imagining something seems to require more effort and better resembles projecting an image up onto a screen to watch.

69. imagination is more focused and controlled, but since 'thinking about something' is a prompt then they are essentially the same because both words act as prompt commands.

70. imagining allows for a more creative picturing while thinking creates a more realistic picture, however i find no difference with sound

71. No

72. Thinking about something involves less detail than imagining something

73. imagine might be more visual or sensoral, thinking might just be about characteristics with "o" think round, or circle or 360 degrees

74. I think thoughts occur naturally and voluntarily? Imagination requires effort to actually formulate a thought content.

75. Yes. In my imagination I can be more creative. If I think about something I apply the boundaries of the world we live in.

76. they're the same to me

77. I think when I think about something I don't see or hear what I am thinking about, however if I actively imagine something I am doing it by seeing/hearing those imaginations.

78. If I am thinking of a scenario I am hearing the words but if I am imagining it is more similar to dreaming i.e. I can see, hear, smell, taste and feel.

79. Thinking is where you're trying to achieve something, but imagining is when you just let your mind wander off.

80. Yes. To me, thinking about something includes imagining it, but you don't necessarily have to imagine something to be thinking about it. Imagining is creating something tangible in your mind, whereas thinking also includes abstract concepts and other things.

81. imaging the letter 'O' I am imagining a woosh noise of making the shape of the circle while thinking about the sound of the letter 'O' i think about the way you'd say it.

82. This is such an interesting question! I have never thought about it. I think neurologically it's the same because they experience feels the same.

83. It depends on what it is

84. Imagining to me is like 'being there' whilst a thought is a lot more simple.

85. Yes, I think there is a difference. Imagining has less constraints, it is a more fluid, unforced process. Thinking about something is more to do with referring to the knowledge you already have of the thing.

86. imagining is more visual, thinking includes hearing more.

87. No

88. To me, in imagining something, I am recalling or creating any associations i have with the object of imagination, while if I am thinking about it, I am trying to analyse or consider specific elements or qualities of the most solid imagining of the object.

89. no

90. Thinking is recalling the concept and your opinions of it, imagining is purposefully creating a mental image to provide yourself with an experience

91. i know the words have different meanings but when I think about something it feels the exact same as imagining something

92. Yes because if i think about something I can not picture it as vividly as if I imagine it.

93. No

94. no difference for me.

95. I think there is a difference between thinking and imagining something when you need to force yourself, but if the action happens autonomously then I think they are the same.

96. Visually and audibly they are the same, but "thinking" implies more focussed intellectual attention, while "imagining" indicates more freedom of thought.

97. Imagining things usually refers to use of imagination and creativity. imagining my involve more vivid sensory 'input,; and more focus. Whereas, thinking normally results in mental images almost automatically and have a more realsitic sense to them. These two things are hard to differentiate by purely thinking about and imagining 'O' - too simple to make differentiaation.

98. Yes, thinking involves analysing, but imagining is purely creative

99. in certain circumstances I believe so. But when it comes to visualising and hearing, I believe there is little difference.

100. Thinking is more abstract, while imagining is trying to create a sensory experience. Like, when I read a work of fiction, I tend to use my imagination to create scenes in my head, but when I am studying or read non-fiction, I tend to use my thoughts to internalize the logic of what I'm reading.

101. I personally do not think there is a large difference in thinking about something and imagining it, unless it is talking about something i have not heard or seen about. then, to imagine it would be to guess what it looked/ sound like etc., while thinking would not really apply as i would not have a clue on what to be thinking about.

102. No

103. Thinking is more verbal, whilst imagining is more visual for me.

104. I think imagining includes a visual aspect whereas thinking does not necessarily mean you are able to visualise that mental process.

105. yes, thinking about something felt more uncomfortable and forced, it was the letter "o" that we had to see but when i could imagine "o" i felt that it was in many different forms. "o" was painted on a building or on a mountain when i could imagine it.

106. When I think about something, I put mental effort into coming up with an answer logically. When I imagine something, I roll with my gut feeling and focus on the first thing that comes to mind.

107. I think imagining something is when you don't have much experience or knowledge about it, for example imagining what something looks like because you don't actually know completely what it looks like. Thinking is more so a recollection from previous knowledge or experiences, for example thinking about a bowl of pasta you know what it looks like, smells like or tastes like and so you recreate that sensation in your mind.

108. thinking about something is an active conscious process leading to thought that yields benefit such as thinking about a chemical reaction leads to a result of understanding the reaction pathway. Imagining something is unconscious and therefore not active process of thought is followed, hence no beneficial yield emerges.

109. for people who don't experience the. usual aspect of thoughts, yes there is a difference. imagining is a specifically visual thought. some people just have the visual aspect of thoughts constantly in operation.

1. The study was run only on SONA students
2. Students were not actually penalised in any way for answering incorrectly, or for not completing the study
3. All confidence questions were arranged horizontally, with “Completely unsure” on the left and “Completely sure” on the right
4. An error meant that Q4.23 and Q5.23 response options were not exactly matched. Q4.23 uses the no-experience option "I did not see my thought in any way", while Q5.23 uses "My thought content was not audible in any way".
5. Attention check
6. An error meant that Q4.28 and Q4.37 response options were not exactly matched. Q4.28 uses the option "No, my thought content was visible, but I could not choose where it seemed to be located", while Q4.37 uses "No, my thought was visible, but I could not choose where it seemed to be located".
7. This question was hidden if participants reported "My thought content was not visible in any way" on Q4.30
8. This question was hidden if participants reported "My thought content was not visible in any way" on Q4.39
9. Attention check
10. An error meant that Q5.28 and Q5.37 response options were not exactly matched. Q5.28 uses the option "No, my thought content was audible, but I could not choose where it seemed to be located", while Q5.37 uses "No, my thought was audible, but I could not choose where it seemed to be located".
11. This question was hidden if participants reported "My thought content was not audible in any way" on Q5.30
12. This question was hidden if participants reported "My thought content was not audible in any way" on Q5.39
13. Cross-modal auditory imagery
14. Cross-modal visual imagery
15. Auditory imagery induced by visual imagery
16. Visual imagery induced by auditory imagery
17. Attention check
