## Supplementary S1-7 for "Comparing mental imagery experiences across visual, auditory, and other sensory modalities": Supplementary S3.ipynb

```
# %% [markdown]
## Imports

# %% [code]
import matplotlib.pyplot as plt
import matplotlib.patches as mpatches
import matplotlib.image as mpimg
import matplotlib.gridspec as mpg
import matplotlib.cm as mpcm
import matplotlib.colors as mpc
import cv2
import pandas as pd
import numpy as np
import os
import textwrap
import random
import textwrap
from statsmodels.stats.contingency_tables import Table
from statsmodels.stats.contingency_tables import SquareTable
from scipy.stats import wilcoxon, pearsonr, mannwhitneyu, levene
from IPython.display import Markdown as md

# %% [markdown]
## Load files

# %% [code]
raw = pd.read_csv('Supplementary S2.csv')

# %% [markdown]
## Excluded subjects
- Too fast (< 20 mins)
- Non-native speaker/reader
- Inattentive (failed attention checks)
- Classified sensory experience as a hallucination
- Reported having a known neurological, psychological, or psychiatric condition 
- Didn't answer Q4.52 (before this was corrected to be a required response)

# %% [code]
# Create a table with question numbers (as strings) as the columns and actual 
# question text is the entry. Convert it to a dictionary. Can look up question
# number to get corresponding text.
qs = raw.iloc[0,:]
qs = qs.to_dict()
filtered = raw.iloc[1:,:].copy()
originaltotal = len(filtered)
original = filtered.copy()
print(f'{originaltotal} before exclusions')

# Start excluding subjects
# Filter out those with disorders
before = len(filtered)
filtered = filtered[filtered['Q9.3'] == 'No']
after = len(filtered)
print(f'{before-after} excluded by disorders screen')

# Filter out inattentive folk
before = len(filtered)
filtered = filtered[filtered['Q4.26'] == 'Always']
filtered = filtered[filtered['Q5.26'] == 'About the same']
filtered = filtered[filtered['Q7.2#1_1'] == 'Agree']
filtered = filtered[filtered['Q7.2#2_1'] == 'Completely sure']
after = len(filtered)
print(f'{before-after} excluded by attention checks')

# Filter those who didn't answer Q4.52 (I accidentally didn't make it a required question)
before = len(filtered)
filtered = filtered[filtered['Q4.52'] != '-99']
after = len(filtered)
print(f'{before-after} excluded by noncompletion')

# Filter unrealistically-fast responders
before = len(filtered)
filtered = filtered[filtered['Duration (in seconds)'].astype('float') >= 1200] #1200
after = len(filtered)
print(f'{before-after} excluded by duration')

# Filter out non-English
before = len(filtered)
filtered = filtered[filtered['Q9.4'] == 'Yes']
filtered = filtered[filtered['Q9.5'] == 'Yes']
after = len(filtered)
print(f'{before-after} excluded by language')

# Check for no underage
before = len(filtered)
filtered = filtered[filtered['Q9.1_1'].astype('float') > 17]
after = len(filtered)
print(f'{before-after} excluded by age')

# Filter those who named the experience as a hallucination. 
# Preserve the dataset with hallucinators though to show how many responded as such later.
prehall_filtered = filtered.copy()
filtered = filtered[filtered['Q4.49'] != 'Hallucination']
filtered = filtered[filtered['Q5.49'] != 'Hallucination']
totalsubs = len(filtered)
print()
print("Total subjects included (nonhallucinators): ", totalsubs)
print(f'Median age {filtered["Q9.1_1"].median()}; {filtered["Q9.2"].value_counts()["Female"]} female, {filtered["Q9.2"].value_counts()["Male"]} male')
print()
print("Total subjects included (inc. hallucinators): ", len(prehall_filtered))
print(f'Median age {prehall_filtered["Q9.1_1"].median()}; {prehall_filtered["Q9.2"].value_counts()["Female"]} female, {prehall_filtered["Q9.2"].value_counts()["Male"]} male')


# %% [markdown]
## Neutralise response labels
Converts modality-specific response labels to modality-neutral response labels so that visual and auditory respones can be easily compared

# %% [markdown]
First, make a legend that converts the modality-specific language of each question and response into modality-neutral language so its easier to compare the modalities on the same graphs and tests

# %% [code]
# This conversion legend was originally an Excel file, but it's been converted to a dictionary here so it doesn't have to be uploaded separately. After the dictionary is made, it gets turned into a DataFrame variable called "legend"
legend_as_dic = {'Qs': {0: 'QX.2',
  1: 'QX.2',
  2: 'QX.5',
  3: 'QX.5',
  4: 'QX.5',
  5: 'QX.7',
  6: 'QX.7',
  7: 'QX.7',
  8: 'QX.7',
  9: 'QX.9',
  10: 'QX.9',
  11: 'QX.9',
  12: 'QX.11',
  13: 'QX.11',
  14: 'QX.11',
  15: 'QX.11',
  16: 'QX.11',
  17: 'QX.13',
  18: 'QX.13',
  19: 'QX.13',
  20: 'QX.13',
  21: 'QX.13',
  22: 'QX.16_1',
  23: 'QX.16_1',
  24: 'QX.18_1',
  25: 'QX.18_1',
  26: 'QX.21',
  27: 'QX.21',
  28: 'QX.21',
  29: 'QX.23#1_1',
  30: 'QX.23#1_1',
  31: 'QX.23#1_1',
  32: 'QX.23#1_2',
  33: 'QX.23#1_2',
  34: 'QX.23#1_2',
  35: 'QX.23#1_3',
  36: 'QX.23#1_3',
  37: 'QX.23#1_3',
  38: 'QX.23#1_4',
  39: 'QX.23#1_4',
  40: 'QX.23#1_4',
  41: 'QX.23#1_5',
  42: 'QX.23#1_5',
  43: 'QX.23#1_5',
  44: 'QX.23#1_6',
  45: 'QX.23#1_6',
  46: 'QX.23#1_6',
  47: 'QX.23#1_7',
  48: 'QX.23#1_7',
  49: 'QX.23#1_7',
  50: 'QX.23#1_8',
  51: 'QX.23#1_8',
  52: 'QX.23#1_8',
  53: 'QX.23#1_9',
  54: 'QX.23#1_9',
  55: 'QX.23#1_9',
  56: 'QX.23#1_10',
  57: 'QX.23#1_10',
  58: 'QX.23#1_10',
  59: 'QX.23#1_11',
  60: 'QX.23#1_11',
  61: 'QX.23#1_11',
  62: 'QX.23#1_12',
  63: 'QX.23#1_12',
  64: 'QX.23#1_12',
  65: 'QX.23#1_13',
  66: 'QX.23#1_13',
  67: 'QX.23#1_13',
  68: 'QX.24',
  69: 'QX.24',
  70: 'QX.24',
  71: 'QX.28',
  72: 'QX.28',
  73: 'QX.28',
  74: 'QX.30',
  75: 'QX.30',
  76: 'QX.30',
  77: 'QX.30',
  78: 'QX.30',
  79: 'QX.30',
  80: 'QX.37',
  81: 'QX.37',
  82: 'QX.37',
  83: 'QX.39',
  84: 'QX.39',
  85: 'QX.39',
  86: 'QX.39',
  87: 'QX.39',
  88: 'QX.39',
  89: 'QX.45',
  90: 'QX.45',
  91: 'QX.45',
  92: 'QX.45',
  93: 'QX.47',
  94: 'QX.47',
  95: 'QX.47',
  96: 'QX.49',
  97: 'QX.49',
  98: 'QX.49',
  99: 'QX.49',
  100: np.nan,
  101: np.nan,
  102: np.nan},
 'Q4s': {0: 'Q4.2',
  1: 'Q4.2',
  2: 'Q4.5',
  3: 'Q4.5',
  4: 'Q4.5',
  5: 'Q4.7',
  6: 'Q4.7',
  7: 'Q4.7',
  8: 'Q4.7',
  9: 'Q4.9',
  10: 'Q4.9',
  11: 'Q4.9',
  12: 'Q4.11',
  13: 'Q4.11',
  14: 'Q4.11',
  15: 'Q4.11',
  16: 'Q4.11',
  17: 'Q4.13',
  18: 'Q4.13',
  19: 'Q4.13',
  20: 'Q4.13',
  21: 'Q4.13',
  22: 'Q4.16_1',
  23: 'Q4.16_1',
  24: 'Q4.18_1',
  25: 'Q4.18_1',
  26: 'Q4.21',
  27: 'Q4.21',
  28: 'Q4.21',
  29: 'Q4.23#1_1',
  30: 'Q4.23#1_1',
  31: 'Q4.23#1_1',
  32: 'Q4.23#1_2',
  33: 'Q4.23#1_2',
  34: 'Q4.23#1_2',
  35: 'Q4.23#1_3',
  36: 'Q4.23#1_3',
  37: 'Q4.23#1_3',
  38: 'Q4.23#1_4',
  39: 'Q4.23#1_4',
  40: 'Q4.23#1_4',
  41: 'Q4.23#1_5',
  42: 'Q4.23#1_5',
  43: 'Q4.23#1_5',
  44: 'Q4.23#1_6',
  45: 'Q4.23#1_6',
  46: 'Q4.23#1_6',
  47: 'Q4.23#1_7',
  48: 'Q4.23#1_7',
  49: 'Q4.23#1_7',
  50: 'Q4.23#1_8',
  51: 'Q4.23#1_8',
  52: 'Q4.23#1_8',
  53: 'Q4.23#1_9',
  54: 'Q4.23#1_9',
  55: 'Q4.23#1_9',
  56: 'Q4.23#1_10',
  57: 'Q4.23#1_10',
  58: 'Q4.23#1_10',
  59: 'Q4.23#1_11',
  60: 'Q4.23#1_11',
  61: 'Q4.23#1_11',
  62: 'Q4.23#1_12',
  63: 'Q4.23#1_12',
  64: 'Q4.23#1_12',
  65: 'Q4.23#1_13',
  66: 'Q4.23#1_13',
  67: 'Q4.23#1_13',
  68: 'Q4.24',
  69: 'Q4.24',
  70: 'Q4.24',
  71: 'Q4.28',
  72: 'Q4.28',
  73: 'Q4.28',
  74: 'Q4.30',
  75: 'Q4.30',
  76: 'Q4.30',
  77: 'Q4.30',
  78: 'Q4.30',
  79: 'Q4.30',
  80: 'Q4.37',
  81: 'Q4.37',
  82: 'Q4.37',
  83: 'Q4.39',
  84: 'Q4.39',
  85: 'Q4.39',
  86: 'Q4.39',
  87: 'Q4.39',
  88: 'Q4.39',
  89: 'Q4.45',
  90: 'Q4.45',
  91: 'Q4.45',
  92: 'Q4.45',
  93: 'Q4.47',
  94: 'Q4.47',
  95: 'Q4.47',
  96: 'Q4.49',
  97: 'Q4.49',
  98: 'Q4.49',
  99: 'Q4.49',
  100: 'Q4.52',
  101: 'Q4.52',
  102: 'Q4.52'},
 'Q5s': {0: 'Q5.2',
  1: 'Q5.2',
  2: 'Q5.5',
  3: 'Q5.5',
  4: 'Q5.5',
  5: 'Q5.7',
  6: 'Q5.7',
  7: 'Q5.7',
  8: 'Q5.7',
  9: 'Q5.9',
  10: 'Q5.9',
  11: 'Q5.9',
  12: 'Q5.11',
  13: 'Q5.11',
  14: 'Q5.11',
  15: 'Q5.11',
  16: 'Q5.11',
  17: 'Q5.13',
  18: 'Q5.13',
  19: 'Q5.13',
  20: 'Q5.13',
  21: 'Q5.13',
  22: 'Q5.16_1',
  23: 'Q5.16_1',
  24: 'Q5.18_1',
  25: 'Q5.18_1',
  26: 'Q5.21',
  27: 'Q5.21',
  28: 'Q5.21',
  29: 'Q5.23#1_1',
  30: 'Q5.23#1_1',
  31: 'Q5.23#1_1',
  32: 'Q5.23#1_2',
  33: 'Q5.23#1_2',
  34: 'Q5.23#1_2',
  35: 'Q5.23#1_3',
  36: 'Q5.23#1_3',
  37: 'Q5.23#1_3',
  38: 'Q5.23#1_4',
  39: 'Q5.23#1_4',
  40: 'Q5.23#1_4',
  41: 'Q5.23#1_5',
  42: 'Q5.23#1_5',
  43: 'Q5.23#1_5',
  44: 'Q5.23#1_6',
  45: 'Q5.23#1_6',
  46: 'Q5.23#1_6',
  47: 'Q5.23#1_7',
  48: 'Q5.23#1_7',
  49: 'Q5.23#1_7',
  50: 'Q5.23#1_8',
  51: 'Q5.23#1_8',
  52: 'Q5.23#1_8',
  53: 'Q5.23#1_9',
  54: 'Q5.23#1_9',
  55: 'Q5.23#1_9',
  56: 'Q5.23#1_10',
  57: 'Q5.23#1_10',
  58: 'Q5.23#1_10',
  59: 'Q5.23#1_11',
  60: 'Q5.23#1_11',
  61: 'Q5.23#1_11',
  62: 'Q5.23#1_12',
  63: 'Q5.23#1_12',
  64: 'Q5.23#1_12',
  65: 'Q5.23#1_13',
  66: 'Q5.23#1_13',
  67: 'Q5.23#1_13',
  68: 'Q5.24',
  69: 'Q5.24',
  70: 'Q5.24',
  71: 'Q5.28',
  72: 'Q5.28',
  73: 'Q5.28',
  74: 'Q5.30',
  75: 'Q5.30',
  76: 'Q5.30',
  77: 'Q5.30',
  78: 'Q5.30',
  79: 'Q5.30',
  80: 'Q5.37',
  81: 'Q5.37',
  82: 'Q5.37',
  83: 'Q5.39',
  84: 'Q5.39',
  85: 'Q5.39',
  86: 'Q5.39',
  87: 'Q5.39',
  88: 'Q5.39',
  89: 'Q5.45',
  90: 'Q5.45',
  91: 'Q5.45',
  92: 'Q5.45',
  93: 'Q5.47',
  94: 'Q5.47',
  95: 'Q5.47',
  96: 'Q5.49',
  97: 'Q5.49',
  98: 'Q5.49',
  99: 'Q5.49',
  100: np.nan,
  101: np.nan,
  102: np.nan},
 'Vistext': {0: 'Yes',
  1: 'No',
  2: 'No, my thought content was not visible in any way',
  3: "Yes, automatically, even if I wasn't trying to make it do so",
  4: 'Yes, but only because I specifically tried to make it do so',
  5: 'My thought content was never visible in any way',
  6: 'As soon as I started thinking',
  7: 'Within a second or two after I started thinking',
  8: 'Several or more seconds after I started thinking',
  9: 'My thought content was not visible in any way',
  10: 'My thought content fluctuated in how visible it was during the period',
  11: 'My thought content was consistently visible for the entire period',
  12: 'My thought content was not visible in any way',
  13: 'Less than half, but not none',
  14: 'About half the time',
  15: 'More than half, but not all',
  16: 'All of the period',
  17: 'My thought content was not visible in any way',
  18: 'Towards the start of my thought',
  19: 'Towards the middle of my thought',
  20: 'Towards the end of my thought',
  21: 'It was equally visible throughout my thought',
  22: 'Duration (in seconds)',
  23: -99,
  24: 'Duration (in seconds)',
  25: -99,
  26: 'No, regardless of whether I moved or not',
  27: 'No, I had to move some part of myself in some way, even if by just a very small amount',
  28: 'Yes, even if I was completely still and unmoving',
  29: 'Agree',
  30: 'Disagree',
  31: 'I did not see my thought in any way',
  32: 'Agree',
  33: 'Disagree',
  34: 'I did not see my thought in any way',
  35: 'Agree',
  36: 'Disagree',
  37: 'I did not see my thought in any way',
  38: 'Agree',
  39: 'Disagree',
  40: 'I did not see my thought in any way',
  41: 'Agree',
  42: 'Disagree',
  43: 'I did not see my thought in any way',
  44: 'Agree',
  45: 'Disagree',
  46: 'I did not see my thought in any way',
  47: 'Agree',
  48: 'Disagree',
  49: 'I did not see my thought in any way',
  50: 'Agree',
  51: 'Disagree',
  52: 'I did not see my thought in any way',
  53: 'Agree',
  54: 'Disagree',
  55: 'I did not see my thought in any way',
  56: 'Agree',
  57: 'Disagree',
  58: 'I did not see my thought in any way',
  59: 'Agree',
  60: 'Disagree',
  61: 'I did not see my thought in any way',
  62: 'Agree',
  63: 'Disagree',
  64: 'I did not see my thought in any way',
  65: 'Agree',
  66: 'Disagree',
  67: 'I did not see my thought in any way',
  68: 'My thought content was not visible in any way',
  69: 'No',
  70: 'Yes (please specify)',
  71: 'No, my thought content was not visible in any way',
  72: 'No, my thought content was visible, but I could not choose where it seemed to be located',
  73: 'Yes, I could choose where it seemed to be located',
  74: 'My thought content was not visible in any way',
  75: 'It had no consistent location',
  76: 'In my head',
  77: 'In my body',
  78: 'In the environment around me',
  79: 'Not in my head, body, or the environment, but somewhere else',
  80: 'No, my thought content was not visible in any way',
  81: 'No, my thought was visible, but I could not choose where it seemed to be located',
  82: 'Yes, I could choose where it seemed to be located',
  83: 'My thought content was not visible in any way',
  84: 'It had no consistent location',
  85: 'In my head',
  86: 'In my body',
  87: 'In the environment around me',
  88: 'Not in my head, body, or the environment, but somewhere else',
  89: 'It is normal for other people, and normal for me',
  90: 'It is normal for other people, but not normal for me',
  91: 'It is not normal for other people, but is normal for me',
  92: 'It is not normal for other people, and not normal for me',
  93: 'More visible',
  94: 'Same as the average person',
  95: 'Less visible',
  96: 'My thought content is not visible in any way',
  97: 'Hallucination',
  98: 'Mental imagery',
  99: 'Other (please specify)',
  100: 'My thought content was not visible in any way',
  101: 'When my thought content was visible, only some parts of it were visible at the same time',
  102: 'When my thought content was visible, all parts of it were visible at the same time'},
 'Audtext': {0: 'Yes',
  1: 'No',
  2: 'No, my thought content was not audible in any way',
  3: "Yes, automatically, even if I wasn't trying to make it do so",
  4: 'Yes, but only because I specifically tried to make it do so',
  5: 'My thought content was never audible in any way',
  6: 'As soon as I started thinking',
  7: 'Within a second or two after I started thinking',
  8: 'Several or more seconds after I started thinking',
  9: 'My thought content was not audible in any way',
  10: 'My thought content fluctuated in how audible it was during the period',
  11: 'My thought content was consistently audible for the entire period',
  12: 'My thought content was not audible in any way',
  13: 'Less than half, but not none',
  14: 'About half the time',
  15: 'More than half, but not all',
  16: 'All of the period',
  17: 'My thought content was not audible in any way',
  18: 'Towards the start of my thought',
  19: 'Towards the middle of my thought',
  20: 'Towards the end of my thought',
  21: 'It was equally audible throughout my thought',
  22: 'Duration (in seconds)',
  23: -99,
  24: 'Duration (in seconds)',
  25: -99,
  26: 'No, regardless of whether I moved or not',
  27: 'No, I had to move some part of myself in some way, even if by just a very small amount',
  28: 'Yes, even if I was completely still and unmoving',
  29: 'Agree',
  30: 'Disagree',
  31: 'My thought content was not audible in any way',
  32: 'Agree',
  33: 'Disagree',
  34: 'My thought content was not audible in any way',
  35: 'Agree',
  36: 'Disagree',
  37: 'My thought content was not audible in any way',
  38: 'Agree',
  39: 'Disagree',
  40: 'My thought content was not audible in any way',
  41: 'Agree',
  42: 'Disagree',
  43: 'My thought content was not audible in any way',
  44: 'Agree',
  45: 'Disagree',
  46: 'My thought content was not audible in any way',
  47: 'Agree',
  48: 'Disagree',
  49: 'My thought content was not audible in any way',
  50: 'Agree',
  51: 'Disagree',
  52: 'My thought content was not audible in any way',
  53: 'Agree',
  54: 'Disagree',
  55: 'My thought content was not audible in any way',
  56: 'Agree',
  57: 'Disagree',
  58: 'My thought content was not audible in any way',
  59: 'Agree',
  60: 'Disagree',
  61: 'My thought content was not audible in any way',
  62: 'Agree',
  63: 'Disagree',
  64: 'My thought content was not audible in any way',
  65: 'Agree',
  66: 'Disagree',
  67: 'My thought content was not audible in any way',
  68: 'My thought content was not audible in any way',
  69: 'No',
  70: 'Yes (please specify)',
  71: 'No, my thought content was not audible in any way',
  72: 'No, my thought content was audible, but I could not choose where it seemed to be located',
  73: 'Yes, I could choose where it seemed to be located',
  74: 'My thought content was not audible in any way',
  75: 'It had no consistent location',
  76: 'In my head',
  77: 'In my body',
  78: 'In the environment around me',
  79: 'Not in my head, body, or the environment, but somewhere else',
  80: 'No, my thought content was not audible in any way',
  81: 'No, my thought was audible, but I could not choose where it seemed to be located',
  82: 'Yes, I could choose where it seemed to be located',
  83: 'My thought content was not audible in any way',
  84: 'It had no consistent location',
  85: 'In my head',
  86: 'In my body',
  87: 'In the environment around me',
  88: 'Not in my head, body, or the environment, but somewhere else',
  89: 'It is normal for other people, and normal for me',
  90: 'It is normal for other people, but not normal for me',
  91: 'It is not normal for other people, but is normal for me',
  92: 'It is not normal for other people, and not normal for me',
  93: 'More audible',
  94: 'Same as the average person',
  95: 'Less audible',
  96: 'My thought content is not audible in any way',
  97: 'Hallucination',
  98: 'Mental imagery',
  99: 'Other (please specify)',
  100: np.nan,
  101: np.nan,
  102: np.nan},
 'Plottext': {0: 'Yes',
  1: 'No exp.',
  2: 'No sensory experience',
  3: 'Automatically',
  4: 'Intentionally',
  5: 'No sensory experience',
  6: 'Immediately',
  7: '<=2 seconds',
  8: '>2 seconds',
  9: 'No sensory experience',
  10: 'Fluctuating',
  11: 'Consistent',
  12: 'No sensory experience',
  13: 'Less than half, but not none',
  14: 'About half the time',
  15: 'More than half, but not all',
  16: 'All of the period',
  17: 'No sensory experience',
  18: 'Start',
  19: 'Middle',
  20: 'End',
  21: 'Equal throughout',
  22: 'Duration (in seconds)',
  23: 'No sensory experience',
  24: 'Duration (in seconds)',
  25: 'No sensory experience',
  26: 'No sensory experience',
  27: 'Movement required',
  28: 'Movement not required',
  29: 'Agree',
  30: 'Disagree',
  31: 'No exp.',
  32: 'Agree',
  33: 'Disagree',
  34: 'No exp.',
  35: 'Agree',
  36: 'Disagree',
  37: 'No exp.',
  38: 'Agree',
  39: 'Disagree',
  40: 'No exp.',
  41: 'Agree',
  42: 'Disagree',
  43: 'No exp.',
  44: 'Agree',
  45: 'Disagree',
  46: 'No exp.',
  47: 'Agree',
  48: 'Disagree',
  49: 'No exp.',
  50: 'Agree',
  51: 'Disagree',
  52: 'No exp.',
  53: 'Agree',
  54: 'Disagree',
  55: 'No exp.',
  56: 'Agree',
  57: 'Disagree',
  58: 'No exp.',
  59: 'Agree',
  60: 'Disagree',
  61: 'No exp.',
  62: 'Agree',
  63: 'Disagree',
  64: 'No exp.',
  65: 'Agree',
  66: 'Disagree',
  67: 'No exp.',
  68: 'No sensory experience',
  69: 'No',
  70: 'Yes (please specify)',
  71: 'No sensory experience',
  72: 'No location control',
  73: 'Location control',
  74: 'No sensory experience',
  75: 'Inconsistent',
  76: 'Head',
  77: 'Body',
  78: 'Environment',
  79: 'Somewhere else',
  80: 'No sensory experience',
  81: 'No location control',
  82: 'Location control',
  83: 'No sensory experience',
  84: 'Inconsistent',
  85: 'Head',
  86: 'Body',
  87: 'Environment',
  88: 'Somewhere else',
  89: 'Normal for others, normal for me',
  90: 'Normal for others, abnormal for me',
  91: 'Abnormal for others, normal for me',
  92: 'Abnormal for others, abnormal for me',
  93: 'More than average',
  94: 'Average',
  95: 'Less than average',
  96: 'No sensory experience',
  97: 'Hallucination',
  98: 'Mental imagery',
  99: 'Other',
  100: 'No visual experience',
  101: 'Some parts visible',
  102: 'All parts visible'}}

legend = pd.DataFrame(legend_as_dic)

# %% [markdown]
Then, do the conversion

# %% [code]
# Store original unneutralised dataframes in case needed later
not_neutral_filtered = filtered.copy()
not_neutral_prehall_filtered = prehall_filtered.copy()

# Function for converting numeric strings to actual floats
def floatStrConv(string):
    try:
        return float(string)
    except:
        return string

# Neutralise most vis-aud questions
for qmod in ['Q4s','Q5s']:
    for q in set(legend[qmod]):
        # Make sure q is not nan (only occurs for where there is a visual, but not audio, question in the legend)
        if type(q) == type(''):
            if '4' in qmod: modtext = 'Vistext'
            if '5' in qmod: modtext = 'Audtext'
            legendqdf = legend[legend[qmod] == q]

            # Make a dictionary mapping original response labels to neutral ones, for each question
            conversiondf = legendqdf.set_index(modtext)['Plottext']
            conversiondic = conversiondf.to_dict()
            if '.18' in q or '.16' in q:
                conversiondic['-99'] = conversiondic[-99]

            # Convert the question's responses to the neutral labels
            filtered.loc[:,q] = filtered[q].replace(conversiondic)
            prehall_filtered.loc[:,q] = prehall_filtered[q].replace(conversiondic)
            
            # Make the series categorical (for categorical, not quantitative, questions)
            if '.18' in q or '.16' in q:
                filtered.loc[:,q] = filtered[q].apply(floatStrConv)
            else:
                catinfo = pd.api.types.CategoricalDtype(categories=list(set(conversiondic.values())), ordered=False)
                filtered.loc[:,q] = filtered[q].astype(catinfo)
                prehall_filtered.loc[:,q] = prehall_filtered[q].astype(catinfo)
            
# Convert other questions
for q in ['Q6.1','Q6.3','Q6.5','Q6.7']:
    if q in ['Q6.1','Q6.3']:
        conversiondic = {'Yes, but only if I specifically try/tried to':'Intentional cross-modal imagery','Yes, automatically':'Automatic cross-modal imagery','No':'No cross-modal sensory experience'}
    elif q in ['Q6.5','Q6.7']:
        conversiondic = {"No, I do not see or hear what I'm thinking about in any way":'No sensory experience',"No, I only see what I'm thinking about in some way":'No cross-modal sensory experience',
             "No, I only hear what I'm thinking about in some way":'No cross-modal sensory experience','Yes, but only if I specifically try/tried to':'Intentional dual-modal imagery','Yes, automatically':'Automatic dual-modal imagery'}
    filtered.loc[:,q] = filtered[q].replace(conversiondic)
    prehall_filtered.loc[:,q] = prehall_filtered[q].replace(conversiondic)
    catinfo = pd.api.types.CategoricalDtype(categories=list(set(conversiondic.values())), ordered=False)
    filtered.loc[:,q] = filtered[q].astype(catinfo)
    prehall_filtered.loc[:,q] = prehall_filtered[q].astype(catinfo)


# %% [markdown]
## Statistics functions & helper functions

# %% [code]
def report(Data, Exclude=None, Collapse2x2=None):
    """Wrapper function for testSquareTable to output pretty-formatted statistics"""
    output = testSquareTable(Data, exclude=Exclude, collapse2x2=Collapse2x2, verbose=False)
    df, p, stat = output[1].df, output[1].pvalue, output[1].statistic
    stat = round(stat,1)
    p = round(p,3)
    if p < 0.001:
        ptext = 'p < .001'
    elif p > 0.9995:
        ptext = 'p > .999'
    else:
        ptext = 'p = ' + f'{p:.3f}'[1:]
    return 'Χ\u00B2'+f'({df}) = {stat}, {ptext}'

def reportWilcoxon(Data1, Data2=None, Method='approx'):
    "Wrapper function for wilcoxon function to output pretty-formatted statistics"
    if type(Data2)==type(None):
        stats = wilcoxon(Data1, method=Method)
    else:
        stats = wilcoxon(Data1, Data2, method=Method)
    p = stats.pvalue
    if p < 0.001:
        ptext = 'p < .001'
    elif p > 0.9995:
        ptext = 'p > .999'
    else:
        ptext = 'p = ' + f'{p:.3f}'[1:]
    return f'Z = {stats.zstatistic:.1f}, {ptext}'

def reportLevene(Data1, Data2, Method='approx'):
    "Wrapper function for levene function to output pretty-formatted statistics"
    stats = levene(Data1, Data2)
    p = stats.pvalue
    if p < 0.001:
        ptext = 'p < .001'
    elif p > 0.9995:
        ptext = 'p = 1.000'
    else:
        ptext = 'p = ' + f'{p:.3f}'[1:]
    return f'W = {stats.statistic:.1f}, {ptext}'

def getMultimodals(df, exclude=['No sensory experience','No exp.','No visual experience','No cross-modal sensory experience']):
    """Returns a dataset which only includes people who experienced imagery in box modalities."""
    # For the dual-modal question, NSE decides whether they are multimodal, but for the cross-modal question, NCMSE decides
    existingcategories = list(df.iloc[:,0].cat.categories)
    if "No cross-modal sensory experience" in existingcategories and "No sensory experience" in existingcategories:
        exclude = exclude[:-1]
    
    columns = df.columns
    df = df[~df.isin(exclude).any(axis=1)].copy(deep=True) # Need to do the copy to avoid SettingWithCopyWarning
    for column in columns:
        for category in exclude:
            if category in list(df.loc[:,column].cat.categories):
                df.loc[:,column] = df[column].cat.remove_categories(category)
    return df

def getUnimodals(df, include=['No sensory experience','No exp.','No visual experience','No cross-modal sensory experience']):
    """Returns a dataset which only includes people who experienced imagery in a single modality"""
    # For the dual-modal question, NSE decides whether they are multimodal, but for the cross-modal question, NCMSE decides
    existingcategories = list(df.iloc[:,0].cat.categories)
    if "No cross-modal sensory experience" in existingcategories and "No sensory experience" in existingcategories:
        include = include[:-1]
    
    columns = df.columns
    df = df[df.isin(include).any(axis=1)].copy(deep=True) # Need to do the copy to avoid SettingWithCopyWarning
    return df

def countPair(df):
    """Returns the proportion of each response option for each modality in a single dataframe"""
    df = round(100*pd.concat((df.iloc[:,0].value_counts(normalize=True), df.iloc[:,1].value_counts(normalize=True)), axis=1),1)
    return df

def testSquareTable(df, exclude=None, testtype='homogeneity', collapse2x2=None, verbose=False):
    
    """Input a CATEGORICAL dataframe with modality-neutral response text
    (i.e. a dataframe returned from neutraliseResponseLabels). Prints
    a square table of counts, and also the p-value for a homogeneity test
    and a symmetry test.
    
    Symmetry test tells, for example, if the proportion of people who said they
    had no visual imagery and did have good auditory was the same as the proportion
    who said they had no auditory imagery and did have good visual. Homogeneity tells 
    whether the marginals are the same, e.g. whether the proportion of people who 
    said they have no visual imagery is the same as the proportion that said they have
    no auditory imagery. Homogeneity should be the correct analysis for my uses here, 
    I think.
    
    Respondents can also be excluded if they returned a particular response(s) for just
    ONE of the columns (modalities), e.g. 'No sensory experience'. This lets you detect
    differences in the distributions that aren't from changes to that category alone
    while still allowing the data to be analysed as paired. Input string or list 
    of strings.
    
    We can also collapse to a 2x2 table, such that the proportion of a given 
    response choice can be compared against how many responded with any responses
    which were not that one. Provide a response label to collapse2x2 to make a 
    2x2 centred around this response option."""
    
    if type(collapse2x2)==type('string'):
        collapsetext = f'Collapsed to 2x2: {collapse2x2} vs not-{collapse2x2}, '
    else:
        collapsetext = 'Distribution-wide, '
    
    # Allows handling of lists of categories to exclude as well as single strings
    if not exclude:
        exclude = []
    if type(exclude)==type('string'):
        exclude = [exclude]
        
    # Remove rows that contain the excluded response category
    columns = df.columns
    df = df[~df.isin(exclude).any(axis=1)].copy(deep=True) # Need to do the copy to avoid SettingWithCopyWarning
    for column in columns:
        for category in exclude:
            if category in list(df.loc[:,column].cat.categories):
                df.loc[:,column] = df[column].cat.remove_categories(category)
    if exclude:
        excludetext = f'Excluding if reported "{exclude}" at least once (n={len(df)})...'
    else:
        excludetext = f'No exclusions (n={len(df)})...'
        
    # Collapse to a 2x2 for direct comparisons if needed
    if type(collapse2x2)==type('string'):
        catinfo = pd.api.types.CategoricalDtype(categories=[collapse2x2,'Others'], ordered=False)
        df = df.astype(catinfo)
        df = df.mask(df != collapse2x2,'Others')
    
    # Do stats
    table = SquareTable.from_data(df, shift_zeros=False)
    homo_stuart_maxwell = table.homogeneity(method='stuart_maxwell')
    homo_bhapkar = table.homogeneity(method='bhapkar')
    sym = table.symmetry()
    
    if verbose:
        if homo_stuart_maxwell.pvalue < 0.0001:
            homo_stuart_maxwellp = '<0.0001'
        else:
            homo_stuart_maxwellp = round(homo_stuart_maxwell.pvalue,3)
        if homo_bhapkar.pvalue < 0.0001:
            homo_bhapkarp = '<0.0001'
        else:
            homo_bhapkarp = round(homo_bhapkar.pvalue,3)
        if sym.pvalue < 0.0001:
            symp = '<0.0001'
        else:
            symp = round(sym.pvalue,3)

        if testtype == 'homogeneity' or testtype == 'both':
            #print(f'{excludetext}...Homogeneity (Stuart-Maxwell) p-value: {homo_stuart_maxwell}')
            print(f'{collapsetext}{excludetext}...Homogeneity (Bhapkar) p-value: {homo_bhapkarp}')
        if testtype == 'symmetry' or testtype == 'both':
            print(f'{collapsetext}{excludetext}...Symmetry p-value: {symp}')
        
    """For some reason, the table.table_orig will incorrectly duplicate the x-axis label 
    for the categories (e.g "auditory") onto the y-axis (such that it also reads "auditory" 
    instead of "visual") if one of the rows or columns contains all zeros. Consequently,
    we return the pandas crosstab of the data rather than table.table_orig"""
    crosstab = pd.crosstab(df.iloc[:,0],df.iloc[:,1], dropna=False)
    return crosstab, homo_bhapkar, homo_stuart_maxwell, sym


# %% [markdown]
# Prevalence of mental imagery

# %% [code]
# Code to find how often people reported no imagery in a given modality across the questionnaire
data = filtered.copy()
labeldic = {'2': 'presence', 
            '5': 'automaticity', 
            '7': 'onset latency (cat)', 
            '9': 'consistency',
            '11': 'proportion', 
            '13': 'peak moment',
            '16_1': 'onset latency (num)', 
            '18_1': 'duration (num)', 
            '21': 'movement req.', 
            '23#1_1': 'req. head', 
            '23#1_2': 'req. brow', 
            '23#1_3': 'req. eyelids',
            '23#1_4': 'req. eyes', 
            '23#1_5': 'req. ears', 
            '23#1_6': 'req. cheeks', 
            '23#1_7': 'req. nose', 
            '23#1_8': 'req. mouth', 
            '23#1_9': 'req. tongue', 
            '23#1_10': 'req. throat',
            '23#1_11': 'req. arms/hands', 
            '23#1_12': 'req. torso', 
            '23#1_13': 'req. legs/feet', 
            '24': 'req. other', 
            '28': 'loc. control open', 
            '30': 'loc. (cat) open', 
            #'32': 'loc. (num) sideview open', #these questions were not shown if they did not report imagery earlier, so they are excluded here
            #'34': 'loc. (num) topview open', 
            '37': 'loc. control closed', 
            '39': 'loc. (cat) closed',
            #'41': 'loc. (num) sideview closed', 
            #'43': 'loc. (num) topview closed', 
            '45': 'normality', 
            '47': 'averageness',
            '49': 'terminology'}
visqs = ['Q6.7']
audqs = ['Q6.5']
totalqs = len(labeldic.keys())
for qi in labeldic.keys():
    visqs.append(f'Q4.{qi}')
    audqs.append(f'Q5.{qi}')
    
# Gets a tally of the number of times NSE was reported once, the number of times it was reported twice, etc...
# Visual
vistally = pd.concat((data[visqs].isin(['No sensory experience','No exp.','No visual experience']), data['Q6.3'].isin(['No cross-modal sensory experience'])), axis=1)
vistally = vistally.sum(axis=1)
vismean = round(vistally.mean(),1)
vistally = round(100*vistally.value_counts(normalize=True),1).sort_index()

# Auditory
audtally = pd.concat((data[audqs].isin(['No sensory experience','No exp.','No auditory experience']), data['Q6.1'].isin(['No cross-modal sensory experience'])), axis=1)
audtally = audtally.sum(axis=1)
audmean = round(audtally.mean(),1)
audtally = round(100*audtally.value_counts(normalize=True),1).sort_index()

# Make sure tallys are present for 0 instances of NSE and the max number of instances of NSE
for tallydataset in [vistally,audtally]:
    if not 0 in tallydataset.index:  tallydataset[0] = 0
    if not 1 in tallydataset.index:  tallydataset[1] = 0
    if not totalqs in tallydataset.index:  tallydataset[totalqs] = 0
    
# Explicitly-reported NSE props
explicits = filtered[['Q4.2', 'Q5.2']]
explicits.columns = ['visual','auditory']
explicits = countPair(explicits)

# %% [code]
text = (f'We first sought to determine the proportion of individuals who experienced mental imagery at all in either the visual or auditory modality. We asked respondents to think about the appearance or sound of the letter "O" and report whether or not they in some way saw or heard their thought content during this specific instance of thinking. When asked explicitly, {explicits.loc["Yes","visual"]}% of included respondents reported a visual experience of their thought content and {explicits.loc["Yes","auditory"]}% reported an auditory experience of their thought content.' + '<br>' +

f'However, there was considerable variation in how well individuals were able to generate mental imagery over the course of the questionnaire. During the questionnaire, respondents were frequently asked to attempt imagery and report only on their experience at that moment. Given that there were 30 questions probing the properties of imagery per visual or auditory modality, participants were therefore presented with 30 opportunities to report whether they experienced imagery at the time of answering. Across these questions, {vistally.iloc[1:-1].sum()}% of participants reported no visual experience of their thought content at least once, while {vistally[totalqs]}% never reported a visual experience of their thought content. For audition, {round(audtally.iloc[1:-1].sum(),1)}% reported no auditory experience of their thought content at least once and {audtally[totalqs]}% never reported an auditory experience of their thought content. On average, participants reported no sensory experience on {vismean}% of visual questions, and {audmean}% of auditory questions. Overall, this variability reflects that mental imagery generation may be far from reliable for many individuals.')
md(text)

# %% [markdown]
# Eyes-open vs. eyes-closed mental imagery

# %% [markdown]
Q4.28, Q5.28, Q4.37, Q5.37; Q4.30, Q4.39, Q5.30, Q5.39

# %% [code]
vis1, vis2 = filtered[['Q4.28', 'Q4.37']], filtered[['Q4.30', 'Q4.39']]
aud1, aud2 = filtered[['Q5.28', 'Q5.37']], filtered[['Q5.30', 'Q5.39']]
for data in [vis1, vis2, aud1, aud2]: data.columns = ['open','closed']

# %% [code]
text = f"We investigated whether the incidence of imagery varied depending on whether a participant conducted imagery with their eyes open or closed. Our questionnaire included two opportunities for this comparison to be made per modality, with the average incidences across these two comparisons shown in Figure 1A. In the visual modality, more respondents reported imagery with their eyes closed rather than open (comparison 1: {report(vis1, Collapse2x2='No sensory experience')}; comparison 2: {report(vis2, Collapse2x2='No sensory experience')}). Yet, eye-closure did not affect the likelihood of hearing one's thought content (comparison 1: {report(aud1, Collapse2x2='No sensory experience')}; comparison 2: {report(aud2, Collapse2x2='No sensory experience')})."
md(text)

# %% [markdown]
# Intentionality: Did mental imagery occur automatically?

# %% [markdown]
Q4.5, Q5.5

# %% [code]
qnum = 'QX.5'
data = filtered[['Q4.' + qnum.split('.')[-1], 'Q5.' + qnum.split('.')[-1]]]
data.columns = ['visual','auditory']
props = countPair(getUnimodals(data))

# %% [code]
text=(f"We probed whether a sensory experience occurred intentionally or automatically as part of thinking. For those reporting an imagery experience in both modalities (i.e. multimodal imagers; Figure 1B), there was no significant difference in whether imagery occurred automatically or intentionally between modalities ({report(getMultimodals(data))}). However, for those reporting only either visual imagery or auditory imagery (unimodal imagers, n = {len(getUnimodals(data))}), responding differed between modalities ({report(getUnimodals(data))}), with visual imagery {round(props.loc['Intentionally','visual']/props.loc['Intentionally','auditory'],1)}x more frequently reported as intentionally-constructed compared to auditory imagery ({props.loc['Intentionally','visual']}% vs. {props.loc['Intentionally','auditory']}% respectively; {report(getUnimodals(data), Collapse2x2='Intentionally')}). Yet, there was no significant difference between modalities in the frequency with which unimodal imagers reported visual or auditory imagery to be automatic when thinking ({props.loc['Automatically','visual']}% vs. {props.loc['Automatically','auditory']}% respectively; {report(getUnimodals(data), Collapse2x2='Automatically')}). Hence, for those who experienced imagery in both modalities, imagery was just as likely to be generated in the same way in either modality. Yet, if visual imagery was the only form of imagery experienced, it was most likely to be intentionally constructed, rather than automatically. If only auditory imagery was experienced, it was most frequently automatic. However, differences between modalities for unimodal imagers may be at least partially attributed to differences in the proportion of individuals who failed to experience visual and auditory imagery at all for this question ({props.loc['No sensory experience','visual']}% vs. {props.loc['No sensory experience','auditory']}% for each modality, respectively; {report(getUnimodals(data), Collapse2x2='No sensory experience')}).")
md(text)

# %% [markdown]
# Consistency: Did mental imagery fluctuate over time?

# %% [markdown]
Q4.9, Q5.9

# %% [code]
qnum = 'QX.9'
data = filtered[['Q4.' + qnum.split('.')[-1], 'Q5.' + qnum.split('.')[-1]]]
data.columns = ['visual','auditory']

# %% [code]
text=f"Participants also reported on whether their thought content was consistently visible or audible during a 5-second period, or whether it fluctuated in how visible or audible it was (Figure 1C). Responding differed between modalities, with imagined sounds more frequently reported as consistent and imagined images more often as fluctuating ({report(getMultimodals(data))})."
md(text)

# %% [markdown]
# Cross-modal imagery: Did real perception in one modality induce mental imagery in another?

# %% [markdown]
Q6.3, Q6.1

# %% [code]
data = filtered[['Q6.3','Q6.1']]
data.columns = ['visual','auditory']
propsmulti = countPair(getMultimodals(data))
propsall = countPair(data)

# %% [code]
text=f"We asked participants whether the real perception of environmental stimuli using vision or audition (i.e. reading written words or listening to spoken words) would induce mental imagery of the same thought content in the other modality (i.e. mentally hearing or seeing the words). We refer to this experience as cross-modal mental imagery. Across all respondents, there were significantly more participants who reported that reading induced auditory imagery (cross-modal auditory imagery) than there were reporting that listening induced visual imagery (cross-modal visual imagery; {report(data, Collapse2x2='No cross-modal sensory experience')}). The distribution of responses regarding cross-modal imagery for all respondents is shown in Figure 1D. To determine whether one form of cross-modal imagery was more or less automatic than the other, we assessed responses only from participants who experienced cross-modal mental imagery in both modalities (i.e. multimodal imagers). For these respondents, cross-modal auditory imagery was far more likely to be automatic than cross-modal visual imagery, with {round(propsmulti.loc['Automatic cross-modal imagery','auditory']/propsmulti.loc['Automatic cross-modal imagery','visual'],1)}x more respondents reporting that they heard words that they were reading automatically compared to automatically seeing words that they were hearing ({report(getMultimodals(data))}). Participants could picture words that they had heard, but generally only if they intended to do so."
md(text)

# %% [markdown]
# Dual-modal imagery: Did mental imagery in one modality induce mental imagery in another?

# %% [markdown]
Q6.7, Q6.5

# %% [code]
data = filtered[['Q6.7','Q6.5']]
data.columns = ['visual','auditory']
props = countPair(getMultimodals(data))

# %% [code]
text=f"We asked participants whether visual or auditory mental imagery in one modality also induced mental imagery in the other modality at the same time. We call this experience dual-modal mental imagery. For respondents who could generate imagery in both modalities separately, imagining in another modality simultaneously was also possible, if not automatic (Figure 1E). Responding did not significantly differ depending on whether auditory imagery was added to visual imagery or whether visual imagery was added to auditory imagery, both for participants who could construct imagery in both modalities separately ({report(getMultimodals(data))}), or only in one of the two modalities separately ({report(getUnimodals(data))})."
md(text)

# %% [markdown]
# Wholeness: Were all parts of a mental image visible at once? (visual only)

# %% [markdown]
Q4.52

# %% [code]
qnum = 'Q4.52'
data = filtered['Q4.52']
props = 100*data.value_counts(normalize=True)

# %% [code]
text=f"We also asked respondents to report whether all parts of their thought content were visible at any given time, or if only some parts were visible (Figure 1F). Partial and whole imagery were reported roughly equally, with slightly more respondents reporting only partial visibility. We did not ask an equivalent question regarding the imagined auditory stimulus as it involved only a single phoneme."
md(text)

# %% [markdown]
## **Figure 1**
Incidence and properties of visual and auditory mental imagery

# %% [code]
# Get datasets to plot
data = filtered.copy()

# Process eyes-open and eyes-closed data from 2 pairs of questions
qdic = {'visual, eyes open':['Q4.28','Q4.30'],'visual, eyes closed':['Q4.37','Q4.39'],'auditory, eyes open':['Q5.28','Q5.30'],'auditory, eyes closed':['Q5.37','Q5.39']}
eyesdf = pd.DataFrame(columns = list(qdic.keys()))
for modality in ['visual','auditory']:
    for eyestate in ['eyes open','eyes closed']:
        df = data[qdic[f'{modality}, {eyestate}']]
        df = df.stack()
        df = df.where(df == 'No sensory experience',other='Sensory experience')
        eyesdf[f'{modality}, {eyestate}'] = df.values

# Make figure
fig, axes = plt.subplots(nrows=3, ncols=2, dpi=600, figsize=(6,6))
c = 0
colours = ['cornflowerblue','orchid']
letters = 'ABCDEF'
axlabelsize = 6
axticklabelsize = 6
titlesize = 10
allow_swapping_to_unimodal_plots = False
for axrow in [0,1,2]:
    for axcol in [0,1]:
        if c == 0: 
            thisdf = eyesdf
            title = 'Eyes closed vs. open'
            subtitle = 'Mean incidence of imagery depending on eye state'
            subjectcount = len(data)
            order = ['sensory experience','no sensory experience']
            qdatas = [data[['Q4.28','Q4.37','Q5.28','Q5.37']], data[['Q4.30','Q4.39','Q5.30','Q5.39']]]
            xlabel = 'occurrence of imagery'
            for i in [0,1]:
                qdata = qdatas[i]
                qdata.columns = ['visual, eyes open','visual, eyes closed','auditory, eyes open','auditory, eyes closed']
                qdata = qdata.apply(lambda x: x.cat.add_categories('Sensory experience'))
                qdata = qdata.where(qdata == 'No sensory experience',other='Sensory experience')
                qdatas[i] = qdata
        elif c == 1:
            qdata = data[['Q4.5','Q5.5']]
            title = 'Intentionality'
            subtitle = 'Did imagery occur automatically?'
            order = ['intentionally','automatically']
            columns = ['visual','auditory']
            xlabel = 'intentionality of imagery'
        elif c == 2:
            qdata = data[['Q4.9','Q5.9']]
            title = 'Consistency'
            subtitle = 'Did imagery fluctuate over time?'
            order = ['fluctuating','consistent']
            columns = ['visual','auditory']
            xlabel = 'consistency of imagery'
        elif c == 3:
            qdata = data[['Q6.3','Q6.1']]
            title = 'Cross-modal imagery'
            subtitle = 'Did real perception in one modality induce imagery in another?'
            order = ['intentional cross-modal imagery','automatic cross-modal imagery', 'no cross-modal sensory experience']
            columns = ['visual imagery \nwhile listening','auditory imagery \nwhile reading']
            xlabel = 'incidence of cross-modal imagery'
        elif c == 4:
            qdata = data[['Q6.7','Q6.5']]
            title = 'Dual-modal imagery'
            subtitle = 'Did imagery in one modality induce imagery in another?'
            order = ['intentional dual-modal imagery','automatic dual-modal imagery','no cross-modal sensory experience']
            columns = ["visual imagery \ninduced by \nauditory imagery","auditory imagery \ninduced by \nvisual imagery"]
            xlabel = 'incidence of imagery in an additional modality'
        elif c == 5:
            qdata = data['Q4.52']
            title = 'Wholeness'
            subtitle = 'Were all parts of a mental image visible at once?'
            order = ['some parts visible','all parts visible','no visual experience']
            xlabel = 'portion of imagined image visible'
              
        # Create ax plot (note: indexes must be sorted before plotting to avoid mislabelling bars)
        if c == 0:
            statsdics = []
            for i in [0,1]:
                qdata = qdatas[i]
                # Do stats for each question
                # Eyes-open vs eyes-closed (within a modality), and also vis vs aud (within a particular eye state)
                statsdic = {'compare eyes':{'visual':None,'auditory':None}, 'compare modalities':{'eyes open':None,'eyes closed':None}}
                for comparison in statsdic.keys():
                    for aspect in statsdic[comparison].keys():
                        if comparison == 'compare eyes': pair = qdata[[f'{aspect}, eyes open',f'{aspect}, eyes closed']]
                        if comparison == 'compare modalities': pair = qdata[[f'visual, {aspect}',f'auditory, {aspect}']]
                        ys = [countPair(pair).loc[label.capitalize(),:].max() for label in order] #Two pieces of code below, before the Plot stats, must also be changed if changing this
                        stats2x2 = [testSquareTable(pair, collapse2x2=label.capitalize(), verbose=False)[1].pvalue for label in order]
                        bonfactor = (len(stats2x2) if len(stats2x2) > 2 else 1)
                        annotations = ['**' if p <= (0.05/bonfactor)/50 else '*' if p <= 0.05/bonfactor else 'n.s.' for p in stats2x2]
                        statsdic[comparison][aspect] = (annotations,ys)
                statsdics.append(statsdic)
                
            # If the stats for both questions have the same annotations, then just pick one and plot those
            # Otherwise, show no stats
            for comparison in statsdic.keys():
                for aspect in statsdic[comparison].keys():
                    if statsdics[0][comparison][aspect][0] == statsdics[1][comparison][aspect][0]:
                        statsdicsequal = True
                    else:
                        statsdicsequal = False
                        print('The stats for each question used to plot the eye state figure are not the same, so you cannot just plot one of them for the average anymore. Stats not shown.')
            if statsdicsequal:
                statsdic = statsdics[0]
        
            # Make a df to plot which is an average of the data of both questions
            stackedqdata = pd.concat(qdatas)
            thisdf = pd.concat((countPair(stackedqdata.iloc[:,[0,1]]), countPair(stackedqdata.iloc[:,[2,3]])), axis=1)
            thisdf.index = [text.lower() for text in thisdf.index]
            thisdf = thisdf.loc[order,:]
            thisdf.index = [textwrap.fill(text,11,break_long_words=False) for text in thisdf.index] # Wraps text
            ax = thisdf.plot(kind='bar', color=['cornflowerblue','yellowgreen','orchid','sandybrown'], rot=0, ax=axes[axrow,axcol], width=0.8)
            ax.set_ylim((0,139))
            ax.set_yticks([0, 25, 50, 75, 100])

            # Plot stats
            # Keep track of previously plotted heights for each xtick
            if statsdicsequal:
                yhistorydic = {}
                for xtick in ax.get_xticks():
                    yhistorydic[xtick] = []
                for comparison in statsdic.keys():
                    for aspect in statsdic[comparison].keys():
                        annotations = statsdic[comparison][aspect][0]
                        ys = statsdic[comparison][aspect][1]

                        for i, annotation in enumerate(annotations):
                            x = ax.get_xticks()[i]
                            y, height = ys[i], (ax.get_ylim()[1]/10)
                            width = ax.patches[0].get_width()  

                            for yhistorical in yhistorydic[x]:
                                while abs(y-yhistorical) < 14:
                                    y += 1

                            # X positions
                            if aspect == 'visual':
                                xleft, xright = x-(1.5*width), x-(width/2)
                            elif aspect == 'auditory':
                                xleft, xright = x+(width/2), x+(1.5*width)
                            elif aspect == 'eyes open':
                                xleft, xright = x-(1.5*width), x+(width/2)
                            elif aspect == 'eyes closed':
                                xleft, xright = x-(width/2), x+(1.5*width)

                            # Only plot if significant here to avoid a mess of non-significant annotations
                            if '*' in annotation:
                                yhistorydic[x].append(y)
                                ax.text((xleft+xright)/2, y+height-(axticklabelsize/2), annotation, ha='center', va='bottom', color='black', size=1.5*axticklabelsize)
                                ax.plot([xleft, xright], [y+height, y+height], lw=width, c='black')       

        if c == 5:
            thisdf = round(100*qdata.value_counts(normalize=True),1)
            thisdf.index = [text.lower() for text in thisdf.index]
            thisdf = thisdf.loc[order]
            thisdf.index = [textwrap.fill(text,18,break_long_words=False) for text in thisdf.index] # Wraps text
            thisdf.name = 'visual'
            ax = thisdf.plot(kind='bar', color=colours if c!=5 else colours[0], rot=0, ax=axes[axrow,axcol])
            ax.set_ylim(axes[2,0].get_ylim())
        
        # Stats and plots
        if c not in [0,5]:
            if c == 4:
                filterlist = ['No sensory experience']
            else:
                filterlist = getMultimodals.__defaults__[0]
            if c == 3:
                stats2x2 = [round(testSquareTable(qdata, collapse2x2=label.capitalize(), verbose=False)[1].pvalue,3) for label in order]
            else:
                stats2x2 = [round(testSquareTable(getMultimodals(qdata, exclude=filterlist), collapse2x2=label.capitalize(), verbose=False)[1].pvalue,3) for label in order]
            bonfactor = (len(stats2x2) if len(stats2x2) > 2 else 1)
            annotations = ['**' if p <= (0.05/bonfactor)/50 else '*' if p <= 0.05/bonfactor else 'n.s.' for p in stats2x2]

            # If no significant multimodal 2x2s, show the unimodal data, if allowed
            if len(set(annotations)) == 1 and 'n.s.' in set(annotations) and allow_swapping_to_unimodal_plots:
                order.append('no sensory experience')
                stats2x2 = [testSquareTable(getUnimodals(qdata, include=filterlist), collapse2x2=label.capitalize(), verbose=False)[1].pvalue for label in order]
                bonfactor = (len(stats2x2) if len(stats2x2) > 2 else 1)
                annotations = ['**' if p <= (0.05/bonfactor)/50 else '*' if p <= 0.05/bonfactor else 'n.s.' for p in stats2x2]
                thisdf = countPair(getUnimodals(qdata, include=filterlist))
                subjectcount = len(getUnimodals(qdata, include=filterlist))
            elif c==3:
                thisdf = countPair(qdata)
                subjectcount = len(qdata)
            else:
                thisdf = countPair(getMultimodals(qdata, exclude=filterlist))
                subjectcount = len(getMultimodals(qdata, exclude=filterlist))
                
            # Make actual plot
            thisdf.columns = columns
            thisdf.index = [text.lower() for text in thisdf.index]
            thisdf = thisdf.loc[order,:]
            thisdf.index = [textwrap.fill(text,18,break_long_words=False) for text in thisdf.index] # Wraps text
            ax = thisdf.plot(kind='bar', color=colours if c!=5 else colours[0], rot=0, ax=axes[axrow,axcol])
            
            # Set ylimits (must come before stats because annotations use them)
            if c in [1]:
                ax.set_ylim(axes[0,0].get_ylim())
            if c in [2,3,4]:
                ax.set_ylim((0,129))
                ax.set_yticks([0, 25, 50, 75, 100])
        
            # Put stats on columns
            for i, annotation in enumerate(annotations):
                x = ax.get_xticks()[i]
                width = ax.patches[0].get_width()
                xleft, xright, y, height = x-(width/2), x+(width/2), thisdf.iloc[i,:].max(), (ax.get_ylim()[1]/10)
                ax.plot([xleft, xright], [y+height, y+height], lw=width, c='black')
                if '*' in annotation:
                    ax.text(x, y+height-(axticklabelsize/2), annotation, ha='center', va='bottom', color='black', size=1.5*axticklabelsize)
                else:
                    ax.text(x, y+height+(axticklabelsize/2), annotation, ha='center', va='bottom', color='black', size=axticklabelsize)
 
        # Customise ax plot, separate to creation, for clarity
        if c==0:
            ax.set_xticklabels(thisdf.index, fontsize=axticklabelsize)
            ax.set_xlabel('number of questions where \nrespondent reported no imagery', fontsize=axlabelsize, labelpad=5)
            ax.bar_label(ax.containers[2], label_type='edge', fontsize=axticklabelsize, fmt='%.0f')
            ax.bar_label(ax.containers[3], label_type='edge', fontsize=axticklabelsize, fmt='%.0f')
            ax.legend(fontsize=4.5, frameon=False, labelcolor=['cornflowerblue','yellowgreen','orchid','sandybrown'], handlelength=0, handletextpad=0, bbox_to_anchor=(0,-0.05,1,1))
        elif c==3:
            ax.legend(fontsize=4.5, frameon=False, labelcolor=colours, handlelength=0, handletextpad=0, bbox_to_anchor=(0,-0.15,1,1))
        elif c==4:
            ax.legend(fontsize=4.5, frameon=False, labelcolor=colours, handlelength=0, handletextpad=0, bbox_to_anchor=(0,-0.05,1,1))
        else:
            ax.legend(fontsize=4.5, frameon=False, labelcolor=colours, handlelength=0, handletextpad=0, loc='upper right')
        if c in [1,3,5]:
            ax.yaxis.set_visible(False)
            ax.spines.left.set_visible(False)
        if c not in [5]:
            ax.bar_label(ax.containers[1], label_type='edge', fontsize=axticklabelsize, fmt='%.0f')
        ax.bar_label(ax.containers[0], label_type='edge', fontsize=axticklabelsize, fmt='%.0f')
        ax.set_xlabel(xlabel, fontsize=axlabelsize, labelpad=5)
        ax.set_title(title, y=1.1, x=0.5, size=titlesize, pad=0, fontweight='bold')
        ax.set_xticklabels(thisdf.index, fontsize=axticklabelsize)
        ax.spines.right.set_visible(False)
        ax.spines.top.set_visible(False)
        ax.spines.left.set_bounds(0, 100)
        ax.text(s=letters[c], y=1.1, x=0, size=titlesize, fontweight='bold', transform=ax.transAxes)
        ax.text(s=subtitle, y=1.01, x=0.5, horizontalalignment='center', size=4.5, transform=ax.transAxes)
        ax.text(s=f'n = {subjectcount}', y=0.93, x=0.5, horizontalalignment='center', size=4.5, transform=ax.transAxes)
        ax.tick_params(axis='y', which='major', labelsize=axticklabelsize)
        ax.set_ylabel('% respondents', fontsize=axlabelsize, labelpad=5)
        
        c += 1
        
fig.tight_layout()
fig.savefig('Figure 1.png')

# %% [markdown]
*Note.* **A-F.** Multiple-choice response distributions. Annotations indicate significant differences between modalities in the proportion of times a specific response option was chosen, after Bonferroni corrections: n.s. = p > 0.05, * = 0.001 < p ≤ 0.05, ** = p ≤ 0.001. **A.** For each modality, there were two opportunities within the questionnaire where eyes-open and eyes-closed imagery could be compared.  Proportions shown here are averaged over these two opportunities. Hypothesis tests yielded the same inferences for both comparisons. For clarity, non-significant results are not shown in this plot, however main effects of modality and eye-state were tested in all combinations. **B.** Responses shown only from those who experienced imagery in both visual and auditory modalities (multimodal imagers). **C.** Responses shown from multimodal imagers only. **D.** Responses pooled from unimodal and multimodal imagers. **E**. Responses shown for participants who could at least generate each of visual and auditory imagery separately **F.** Responses shown in the visual modality only.

# %% [markdown]
# Peak: When was mental imagery most sensory within a thought period?

# %% [markdown]
Q4.13, Q5.13

# %% [code]
qnum = 'QX.13'
data = filtered[['Q4.' + qnum.split('.')[-1], 'Q5.' + qnum.split('.')[-1]]]
data.columns = ['visual','auditory']
props = countPair(getMultimodals(data))

# %% [code]
text=f"Participants reported when their thought content was most visible or audible during a given 5-second period where they actively attempted imagery. For multimodal imagers (Figure 2A), the point when imagery was most sensory significantly differed depending on imagery modality ({report(getMultimodals(data))}). Thought content was most frequently reported as being most sensory during the middle of the imagery period for visual imagery, significantly more frequently than for auditory imagery ({report(getMultimodals(data), Collapse2x2='Middle')}). Auditory imagery, however, was most frequently reported as equally audible throughout the imagery period, by {round(props.loc['Equal throughout','auditory']/props.loc['Equal throughout','visual'],1)}x as many respondents as for visual imagery ({report(getMultimodals(data), Collapse2x2='Equal throughout')})."
md(text)

# %% [markdown]
# Proportion: What proportion of a thought period involved mental imagery?

# %% [markdown]
Q4.11, Q5.11

# %% [code]
qnum = 'QX.11'
data = filtered[['Q4.' + qnum.split('.')[-1], 'Q5.' + qnum.split('.')[-1]]]
data.columns = ['visual','auditory']
props = countPair(getMultimodals(data))

# %% [code]
text=f"We asked participants to report the proportion of time in which sensory content was visible or audible during a 5-second period where they actively attempted imagery. For multimodal imagers (Figure 2B), the proportion of the period which involved imagery significantly differed depending on imagery modality ({report(getMultimodals(data))}). Thought content was most frequently reported as visible for more than half, but not all, of the thought period for visual imagery, significantly more frequently than for auditory imagery ({report(getMultimodals(data), Collapse2x2='More than half, but not all')}). Yet, auditory imagery was most frequently reported as occurring throughout the entire thought period, by {round(props.loc['All of the period','auditory']/props.loc['All of the period','visual'],1)}x as many respondents as for visual imagery ({report(getMultimodals(data), Collapse2x2='All of the period')})."
md(text)

# %% [markdown]
# Onset latency: How long did mental imagery take to initiate?

# %% [markdown]
Q4.16, Q5.16

# %% [code]
# Categorical
qnum = 'QX.7'
data = filtered[['Q4.' + qnum.split('.')[-1], 'Q5.' + qnum.split('.')[-1]]]
data.columns = ['visual','auditory']
props = countPair(getMultimodals(data))

# Numerical
qnum = 'QX.16_1'

# Get data
rawtimes = filtered[[qnum.replace('X','4'),qnum.replace('X','5')]]
rawtimes.columns = ['visual','auditory']

# Run a Wilcoxon signed-rank on all the non-NSE data, treating it as ordinal but not continuous
objectdf = rawtimes.copy()
objectdf = objectdf[~objectdf.isin(['No sensory experience']).any(axis=1)]
floatdf = objectdf.astype('float')
reportwilcoxon = reportWilcoxon(floatdf['visual'], floatdf['auditory'])

# %% [code]
text=f"We asked participants to report how long it took for thought content to become visible or audible, both using a multiple choice question, and  using a sliding numerical scale. For categorical responses from multimodal imagers (Figure 2C), the onset latency of imagery significantly differed depending on modality ({report(getMultimodals(data))}). While imagery was most frequently reported as beginning immediately for both modalities, {round(props.loc['Immediately','auditory']/props.loc['Immediately','visual'],1)}x as many respondents reported immediate auditory imagery compared to visual imagery ({report(getMultimodals(data), Collapse2x2='Immediately')}). Using a sliding scale, participants could report their imagery onset latency with a numerical value from 0-10 seconds or otherwise report that their imagery took 10 or more seconds to initiate (Figure 2D). For multimodal imagers, a two-sided Wilcoxon signed-rank test showed a significant difference in onset latency between visual and auditory imagery ({reportwilcoxon}). Onset latencies for auditory imagery (median = {floatdf['auditory'].median():.1f}s) were shorter than for visual imagery (median = {floatdf['visual'].median():.1f}s). Overall, these findings indicate auditory imagery began more quickly than visual imagery."
md(text)

# %% [markdown]
# Duration: How long could mental imagery definitely and continuously persist?

# %% [markdown]
Q4.18, Q5.18

# %% [code]
# Numerical
qnum = 'QX.18_1'

# Get data
rawtimes = filtered[[qnum.replace('X','4'),qnum.replace('X','5')]]
rawtimes.columns = ['visual','auditory']

# Turn into categorical data (NSE, <10, >=10)
stringdf = rawtimes.copy().astype('string')
stringdf = stringdf.replace({'10.0':'>=10'})
categorical = stringdf.mask(((stringdf != 'No sensory experience') & (stringdf != '>=10')),'<10') # Converts all remaining numbers to '<10'
catinfo = pd.api.types.CategoricalDtype(categories=['No sensory experience','<10','>=10'], ordered=True)
categorical = categorical.astype(catinfo)
props = countPair(getMultimodals(categorical))

# Run a Wilcoxon signed-rank on all the non-NSE data, treating it as ordinal but not continuous
objectdf = rawtimes.copy()
objectdf = objectdf[~objectdf.isin(['No sensory experience']).any(axis=1)]
floatdf = objectdf.astype('float')
reportwilcoxon = reportWilcoxon(floatdf['visual'], floatdf['auditory'])

# %% [code]
text=f"We asked participants to report how long they could definitely and continuously see or hear their thought content. Using a sliding scale, participants could report their imagery duration with a numerical value from 0-10 seconds or otherwise report that their imagery could last for 10 or more seconds (Figure 2E). For multimodal imagers, a two-sided Wilcoxon signed-rank test showed no significant difference in imagery duration between modalities ({reportwilcoxon}). However, visual imagery (median = {floatdf['visual'].median():.1f}s) tended not to last as long as auditory imagery (median = {floatdf['auditory'].median():.1f}s), and there were significant differences between modalities in the proportion of multimodal imagers reporting that their imagery could last for 10 or more seconds, as opposed to below 10 seconds ({report(getMultimodals(categorical))}). Indeed, auditory imagery ({props.loc['>=10','auditory']}%) was {round(props.loc['>=10','auditory']/props.loc['>=10','visual'],1)}x as frequently reported as lasting 10 or more seconds than visual imagery ({props.loc['>=10','visual']}%). Overall, these findings suggest that most respondents could generate auditory imagery for a longer period than visual imagery."
md(text)

# %% [markdown]
## **Figure 2**
Temporal properties of visual and auditory mental imagery

# %% [code]
# Get datasets to plot
data = filtered.copy()

# Onset latency
onsetnumtimes = data[['Q4.16_1','Q5.16_1']].copy()
onsetnumtimes.replace({'No sensory experience':-99}, inplace=True)
onsetnum = pd.DataFrame()
for column in onsetnumtimes.columns:
    onsetnum[column] = pd.cut(onsetnumtimes[column].astype(float), bins=[-99,0,1,2,3,4,5,6,7,8,9,10,11], labels=['No sensory experience','<1','1-1.9','2-2.9','3-3.9','4-4.9','5-5.9','6-6.9','7-7.9','8-8.9','9-9.9','>=10'], right=False)
onsetnum = getMultimodals(onsetnum)
onsetnum = round(100*pd.concat((onsetnum['Q4.16_1'].value_counts(normalize=True), onsetnum['Q5.16_1'].value_counts(normalize=True)), axis=1),1)
onsetnum.columns = ['visual','auditory']

# Duration
durationnumtimes = data[['Q4.18_1','Q5.18_1']].copy()
durationnumtimes.replace({'No sensory experience':-99}, inplace=True)
durationnum = pd.DataFrame()
for column in durationnumtimes.columns:
    durationnum[column] = pd.cut(durationnumtimes[column].astype(float), bins=[-99,0,1,2,3,4,5,6,7,8,9,10,11], labels=['No sensory experience','<1','1-1.9','2-2.9','3-3.9','4-4.9','5-5.9','6-6.9','7-7.9','8-8.9','9-9.9','>=10'], right=False)
durationnum = getMultimodals(durationnum)
durationnum = round(100*pd.concat((durationnum['Q4.18_1'].value_counts(normalize=True), durationnum['Q5.18_1'].value_counts(normalize=True)), axis=1),1)
durationnum.columns = ['visual','auditory']

# Make figure
fig = plt.figure(dpi=600, figsize=(6,4))
spec = mpg.GridSpec(ncols=6, nrows=2)
axes = []
axes.append(fig.add_subplot(spec[0,0:2]))
axes.append(fig.add_subplot(spec[0,2:4]))
axes.append(fig.add_subplot(spec[0,4:]))
axes.append(fig.add_subplot(spec[1,0:3]))
axes.append(fig.add_subplot(spec[1,3:]))
colours = ['cornflowerblue','orchid']
letters = 'ABCDEF'
axlabelsize = 6
axticklabelsize = 6
titlesize = 10
allow_swapping_to_unimodal_plots = False
for c in range(5):
        if c == 0:
            qdata = data[['Q4.13','Q5.13']]
            title = 'Peak'
            subtitle = 'When was imagery most sensory \nwithin a thought period?'
            order = ['start','middle','end','equal throughout']
            xlabel = 'time of imagery peak'
        elif c == 1:
            qdata = data[['Q4.11','Q5.11']]
            title = 'Proportion'
            subtitle = 'What proportion of a thought \nperiod involved imagery?'
            order = ['less than half, but not none','about half the time','more than half, but not all','all of the period']
            xlabel = 'proportion of thought \nperiod containing imagery'
        elif c == 2:
            qdata = data[['Q4.7','Q5.7']]
            title = 'Coarse onset \nlatency'
            subtitle = 'How long did imagery\n take to initiate?'
            order = ['immediately','<=2 seconds','>2 seconds']
            xlabel = 'imagery onset latency'
        elif c == 3: 
            qdata = onsetnumtimes
            thisdf = onsetnum
            title = 'Onset latency'
            subtitle = 'How long did imagery \ntake to initiate?'
            order = ['<1','1-1.9','2-2.9','3-3.9','4-4.9','5-5.9','6-6.9','7-7.9','8-8.9','9-9.9','>=10']
            xlabel = 'onset latency of imagery (seconds)'
        elif c == 4:
            qdata = durationnumtimes
            thisdf = durationnum
            title = 'Duration'
            subtitle = 'How long could imagery \ndefinitely and continuously persist?'
            order = ['<1','1-1.9','2-2.9','3-3.9','4-4.9','5-5.9','6-6.9','7-7.9','8-8.9','9-9.9','>=10']
            xlabel = 'duration of imagery (seconds)'
            
        # Create ax plot (note: indexes must be sorted before plotting to avoid mislabelling bars)
        if c in [3,4]:
            multimodalqdata = qdata[~qdata.isin([-99]).any(axis=1)].copy(deep=True)
            thisdf.index = [text.lower() for text in thisdf.index]
            thisdf = thisdf.loc[order,:]
            thisdf.index = [textwrap.fill(text,10,break_long_words=False) for text in thisdf.index] # Wraps text
            ax = thisdf.iloc[:,0].plot(kind='bar', color=colours[0], rot=0, ax=axes[c], width=1)
            ax = (-1*thisdf.iloc[:,1]).plot(kind='bar', color=colours[1], rot=0, ax=axes[c], width=1)
            ylim = 90
            ax.set_ylim((-ylim,ylim))
            visvaluelabels = [int(val) if val > 1 else val if val > 0 else '' for val in ax.containers[0].datavalues]
            audvaluelabels = [int(abs(val)) if val < -1 else abs(val) if val < 0 else '' for val in ax.containers[1].datavalues]
            ax.bar_label(ax.containers[0], labels=visvaluelabels, label_type='edge', fontsize=axticklabelsize, fmt='%.1f')
            ax.bar_label(ax.containers[1], labels=audvaluelabels, label_type='edge', fontsize=axticklabelsize, fmt='%.1f', padding=0.8)
            yticks = np.concatenate((-np.arange(0,ylim,20)[::-1],np.arange(0,ylim,20)))
            yticklabels = np.abs(yticks)
            ax.set_yticks(yticks)
            ax.set_yticklabels(yticklabels)
            ax.set_xticks([tick-0.5 for tick in ax.get_xticks()] + [ax.get_xticks()[-1]+0.5])
            ax.set_xticklabels([0,1,2,3,4,5,6,7,8,9,10,'\u221e'], fontsize=axticklabelsize)
        else:
            stats2x2 = [testSquareTable(getMultimodals(qdata), collapse2x2=label.capitalize(), verbose=False)[1].pvalue for label in order]
            bonfactor = (len(stats2x2) if len(stats2x2) > 2 else 1)
            annotations = ['**' if p <= (0.05/bonfactor)/50 else '*' if p <= 0.05/bonfactor else 'n.s.' for p in stats2x2]
            # If no significant multimodal 2x2s, use the unimodal data, if allowed
            if len(set(annotations)) == 1 and 'n.s.' in set(annotations) and allow_swapping_to_unimodal_plots:
                order.append('no sensory experience')
                stats2x2 = [testSquareTable(getUnimodals(qdata), collapse2x2=label.capitalize(), verbose=False)[1].pvalue for label in order]
                bonfactor = (len(stats2x2) if len(stats2x2) > 2 else 1)
                annotations = ['**' if p <= (0.05/bonfactor)/50 else '*' if p <= 0.05/bonfactor else 'n.s.' for p in stats2x2]
                thisdf = countPair(getUnimodals(qdata))
                subjectcount = len(getUnimodals(qdata))
            else:
                thisdf = countPair(getMultimodals(qdata))
                subjectcount = len(getMultimodals(qdata))
            thisdf.columns = ['visual','auditory']
            thisdf.index = [text.lower() for text in thisdf.index]
            thisdf = thisdf.loc[order,:]
            thisdf.index = [textwrap.fill(text,10,break_long_words=False) for text in thisdf.index] # Wraps text
            ax = thisdf.plot(kind='bar', color=colours, rot=0, ax=axes[c])
            ax.set_ylim((0,100))
            ax.bar_label(ax.containers[0], label_type='edge', fontsize=axticklabelsize, fmt='%.0f')
            ax.bar_label(ax.containers[1], label_type='edge', fontsize=axticklabelsize, fmt='%.0f')
            ax.legend(fontsize=4.5, frameon=False, labelcolor=colours, handlelength=0, handletextpad=0, loc='upper right')
            ax.set_xticklabels(thisdf.index, fontsize=axticklabelsize)
            if c==1:
                ax.set_xticklabels(thisdf.index, fontsize=5)
            
            # Put stats on columns
            for i, annotation in enumerate(annotations):
                x = ax.get_xticks()[i]
                width = ax.patches[0].get_width()
                xleft, xright, y, height = x-(width/2), x+(width/2), thisdf.iloc[i,:].max(), (ax.get_ylim()[1]/10)
                ax.plot([xleft, xright], [y+height, y+height], lw=width, c='black')
                if '*' in annotation:
                    ax.text(x, y+height-(axticklabelsize/2), annotation, ha='center', va='bottom', color='black', size=1.5*axticklabelsize)
                else:
                    ax.text(x, y+height+(axticklabelsize/2), annotation, ha='center', va='bottom', color='black', size=axticklabelsize)
            
        # Customise ax plot, separate to creation, for clarity
        if c == 3:
            ax.legend(fontsize=4.5, frameon=False, labelcolor=colours, handlelength=0, handletextpad=0, loc='upper right')
        if c == 4:
            ax.legend(fontsize=4.5, frameon=False, labelcolor=colours, handlelength=0, handletextpad=0, loc='upper right')
            ax.set_yticklabels(['' for i in ax.get_yticks()])
            ax.set_ylabel('')
            ax.yaxis.set_visible(False)
            ax.spines.left.set_visible(False)
            
        if c in [3,4]:           
            # Calculate median/mean and put it on the plot
            visaverage = multimodalqdata.iloc[:,0].median(axis=0)-0.5 # Have to subtract a half because everything is offset on the plot to allow the columns to be between ticks
            audaverage = multimodalqdata.iloc[:,1].median(axis=0)-0.5
            ax.scatter(y=0, x=visaverage, alpha=1, s=20, marker='X', facecolors='white', edgecolors=colours[0], linewidth=1)#, zorder=zorder+1)
            ax.scatter(y=0, x=audaverage, alpha=1, s=20, marker='X', facecolors='white', edgecolors=colours[1], linewidth=1)#, zorder=zorder+1)
          
        if c in [1,2]:
            ax.yaxis.set_visible(False)
            ax.spines.left.set_visible(False)
        if c in [0,3]:
            ax.set_ylabel('% respondents', fontsize=axlabelsize, labelpad=5)
        if c == 2:
            ax.set_title(title, y=1.2, x=0.5, size=titlesize, pad=0, fontweight='bold', va='top')
            ax.text(s=subtitle, y=.79, x=0.5, horizontalalignment='center', size=4.5, transform=ax.transAxes)
            ax.text(s=f'n = {subjectcount}', y=0.69, x=0.5, horizontalalignment='center', size=4.5, transform=ax.transAxes)
        else:
            ax.set_title(title, y=1.1, x=0.5, size=titlesize, pad=0, fontweight='bold')
            ax.text(s=subtitle, y=.93, x=0.5, horizontalalignment='center', size=4.5, transform=ax.transAxes)
            ax.text(s=f'n = {subjectcount}', y=0.83, x=0.5, horizontalalignment='center', size=4.5, transform=ax.transAxes)
            
        ax.set_xlabel(xlabel, fontsize=axlabelsize, labelpad=5)
        ax.spines.right.set_visible(False)
        ax.spines.top.set_visible(False)
        ax.text(s=letters[c], y=1.1, x=0, size=titlesize, fontweight='bold', transform=ax.transAxes)#x=-0.12 to put label far left
        ax.tick_params(axis='y', which='major', labelsize=axticklabelsize)
        
fig.tight_layout()
fig.subplots_adjust(wspace=0.2)
fig.savefig('Figure 2.png')

# %% [markdown]
*Note.* **A-C.** Multiple-choice response distributions. Annotations indicate significant differences between modalities in the proportion of times a specific response option was chosen, after Bonferroni corrections: n.s. = p > 0.05, * = 0.001 < p ≤ 0.05, ** = p ≤ 0.001. Responses shown only from those who experienced imagery in both visual and auditory modalities (multimodal imagers). **D-E.** Histograms of responses from multimodal imagers taken using a sliding scale. Responses are grouped into 1s bins (lower-limit inclusive and upper-limit exclusive), except for the final bin, which shows the proportion of all responses of ≥ 10s. Crosses indicate the median response for each modality. **D.** Auditory imagery had a significantly shorter onset latency than visual imagery. **E.** Auditory imagery has a significantly longer duration than visual imagery.

# %% [markdown]
# Movement-dependence: Which body movements were required for mental imagery?

# %% [markdown]
Q4.21, Q5.21, Q4.23, Q5.23, Q4.24, Q5.24

# %% [code]
# Count whether or not movement was required generally
qnum = 'QX.21'
data = filtered[['Q4.' + qnum.split('.')[-1], 'Q5.' + qnum.split('.')[-1]]]
data.columns = ['visual','auditory']
props = countPair(getMultimodals(data))

# Make a df with the proportion of respondents who agree to requiring each specific body part movement
bodylist = ['head','brow','eyelids','eyes','ears','cheeks','nose','mouth','tongue','throat','arms/hands','torso','legs/feet']
bodypropsdf = pd.DataFrame(index=bodylist, columns=['visual','auditory'])
for i in range(1,14):
    neutralqid = '.23#1_' + str(i)
    bodypart = bodylist[i-1]
    bodypartdata = filtered.loc[:,['Q4'+neutralqid,'Q5'+neutralqid]]
    bodypartprops = countPair(getMultimodals(bodypartdata))
    bodypropsdf.loc[bodypart,'visual'] = bodypartprops.loc['Agree','Q4'+neutralqid]
    bodypropsdf.loc[bodypart,'auditory'] = bodypartprops.loc['Agree','Q5'+neutralqid]
top3visual = bodypropsdf['visual'].sort_values(ascending=False).iloc[:3]
top3auditory = bodypropsdf['auditory'].sort_values(ascending=False).iloc[:3]

# Figure out proportion of respondents who said that some necessary movement was not listed
qnum2 = 'QX.24'
data2 = filtered[['Q4.' + qnum2.split('.')[-1], 'Q5.' + qnum2.split('.')[-1]]]
props2 = countPair(data2)

# %% [markdown]
## (this text requires Zotero citations in the main document)

# %% [code]
text=(f"Previous research indicates that mental imagery often involves motor activity related to the content being imagined. For instance, eye movements during visual imagery tend to relate to the spatial content of visual mental images (Fourtassi et al., 2017), while muscle movements related to vocalisation are detectable when imagining sounds (Pruitt et al., 2019). Here we probed whether or not participants believed that they required any such movements to perform imagery in each modality. Specifically, we asked participants to report whether they could see or hear their thought content while remaining completely still, or if they had to move some part of their body in some way for imagery to occur (Figure 3A). We also probed which specific body parts were most frequently reported as necessary for imagery in each modality (Figure 3B). The majority of multimodal imagers reported that body movements were not required for imagery in both modalities. However {round(props.loc['Movement required','visual']/props.loc['Movement required','auditory'],1)}x more multimodal imagers reported requiring movement for visual, but not auditory, imagery ({report(getMultimodals(data))})." + "<br>" +

f"Despite most multimodal imagers reporting that movement was not required for imagery when asked broadly, many more respondents reported that body movements were required when they were asked about the involvement of individual body parts one-by-one. Figure 3B shows the proportion of multimodal imagers who reported that a particular body movement was necessary for imagery. Respondents may have had difficulty introspecting on the involvement of body parts during imagery until they were required to systematically test a list of body parts. They may also have interpreted the phrasing of the explicit question concerning whether imagery occurred “even if you were remaining completely still” to refer to body, but not head, stillness. For visual imagery, the three most frequently reported body movements, from most-reported to least-reported, were the {', '.join(list(top3visual.index)[:2])}, and {list(top3visual.index)[-1]}. Note that, for visual imagery, the majority of multimodal imagers reported that seeing thought content required eye movements ({top3visual['eyes']}%) when explicitly asked about eye movements, yet only a minority reported that movements of any form were required when asked about body movements generally. For auditory imagery, few participants reported that body movements were necessary when asked broadly or specifically, with the three most frequently reported body movements being those of the {', '.join(list(top3auditory.index)[:2])}, and {list(top3auditory.index)[-1]}. We do note that, for both modalities, participants were first instructed to conduct imagery with their eyes closed and then comment on their experience at the time. However, participants were explicitly instructed not to report body movements purely associated with closing their eyes for the task." + "<br>" +

f"In either modality, {props2.loc['Yes (please specify)',:].max()}% or fewer multimodal imagers reported that they needed to move a body part which was not listed in Figure 3B. However, almost all open text responses (Supplementary S1.8) regarding the unlisted body part referred to body parts already listed, or related to changes indirectly related to imagery (e.g. moving to change posture or put something away, or moving in order to keep breathing). One person mentioned that they had to rub their closed eyes to produce visual imagery. Jaw movements and stomach or diaphragm movements were each mentioned once for auditory imagery. ")

md(text)


# %% [markdown]
## **Figure 3**
Body movement during visual and auditory mental imagery, and beliefs about mental imagery

# %% [code]
# Combine movement dependence questions and also terminology/normalness/averageness

# Body parts
# Make a dictionary of neutralid:bodypart, and then one of bodypart:visaud data, then dataframe with columns of modality, indices of bodypart names
bodydata = filtered.copy()
bodylist = ['Head','Brow','Eyelids','Eyes','Ears','Cheeks','Nose','Mouth','Tongue','Throat','Arms/hands','Torso','Legs/feet']
bodyqlist = []
bodyqdic = {}
bodyqdatadic = {}
bodypropsdf = pd.DataFrame(index=bodylist, columns=['visual','auditory'])
subjectcountmin = 999
subjectcountmax = 0
for i in range(1,14):
    neutralqid = '.23#1_' + str(i)
    bodypart = bodylist[i-1]
    bodyqlist.append(neutralqid)
    bodyqdic[neutralqid] = bodypart
    bodyqdata = bodydata.loc[:,['Q4'+neutralqid,'Q5'+neutralqid]]
    bodyqdatadic[bodypart] = bodyqdata
    multimodals = getMultimodals(bodyqdata)
    if len(multimodals) < subjectcountmin: subjectcountmin = len(multimodals)
    if len(multimodals) > subjectcountmax: subjectcountmax = len(multimodals)
    bodyqprops = countPair(multimodals)
    bodypropsdf.loc[bodypart,'visual'] = bodyqprops.loc['Agree','Q4'+neutralqid]
    bodypropsdf.loc[bodypart,'auditory'] = bodyqprops.loc['Agree','Q5'+neutralqid]

# Meta/belief questions
beliefqdic = {'Q4.45':'normality','Q5.45':'normality','Q4.47':'averageness','Q5.47':'averageness','Q4.49':'terminology','Q5.49':'terminology'}
beliefdata = prehall_filtered.copy()
splitresultsdic = {'visual':{},'auditory':{}}
# Magically split the first qnum by the responses of the second qnum
XsplitbyYpairs = [['Q4.45', 'Q4.47'],['Q4.47', 'Q4.45'],['Q4.49', 'Q4.45'],['Q5.45', 'Q5.47'],['Q5.47', 'Q5.45'],['Q5.49', 'Q5.45']]
for pair in XsplitbyYpairs:
    splitdf = beliefdata.groupby(pair).size().unstack()
    splitdf = round(100*splitdf/(splitdf.sum().sum()),1)
    if 'Q4' in pair[0]:
        splitresultsdic['visual'][f'{beliefqdic[pair[0]]}|{beliefqdic[pair[-1]]}'] = splitdf
    elif 'Q5' in pair[0]:
        splitresultsdic['auditory'][f'{beliefqdic[pair[0]]}|{beliefqdic[pair[-1]]}'] = splitdf

# Begin figure creation
fig = plt.figure(dpi=600, figsize=(6,6))
spec = mpg.GridSpec(ncols=6, nrows=2)
axes = []
axes.append(fig.add_subplot(spec[0,:2]))
axes.append(fig.add_subplot(spec[0,2:]))
axes.append(fig.add_subplot(spec[1,:2]))
axes.append(fig.add_subplot(spec[1,2:4]))
axes.append(fig.add_subplot(spec[1,4:]))
colours = ['cornflowerblue','orchid']
cmap3 = mpcm.get_cmap('Greens')
cmap4 = mpcm.get_cmap('Reds')
colours3 = [cmap3(0.5),cmap3(0.7),cmap3(0.9)]
colours4 = [cmap4(0.4),cmap4(0.6),cmap4(0.8), cmap4(0.99)]
letters = 'ABCDEF'
axlabelsize = 6
axticklabelsize = 6
titlesize = 10
markersize=5
allow_swapping_to_unimodal_plots = False
for c in range(5):
        if c == 0:
            qdata = bodydata.loc[:,['Q4.21','Q5.21']]
            group = 'multimodals'
            title = 'Movement- \ndependence'
            subtitle = 'Were body-movements required\n for imagery to occur?'
            order = ['movement required','movement not required']
            xlabel = 'dependence of imagery\n on body movement'
        elif c == 1: 
            group = 'multimodals'
            thisdf = bodypropsdf
            title = 'Body-parts\n required'
            subtitle = 'Which specific body parts needed to be \nmoved for imagery to occur?'
            order = [responselabel.lower() for responselabel in bodylist]
            xlabel = 'part of body which must move for imagery to occur'
        elif c == 2: 
            qdata = beliefdata[['Q4.45','Q5.45']] # Dataset stats are done on, but not necessarily data that is plotted
            group = 'all' 
            split = 'normality|averageness'
            title = 'Typicality of\n imagery'
            subtitle = 'Is it normal to see/hear \nyour thought content?\n(stacked by relative imagery ability)'
            order = ['normal for others, normal for me','normal for others, abnormal for me','abnormal for others, normal for me','abnormal for others, abnormal for me']
            splitlegend = ['less than average','average','more than average']
            legendtitle = 'Relative imagery ability'
            xlabel = 'self-reported normality \nof imagery experience'
        elif c == 3:
            qdata = beliefdata[['Q4.47','Q5.47']]
            group = 'all'
            split = 'averageness|normality'
            title = 'Relative \nimagery ability'
            subtitle = 'Are your thoughts more or less \nvisible/audible than average?\n(stacked by typicality of imagery)'
            order = ['less than average','average','more than average']
            splitlegend = ['normal for others, normal for me','normal for others, abnormal for me','abnormal for others, normal for me','abnormal for others, abnormal for me']
            legendtitle = 'Typicality of imagery'
            xlabel = 'self-reported judgement\n of imagery ability'
        elif c == 4:
            qdata = beliefdata[['Q4.49','Q5.49']]
            group = 'all'#'multimodals' #Changing this will not change actual values here; multimodaling needs to be done at the magic stacking stage earlier
            split = 'terminology|normality'
            title = 'Imagery \nterminology'
            subtitle = 'Which term best describes your experience \nwhen seeing/hearing thought content?\n(stacked by typicality of imagery)'
            order = ['mental imagery','hallucination','other','no sensory experience']
            splitlegend = ['normal for others, normal for me','normal for others, abnormal for me','abnormal for others, normal for me','abnormal for others, abnormal for me']
            legendtitle = 'Typicality of imagery'
            xlabel = 'chosen term'
        
        # Choose group to include
        if group == 'multimodals':
            qdata = getMultimodals(qdata)
            if 'no sensory experience' in order:
                order.remove('no sensory experience')
        elif group == 'unimodals':
            qdata = getUnimodals(qdata)
            order.append('no sensory experience')
        
        # Get stats
        if c ==1:
            stats2x2 = [testSquareTable(qdata, collapse2x2='Agree', verbose=False)[1].pvalue for qdata in bodyqdatadic.values()]
            bonfactor = (len(stats2x2) if len(stats2x2) > 2 else 1)
            annotations = ['**' if p <= (0.05/bonfactor)/50 else '*' if p <= 0.05/bonfactor else 'n.s.' for p in stats2x2]
        else:
            stats2x2 = [testSquareTable(qdata, collapse2x2=responselabel.capitalize(), verbose=False)[1].pvalue for responselabel in order]
            bonfactor = (len(stats2x2) if len(stats2x2) > 2 else 1)
            annotations = ['**' if p <= (0.05/bonfactor)/50 else '*' if p <= 0.05/bonfactor else 'n.s.' for p in stats2x2]
            thisdf = countPair(qdata)
            subjectcount = len(qdata)
        
        # Create plots
        if c == 0:
            thisdf.columns = ['visual','auditory']
            thisdf.index = [text.lower() for text in thisdf.index]
            thisdf = thisdf.loc[order,:]
            thisdf.index = [textwrap.fill(text,10,break_long_words=False) for text in thisdf.index] # Wraps text
            ax = thisdf.plot(kind='bar', color=colours, rot=0, ax=axes[c])
            ax.text(s=f'n = {subjectcount}', y=0.9, x=0.5, horizontalalignment='center', size=5, transform=ax.transAxes)
        elif c == 1:
            thisdf.index = [text.lower() for text in thisdf.index]
            thisdf = thisdf.loc[order,:]
            thisdf.index = [text.replace('/','/\n') if '/' in text else text for text in thisdf.index] # Wraps text
            ax = thisdf.plot(kind='bar', color=colours, rot=0, ax=axes[c], width=0.7)
            ax.yaxis.set_visible(False)
            ax.spines.left.set_visible(False)
            ax.text(s=f'n = {subjectcountmin}\u2014{subjectcountmax}', y=0.9, x=0.5, horizontalalignment='center', size=5, transform=ax.transAxes)
        elif c in [2,3,4]:
            ax = axes[c]
            for modality in ['visual','auditory']:
                thisdf = splitresultsdic[modality][split]
                thisdf.index = [text.lower() for text in thisdf.index]
                thisdf = thisdf.loc[order]
                thisdf.index = [textwrap.fill(text,12,break_long_words=False) for text in thisdf.index] # Wraps text
                thisdf.columns = [text.lower() for text in thisdf.columns]
                thisdf = thisdf.loc[:,splitlegend]
                if c==2: colours=colours3
                else: colours=colours4
                if modality == 'visual': 
                    thisdf.plot(kind='bar', stacked=True, color=colours, rot=0, ax=ax, width=.35, position=1) # These positions seem odd, but work
                elif modality == 'auditory':
                    thisdf.plot(kind='bar', stacked=True, color=colours, rot=0, ax=ax, width=.35, position=0)
        
        # Set ylims so stats annotations can use them
        ax.set_ylim((0,139))
        ax.set_yticks([0, 20, 40, 60, 80, 100])
        
        # Add stats annotations
        for i, annotation in enumerate(annotations):
            x = ax.get_xticks()[i]
            width = ax.patches[0].get_width()   
            xleft, xright, height = x-(width/2), x+(width/2), (ax.get_ylim()[1]/10)
            if c == 1: y = thisdf.iloc[i,:].max()
            else: y = countPair(qdata).loc[order[i].capitalize(),:].max()
            ax.plot([xleft, xright], [y+height, y+height], lw=width, c='black')
            if '*' in annotation:
                ax.text(x, y+height-(axticklabelsize/2), annotation, ha='center', va='bottom', color='black', size=1.5*axticklabelsize)
            else:
                ax.text(x, y+height+(axticklabelsize/2), annotation, ha='center', va='bottom', color='black', size=axticklabelsize)        

        # Customise ax plot
        subjectcount = len(qdata)
        if c == 0:
            ax.bar_label(ax.containers[0], label_type='edge', fontsize=axticklabelsize, fmt='%.0f')
            ax.bar_label(ax.containers[1], label_type='edge', fontsize=axticklabelsize, fmt='%.0f')    
        if c == 1:
            ax.bar_label(ax.containers[0], label_type='edge', fontsize=5, fmt='%.0f')
            ax.bar_label(ax.containers[1], label_type='edge', fontsize=5, fmt='%.0f')    
            ax.set_xticklabels(thisdf.index, fontsize=4.5)
        if c == 2:
            ax.set_xticklabels(thisdf.index, fontsize=4)
            ax.legend(splitlegend[::-1], fontsize=4, frameon=False, labelcolor=colours[::-1], handlelength=0, handleheight=0, handletextpad=0, bbox_to_anchor=(0,-0.4,1,1), bbox_transform = ax.transAxes)
            ax.text(s=f'{legendtitle.lower()}:', y=0.6, x=0.6, horizontalalignment='left', size=4, transform=ax.transAxes)
        if c == 3:
            ax.set_xlim((-0.5,2.5))
            ax.get_legend().remove()
        if c == 4:
            ax.set_xticklabels(thisdf.index, fontsize=5)
            ax.legend(splitlegend[::-1], fontsize=4, frameon=False, labelcolor=colours[::-1], handlelength=0, handleheight=0, handletextpad=0, bbox_to_anchor=(0,-0.4,1,1), bbox_transform = ax.transAxes)
            ax.text(s=f'{legendtitle.lower()}:', y=0.6, x=0.3, horizontalalignment='left', size=4, transform=ax.transAxes)
        if c in [0,1]:
            ax.legend(fontsize=axticklabelsize, frameon=False, labelcolor=colours, handlelength=0, handletextpad=0, loc='upper left')
            ax.text(s=subtitle, y=1.05, x=0.5, horizontalalignment='center', size=5, transform=ax.transAxes, verticalalignment='top')
        if c in [0,3]:
            ax.set_xticklabels(thisdf.index, fontsize=axticklabelsize)
        if c in [3,4]:
            ax.yaxis.set_visible(False)
            ax.spines.left.set_visible(False)
        if c in [2,4]:
            ax.set_xlim((-0.5,3.5))   
        if c in [2,3,4]:
            ax.tick_params(axis='x', which='major', pad=8)
            ax.text(s=f'n = {subjectcount}', y=0.85, x=0.5, horizontalalignment='center', size=5, transform=ax.transAxes)
            ax.text(s=subtitle, y=1.05, x=0.5, horizontalalignment='center', size=5, transform=ax.transAxes, verticalalignment='top')
            
            # Literally just to get the position at the top of each bar and plot bar totals/labels
            barxys = []
            xys={}
            for container in ax.containers:
                for patch in container.patches:
                    x=patch.xy[0] + patch._width/2
                    y=patch.xy[1] + patch._height
                    if x in xys.keys():
                        xys[x].append(y)
                    else:
                        xys[x] = [y]
            for x in xys.keys():
                barxys.append((x, max(xys[x])))
            modtext = ['vis','aud']
            barxys.sort()
            for i, xy in enumerate(barxys):
                ax.text(s=int(xy[1]),x=xy[0], y=xy[1]+0.5, fontsize=axticklabelsize, horizontalalignment='center', verticalalignment='bottom')#, color=modcolours[i%2])
                ax.text(s=modtext[i%2],x=xy[0], y=-7, fontsize=4.5, horizontalalignment='center', verticalalignment='bottom')#, color=modcolours[i%2]

        # General customisation
        ax.spines.right.set_visible(False)
        ax.spines.top.set_visible(False)
        ax.spines.left.set_bounds(0, 100)
        ax.set_xlabel(xlabel, fontsize=axlabelsize, labelpad=5)
        ax.set_title(title, y=1.1, x=0.5, size=titlesize, pad=0, fontweight='bold')
        ax.text(s=letters[c], y=1.2, x=0, size=titlesize, fontweight='bold', transform=ax.transAxes)#x=-0.12 to put label far left
        ax.tick_params(axis='y', which='major', labelsize=axticklabelsize)
        ax.set_ylabel('% respondents', fontsize=axlabelsize, labelpad=5)
            
fig.tight_layout()
fig.subplots_adjust(wspace=0.15)
fig.savefig('Figure 3.png')

# %% [markdown]
*Note.* **A-E.** Multiple-choice response distributions. Annotations indicate significant differences between modalities in the proportion of times a specific response option was chosen, after Bonferroni corrections: n.s. = p > 0.05, * = 0.001 < p ≤ 0.05, ** = p ≤ 0.001. **A.** Responses shown only from those who experienced imagery in both visual and auditory modalities (multimodal imagers). **B**. Responses shown only for multimodal imagers. Participants answered a separate question for each body part. **C-E.** Beliefs about mental imagery, with responses in each column split by response choice in neighbouring questions. Statistics apply only to comparisons of proportions for the main question; the breakdown of each column by secondary response was not assessed statistically. Responses presented in C-F include those from respondents who reported mental imagery as a hallucination in E.

# %% [markdown]
# Coarse position and location control: Where did mental images seem to be located, and could the apparent location of mental images be controlled?

# %% [markdown]
Q4.28, Q5.28, Q4.37, Q5.37; Q4.30, Q4.39, Q5.30, Q5.39

# %% [code]
qs = ['Q4.30','Q4.39','Q5.30','Q5.39']
qlabels = ['visual, eyes open','visual, eyes closed','auditory, eyes open','auditory, eyes closed']

# %% [code]
text=(f"We investigated the spatial properties of imagery, namely whether the spatial location of seen or heard thought content could be controlled, and where such thought content most frequently seemed to originate from or be spatially positioned at. We investigated each property during both eyes-open and eyes-closed imagery. For respondents that experienced imagery in both modalities and eye states, a significantly larger proportion of respondents could control the apparent position of seen thought content rather than heard thought content (Figure 4A), both when eyes were open ({report(getMultimodals(filtered[['Q4.28','Q5.28']]))}) and closed ({report(getMultimodals(filtered[['Q4.37','Q5.37']]))}). Eye-closure did not significantly affect the proportion of these respondents reporting that they could control their visual imagery ({report(getMultimodals(filtered[['Q4.28','Q4.37']]))}) or auditory imagery ({report(getMultimodals(filtered[['Q5.28','Q5.37']]))})." + "<br>" +
f"As for the reported position of imagined sounds and images, when asked in a multiple choice question, most individuals reported that imagined sounds and images were located within their head, except for during eyes-open visual imagery (Figure 4B). Eyes-open visual imagery was significantly less frequently reported as occurring in the head compared to eyes-closed visual imagery ({report(getMultimodals(filtered[['Q4.30','Q4.39']]), Collapse2x2='Head')}), and relative to eyes-open auditory imagery ({report(getMultimodals(filtered[['Q4.30','Q5.30']]), Collapse2x2='Head')}). Instead, images visualised with eyes open were significantly more likely to be reported as being positioned in the environment relative to eyes-closed imagined images ({report(getMultimodals(filtered[['Q4.30','Q4.39']]), Collapse2x2='Environment')}), or eyes-open imagined sounds ({report(getMultimodals(filtered[['Q4.30','Q5.30']]), Collapse2x2='Environment')})."
      )
md(text)

# %% [markdown]
# Precise spatial position of mental imagery

# %% [markdown]
Q4.32, Q5.32, Q4.34, Q5.34, Q4.41, Q5.41, Q4.43, Q5.43

# %% [code]
# Create masks for each region in case we want to spatially bin the precise data into categories of head, body, or environment
plotbinnedcoordinates = False
# Body profile: set the region masks
profilemask = cv2.imread('Supplementary S5.png')
profileheadmask = profilemask[:,:,2] > 0
profilebodymask = profilemask[:,:,0] > 0
profileenviromask = profilemask[:,:,1] > 0
# Top-down head: set the region masks
topdownmask = cv2.imread('Supplementary S7.png')
topdownheadmask = topdownmask[:,:,2] > 0
topdownenviromask = topdownmask[:,:,1] > 0

# Question numbers
qdic = {'open sideview visual':'Q4.32','open sideview auditory':'Q5.32','open topview visual':'Q4.34','open topview auditory':'Q5.34',
         'closed sideview visual':'Q4.41','closed sideview auditory':'Q5.41','closed topview visual':'Q4.43','closed topview auditory':'Q5.43'}
binnedregiondic = {}
coordsdic = {}

# Get coordinates and bin them into categorical regions as well
for q in qdic.keys():
    qdf = pd.DataFrame(columns=[q])
    qy = qdic[q]+'_1_y'
    qx = qdic[q]+'_1_x'
    coords = filtered[[qy,qx]]
    coorddf = pd.DataFrame(columns=['y','x'])
    for index, row in coords.iterrows():
        categoricalresponse = None
        y=row[0]
        x=row[1]
        # Remove nans, convert from string number
        if type(y) == type(""):
            y=int(y)
            x=int(x)
            # Store coordinates so we can visualise heatmaps later
            coorddf.at[index,'y'] = y
            coorddf.at[index,'x'] = x
            # Check the right regions depending on the image used
            if 'sideview' in q:
                if profileheadmask[y,x]: categoricalresponse = 'Head'
                if profilebodymask[y,x]: categoricalresponse = 'Body'
                if profileenviromask[y,x]: categoricalresponse = 'Environment'
            elif 'topview' in q:
                if topdownheadmask[y,x]: categoricalresponse = 'Head'
                if topdownenviromask[y,x]: categoricalresponse = 'Environment'
        else:
            categoricalresponse = 'No sensory experience'
        qdf.at[index,q] = categoricalresponse
    categories = ['No sensory experience','Head','Environment']
    if 'sideview' in q:
        catinfo = pd.api.types.CategoricalDtype(categories=categories+['Body'], ordered=False)
    if 'topview' in q:
        catinfo = pd.api.types.CategoricalDtype(categories=categories, ordered=False)
    binnedregiondic[q] = qdf.astype(catinfo)
    coordsdic[q] = coorddf

# Plot proportion of binned coordinates in categorical regions, if needed
if plotbinnedcoordinates:
    fig, axes = plt.subplots(ncols=2, nrows=2, figsize=(15,6))
    axes=axes.ravel()
    figlabs = ['A','B','C','D']
    for c, ax in enumerate(axes):
        if c in [0,2]: view = 'sideview'
        elif c in [1,3]: view = 'topview'
        if c in [0,1]: modality = 'visual'
        elif c in [2,3]: modality = 'auditory'
        tempdf = 100*pd.concat((binnedregiondic[f'open {view} {modality}'].value_counts(normalize=True), 
                                binnedregiondic[f'closed {view} {modality}'].value_counts(normalize=True)), axis=1)

        tempdf.columns = [f'eyes open, {view}, {modality}',f'eyes closed, {view}, {modality}']
        tempdf.plot(kind='bar', color=['Red','Orange'], ax=ax)
        ax.bar_label(ax.containers[0], label_type='edge', fontsize=10, fmt='%.0f')
        ax.bar_label(ax.containers[1], label_type='edge', fontsize=10, fmt='%.0f')
        if view == 'sideview':
            ax.set_xticklabels(['environment','head','no sensory \nexperience','body'], fontsize=10, rotation=0)
        elif view == 'topview':
            ax.set_xticklabels(['environment','head','no sensory \nexperience'], fontsize=10, rotation=0)
        ax.set_ylabel('% respondents')
        ax.set_title(f'data from Fig {figlabs[c]}')
        ax.set_ylim((0,85))
    fig.tight_layout()

# Convert from pixels to vaguely interpretable metrics
av_height_in_cm = (round((163.8+177.8)/2,0)) # Height of Australians from: https://www.abs.gov.au/ausstats/abs@.nsf/lookup/4338.0main+features212011-13
av_head_circumference = 56.0 #56cm number estimated from EEG experience
av_head_diameter_in_cm = av_head_circumference/np.pi 
approx_image_height_in_pixels = 1125*((7.31-2.12)/7.81)
approx_image_head_diameter_in_pixels = (1280*((7.25-6.15)/13.34) + 720*((4.43-3.2)/7.5))/2
sideviewpix2cm = av_height_in_cm/approx_image_height_in_pixels
topviewpix2cm = av_head_diameter_in_cm/approx_image_head_diameter_in_pixels

# Compare the coordinates as continuous data, and do stats
comparisons = [('open sideview visual','closed sideview visual'),('open topview visual','closed topview visual'),('open sideview auditory','closed sideview auditory'),('open topview auditory','closed topview auditory')]
meandiffs = {'y':[],'x':[]}
devdiffs = {'y':[],'x':[]}
wilcoxons = {'y':[],'x':[]}
levenes = {'y':[],'x':[]}
for comparison in comparisons:
    for axis in ['y','x']:
            a = coordsdic[comparison[0]][axis]
            b = coordsdic[comparison[1]][axis]
            pairdf = pd.concat((a,b), axis=1)
            pairdf = pairdf.dropna(how='any')
            pairdf.columns = ['a','b']
            if 'topview' in comparison[0]: 
                multiplier = topviewpix2cm
            if 'sideview' in comparison[0]: 
                multiplier = sideviewpix2cm
            wilcoxons[axis].append(reportWilcoxon(pairdf['a'],pairdf['b']))
            levenes[axis].append(reportLevene(pairdf['a'],pairdf['b']))
            meandiffs[axis].append(((round(multiplier*(pairdf['a'].mean() - pairdf['b'].mean()),1))))
            devdiffs[axis].append(((round(multiplier*(pairdf['a'].std() - pairdf['b'].std()),0))))

# %% [code]
text=(f"If participants reported a visual or auditory experience of their thoughts for a particular eye state, we also asked them to indicate the apparent location of the thought content they were seeing or hearing by placing a single marker on an image of a human in a room, as viewed from the side (Figure 4C & E) or above (Figure 4D & F). This allowed us to more precisely visualise where in space imagined images and sounds were perceived as originating from or being located at. In general, imagined content was perceived either in the head, directly in front of or beside the head, or projected forward from the head. When eyes were open, the content of visual mental images appeared to shift forward compared to when eyes were closed, projecting away from the head and into the environment. We were able to estimate this distance in centimetres by assuming an average height of {av_height_in_cm}cm and an average head circumference of {av_head_circumference}cm for the figures portrayed in our images. Two-tailed Wilcoxon signed-rank tests confirmed a statistically significant forward projection of visual imagery when eyes were open rather than closed, with a mean projection of approximately {abs(meandiffs['x'][0])}cm along the horizontal axis in Figure 4C ({wilcoxons['x'][0]}) and {abs(meandiffs['y'][1])}cm along the vertical axis in Figure 4D ({wilcoxons['y'][1]}). Levene tests also showed that the reported imagery position was more variable along these axes when eyes were open relative to closed, with standard deviation increases of {devdiffs['x'][0]}cm ({levenes['x'][0]}) and {devdiffs['y'][1]}cm ({levenes['y'][1]}) respectively. Up-down position variance also increased with eye opening along the vertical axis of Figure 4C, with a standard deviation increase of {devdiffs['y'][0]}cm ({levenes['y'][0]}). All other comparisons of variance or position between an eyes-open and eyes-closed state within a modality were non-significant after Bonferroni correction (p > .013)."
     )
md(text)

# %% [markdown]
## **Figure 4**
Spatial properties of visual and auditory mental imagery

# %% [code]
# !This cell requires coordsdic from previous cell

# Make a colourmap that includes transparency
qdic = {'open sideview visual':'Q4.32','open sideview auditory':'Q5.32','open topview visual':'Q4.34','open topview auditory':'Q5.34',
         'closed sideview visual':'Q4.41','closed sideview auditory':'Q5.41','closed topview visual':'Q4.43','closed topview auditory':'Q5.43'}

# Load images
sideviewimage = mpimg.imread('Supplementary S4.png')
topviewimage = mpimg.imread('Supplementary S6.png')

# Get data
data = filtered.copy()

# Establish plotting parameters
fig = plt.figure(dpi=600, figsize=(6,6), layout='constrained')
spec = mpg.GridSpec(figure=fig, ncols=2, nrows=3)
axes = []
axes.append(fig.add_subplot(spec[0,0:1]))
axes.append(fig.add_subplot(spec[0,1:2]))
axes.append(fig.add_subplot(spec[1,0:1]))
axes.append(fig.add_subplot(spec[1,1:2]))
axes.append(fig.add_subplot(spec[2,0:1]))
axes.append(fig.add_subplot(spec[2,1:2]))

# Plotting custom attributes
colours = ['cornflowerblue','yellowgreen','orchid','sandybrown']
axlabelsize = 6
axticklabelsize = 6
titlesize = 10
colourlist=colours
letters = 'ABCDEF'

for c in range(6):
   
    # For regular bar plots
    if c in [0,1]:
        if c == 0: 
            qdata = getMultimodals(data[['Q4.28','Q4.37','Q5.28','Q5.37']])
            qdata.columns = ['visual, eyes open','visual, eyes closed','auditory, eyes open','auditory, eyes closed']
            props = [qdata[column].value_counts(normalize=True) for column in list(qdata.columns)]
            thisdf = round(100*pd.concat(props, axis=1),1)
            title = 'Location control'
            subtitle = 'Could the apparent location of \nmental images be controlled?'
            order = ['location control','no location control']#,'no sensory experience']
            xlabel = 'ability to control imagery location'
        if c == 1:
            qdata = getMultimodals(data[['Q4.30','Q4.39','Q5.30','Q5.39']])
            qdata.columns = ['visual, eyes open','visual, eyes closed','auditory, eyes open','auditory, eyes closed']
            props = [qdata[column].value_counts(normalize=True) for column in list(qdata.columns)]
            thisdf = round(100*pd.concat(props, axis=1),1)
            title = 'Coarse position'
            subtitle = 'Where did imagery\n seem to be located?'
            order = ['head','body','environment','inconsistent','somewhere else']#,'no sensory experience']
            xlabel = 'apparent spatial location of imagery'
        
        # Do stats
        # Eyes-open vs eyes-closed (within a modality), and also vis vs aud (within a particular eye state)
        statsdic = {'compare eyes':{'visual':None,'auditory':None}, 'compare modalities':{'eyes open':None,'eyes closed':None}}
        for comparison in statsdic.keys():
            for aspect in statsdic[comparison].keys():
                if comparison == 'compare eyes': pair = qdata[[f'{aspect}, eyes open',f'{aspect}, eyes closed']]
                if comparison == 'compare modalities': pair = qdata[[f'visual, {aspect}',f'auditory, {aspect}']]
                ys = [countPair(pair).loc[label.capitalize(),:].max() for label in order]
                stats2x2 = [testSquareTable(pair, collapse2x2=label.capitalize(), verbose=False)[1].pvalue for label in order]
                bonfactor = (len(stats2x2) if len(stats2x2) > 2 else 1)
                annotations = ['**' if p <= (0.05/bonfactor)/50 else '*' if p <= 0.05/bonfactor else 'n.s.' for p in stats2x2]
                statsdic[comparison][aspect] = (annotations,ys)
                
        # Sort labels, then make plots
        thisdf.index = [text.lower() for text in thisdf.index]
        thisdf = thisdf.loc[order,:]
        thisdf.index = [textwrap.fill(text,11,break_long_words=False) for text in thisdf.index] # Wraps text
        ax = thisdf.plot(kind='bar', color=colourlist, rot=0, ax=axes[c], ylim=(0,80), width=0.8)

        # Plot stats
        # Keep track of previously plotted heights for each xtick
        yhistorydic = {}
        for xtick in ax.get_xticks():
            yhistorydic[xtick] = []
        for comparison in statsdic.keys():
            for aspect in statsdic[comparison].keys():
                annotations = statsdic[comparison][aspect][0]
                ys = statsdic[comparison][aspect][1]
                
                for i, annotation in enumerate(annotations):
                    x = ax.get_xticks()[i]
                    y, height = ys[i], (ax.get_ylim()[1]/10)
                    width = ax.patches[0].get_width()  
                    
                    for yhistorical in yhistorydic[x]:
                        while abs(y-yhistorical) < 8:
                            y += 1

                    # Set annotation positions
                    if aspect == 'visual':
                        xleft, xright = x-(1.5*width), x-(width/2)
                    elif aspect == 'auditory':
                        xleft, xright = x+(width/2), x+(1.5*width)
                    elif aspect == 'eyes open':
                        xleft, xright = x-(1.5*width), x+(width/2)
                    elif aspect == 'eyes closed':
                        xleft, xright = x-(width/2), x+(1.5*width)
                    
                    # Only plot if significant here to avoid crowding with non-significant annotations
                    if '*' in annotation:
                        yhistorydic[x].append(y)
                        ax.text((xleft+xright)/2, y+height-(axticklabelsize/2), annotation, ha='center', va='bottom', color='black', size=1.5*axticklabelsize)
                        ax.plot([xleft, xright], [y+height, y+height], lw=width, c='black')

        # Customise plots
        if c == 0:
            ax.get_legend().set_visible(False)
            ax.set_xticklabels(thisdf.index, fontsize=axticklabelsize)
        if c == 1:
            ax.yaxis.set_visible(False)
            ax.spines.left.set_visible(False)
            ax.legend(fontsize=axticklabelsize, frameon=False, labelcolor=colours, handlelength=0, handletextpad=0, loc='center right')
            ax.set_xticklabels(thisdf.index, fontsize=5.5)
        
        subjectcount = len(qdata) 
        ax.set_ylim((0,119))
        ax.set_title(title, y=1.1, x=0.5, size=titlesize, pad=0, fontweight='bold')
        ax.set_xlabel(xlabel, fontsize=axlabelsize, labelpad=5)
        ax.spines.right.set_visible(False)
        ax.spines.top.set_visible(False)
        ax.spines.left.set_bounds(0, 100)
        ax.text(s=letters[c], y=1.1, x=0.1, size=titlesize, fontweight='bold', transform=ax.transAxes)#x=-0.12 to put label far left
        ax.text(s=subtitle, y=.97, x=0.5, horizontalalignment='center', size=axlabelsize, transform=ax.transAxes)
        ax.text(s=f'n = {subjectcount}', y=0.9, x=0.5, horizontalalignment='center', size=axlabelsize, transform=ax.transAxes)
        ax.tick_params(axis='y', which='major', labelsize=axticklabelsize)
        ax.set_ylabel('% respondents', fontsize=axlabelsize, labelpad=5)
        ax.bar_label(ax.containers[0], label_type='edge', fontsize=axticklabelsize, fmt='%.0f')
        ax.bar_label(ax.containers[1], label_type='edge', fontsize=axticklabelsize, fmt='%.0f')
        ax.bar_label(ax.containers[2], label_type='edge', fontsize=axticklabelsize, fmt='%.0f')
        ax.bar_label(ax.containers[3], label_type='edge', fontsize=axticklabelsize, fmt='%.0f')
        
    # For heatmaps
    elif c in [2,3,4,5]:
        if c in [2,3]: 
            modality = 'visual'
            colidx1 = 0
        elif c in [4,5]: 
            modality = 'auditory'
            colidx1 = 2
        if c in [2,4]:
            image = sideviewimage
            view = 'sideview'
        elif c in [3,5]:
            image = topviewimage
            view = 'topview'
        axes[c].imshow(image,zorder=0)
        axes[c].axis('off')                          
        for eyestate in ['open','closed']:
            if eyestate == 'open':
                colidx2 = 0
                zorder = 2
            else:
                colidx2 = 1
                zorder = 1
            colidx = colidx1 + colidx2
            name = f'{eyestate} {view} {modality}'
            leg = f'{modality}, eyes {eyestate}'
            df = coordsdic[name]
            yaverage, xaverage = df['y'].mean(), df['x'].mean()
            axes[c].scatter(y=df['y'], x=df['x'], color=colourlist[colidx], alpha=0.33, s=20, marker='.', facecolors=colourlist[colidx], edgecolors=colourlist[colidx], linewidth=.1, zorder=zorder, label=leg)
            axes[c].scatter(y=yaverage, x=xaverage, alpha=1, s=20, marker='X', facecolors='white', edgecolors=colourlist[colidx], linewidth=1, zorder=zorder+1)
            axes[c].set_ylim((image.shape[0],0))
            axes[c].set_xlim((0,image.shape[1]))
            axes[c].legend(fontsize=axticklabelsize, loc='lower left', frameon=False, bbox_to_anchor=(-0.01, 0.05, 1, 1), labelcolor=[colourlist[colidx1], colourlist[colidx1+1]], markerscale=0)
            axes[c].text(s=letters[c], y=0.83, x=0.1, size=titlesize, fontweight='bold', transform=axes[c].transAxes)#x=-0.12 to put label far left
            if eyestate == 'closed': axes[c].text(s=f'n = {len(df)}', y=0.11, x=0.775, size=axticklabelsize, transform=axes[c].transAxes, color=colourlist[colidx])
            if eyestate == 'open': axes[c].text(s=f'n = {len(df)}', y=0.11+0.075, x=0.775, size=axticklabelsize, transform=axes[c].transAxes, color=colourlist[colidx])

fig.savefig('Figure 4.png')

# %% [markdown]
*Note.* **A-B.** Multiple-choice response distributions. Annotations indicate significant differences between modalities in the proportion of times a specific response option was chosen, after Bonferroni corrections: * = 0.001 < p ≤ 0.05, ** = p ≤ 0.001. Responses shown only from those who experienced imagery in both visual and auditory modalities (multimodal imagers). For clarity, non-significant results are not shown, however main effects of modality and eye-state were tested in all combinations. **C-F.** Distribution maps of precise locations of imagery. Each dot indicates a single respondent's chosen location for where imagery most commonly seems to be located or originate from. Crosses indicate mean positions. Participants were not given the option to place a marker if they had indicated that they did not have a sensory experience of their thoughts when previously asked about the coarse location of their imagery categorically. **C-D.** Visual imagery was projected forward, away from the head, when eyes were open, significantly more so than when eyes were closed. **E-F.** Eye state did not significantly affect the location of auditory imagery.

# %% [markdown]
# Typicality, ability, and terminology: Is mental imagery normal, how do individuals compare to others, and which term best describes their experience?

# %% [markdown]
Q4.45, Q5.45; Q4.47, Q5.47; Q4.49, Q5.49

# %% [code]
# normality Q4.45, Q5.45
# averageness Q4.47, Q5.47
# terminology Q4.49, Q5.49
text=(f"We asked participants to report on whether they considered seeing or hearing their thoughts in some way to be a normal experience for themselves and for others (typicality; Figure 3C), whether they considered their thoughts to be more or less sensory than average (ability; Figure 3D), and which term they would use to best describe the experience they most commonly have when seeing or hearing their thought content (terminology; Figure 3E). Note that participants who described their experience as a hallucination were included in the analyses of these three questions despite being excluded from analyses in previous questions. Although most respondents thought that seeing or hearing their thought content was a normal experience for others and themselves, there was a slight bias towards reporting that seeing thought content was normal, even if it was not a normal event for the participant, while hearing thought content was slightly more likely to be considered an abnormal process for all. Most participants felt that their imagery abilities were average, with slightly more participants suspecting that their visual imagery, rather than auditory imagery, was less sensory than average. Note that many of these individuals still recognised visual imagery as a normal process for others even if they had a low self-reported visibility of their own thought content. Significant differences are shown in Figure 3." + "<br>" +

f"""Throughout this study, we deliberately avoided asking participants to report on "mental imagery" for reasons previously detailed (see Methods). Instead, we asked participants to report on whether they in some way saw or heard their thought content, or had a visual or auditory experience of their thought content. We confirm that most participants understood this phenomenon to be synonymous with mental imagery (Figure 3E), although a substantial proportion of respondents described hearing thought content as a hallucination, or using another term, despite using "mental imagery" to describe seeing thought content. This is not unexpected given that "mental imagery" is traditionally associated with the visual modality. Alternative terms provided by respondents for the common experience of hearing one's thoughts are listed in Supplementary S1.8, with most still describing the phenomenon of auditory mental imagery despite not using the term explicitly, or using an auditory-specific term instead e.g. an inner or internal voice or monologue, hearing one's thoughts, or hearing a voice in their head."""
     )
md(text)

# %% [markdown]
# Explicit comparisons of mental imagery in different modalities

# %% [markdown]
Q6.9, Q8.2-8.22 (even numbers)

# %% [code]
text=("""Until now, we have compared imagery modalities by asking participants to make judgements about their thought content in each modality separately and then comparing their judgements after the fact. However, we also asked participants to make explicit judgements comparing imagery properties between modalities. In Figure 5A, we show the distribution of responses from when participants were explicitly asked to compare their experiences and judge which of visual or auditory thought content most met a given criteria. Participants could also report that sensory thoughts in both modalities were experienced in the same way with respect to the criterion being inquired about, or whether they had no sensory experience at all (e.g. "Which is the easiest to do in some way?" "Seeing my thought content in some way", "Hearing my thought content in some way", "There is no difference", or "I do not see or hear my thought content in any way"). We also asked participants to make judgements about not just visual and auditory thought content, but sensory thought content in the five traditional senses (sight, sound, smell, taste, and touch). The distribution of these judgements is shown in Figure 5B. Note that, for these latter judgements, participants could choose multiple sensory modalities if they deemed that the criteria could apply to multiple modalities equally, or they could report that they had no experience in any modality.""" + "<br>" +

"Overall, auditory thought content tended to be most frequently ranked as the best match on each criterion, except for location control. Most participants judged that it was easier to control the apparent location of visual thought content rather than auditory thought content. Although visual imagery was generally less dominant than auditory imagery with respect to most criteria, both of these modalities still tended to eclipse the other sensory modes of thinking for each criterion probed."
     )
md(text)

# %% [markdown]
# Sensory and non-sensory experiences in thoughts, imagination, and dreams

# %% [markdown]
Q7.1, Q7.2, Q7.3

# %% [code]
text=("""Given that mental images can be defined as sensory thoughts, it should follow that seeing or hearing one's thought content is, by definition, mental imagery. However, despite sharing a common root word, "mental imagery" need not necessarily be considered equivalent to, or a part of "imagining", nor need "thinking" in a sensory way be necessarily considered equivalent to "imagining". To explore public intuition concerning these phenomena, we asked participants to provide a binary response as to whether they considered imagining the appearance or sound of a letter to be the same process as thinking about the appearance or sound of the same letter. We also probed which forms of sensory and non-sensory experience participants could recall occurring at least sometimes and in some way during their imagination and thoughts (Figure 5C). To compare to another form of internally-generated phenomenon, we also asked participants to specify their experiences during dreaming.""" + "<br>"

f"Across {len(prehall_filtered)} respondents, {100*prehall_filtered.loc[:,'Q9.12'].value_counts(normalize=True)['Yes']:.1f}% reported imagining and thinking to be the same process regarding visual appearance, and {100*prehall_filtered.loc[:,'Q9.13'].value_counts(normalize=True)['Yes']:.1f}% reported imagining and thinking to be the same regarding sound ({report(prehall_filtered[['Q9.12','Q9.13']])}; Supplementary S1.1, Q9.12 & Q9.13). We also provided an optional free-text field for participants to report on any general differences they perceived between imagining and thinking (i.e. not specifically in reference to thinking or imagining sensory content; Supplementary S1.1, Q9.14). Although  many reported no difference even for this general comparison, recurring comments (Supplementary S1.8) regarded that imagination tended to be more associated with sensory content than thinking, and generally more so with visual content. Many also differentiated imagination and thinking as a difference in content, but not process: imagination was frequently regarded as involving more speculative or creative content which was less grounded in immediate reality compared to thinking." + "<br>" +

"""The incidence of particular sensory or non-sensory experiences during thoughts, imagination, and dreams is summarised in Figure 5C. Detailed statistical comparisons within and between phenomena are listed in Supplementary S1.2 & S1.3. Care must be taken to avoid confusing the proportion of individuals reporting a particular experience with the intensity or degree to which individuals believe that experience to occur. Seeing and hearing tend to be the dominant sensory modalities in each experience. Although most individuals consider the three phenomena to involve largely the same experiences, we also observed that there is a small proportion of respondents who experience more sensory, motor, and emotional content moving from thoughts to imagination to dreams, except concerning the auditory experiences of hearing and speaking, which appear slightly more frequently associated with thinking rather than imagining. However, many of these comparisons were not significant (Supplementary S1.3). Interestingly, touch and movement are significantly more frequently reported in dreams relative to other phenomena (Supplementary S1.3). However, our probe prompt for movement, "I feel like I’m moving as I am in my thoughts/imagination/dreams" may not clearly disambiguate between local movements of body parts and global movements of body position."""
     )
md(text)

# %% [markdown]
## **Figure 5**
Explicit comparisons of mental imagery modalities, thoughts, imagination, and dreams

# %% [code]
data = filtered.copy()
compare2qdic = {'Q6.9#1_1': 'easiest', 'Q6.9#1_2': 'easiest to initiate', 'Q6.9#1_3': 'quickest to initiate', 
                'Q6.9#1_4': 'most sustainable', 'Q6.9#1_5': 'most consistent', 'Q6.9#1_6': 'least movement needed', 
                'Q6.9#1_7': 'most location control','Q6.9#1_8': 'most automatic', 'Q6.9#1_9': 'clearest', 
                'Q6.9#1_10': 'most vivid','Q6.9#1_11': 'realest'}
compare5qdic = {'Q8.2': 'easiest', 'Q8.4': 'easiest to initiate', 'Q8.6': 'quickest to initiate', 
                'Q8.8': 'most sustainable','Q8.10': 'most consistent', 'Q8.12': 'least movement needed',
                'Q8.14': 'most location control', 'Q8.16': 'most automatic', 'Q8.18': 'clearest',
                'Q8.20': 'most vivid', 'Q8.22': 'realest'}
compare3qdic = {'Q7.1#1':'thoughts', 'Q7.2#1':'imagination', 'Q7.3#1':'dreams'}

# Make figure
fig = plt.figure(dpi=600, figsize=(6,6))
spec = mpg.GridSpec(ncols=3, nrows=3)
axes = []
axes.append(fig.add_subplot(spec[0,:]))
axes.append(fig.add_subplot(spec[1,:]))
axes.append(fig.add_subplot(spec[2,:]))
colours2 = ['cornflowerblue','orchid','silver','grey']
colours5 = ['cornflowerblue','orchid','indianred','sandybrown','olivedrab','grey']
cmap = mpc.LinearSegmentedColormap.from_list("", ["white","mediumslateblue"])
colours3 = [cmap(0.4),cmap(0.7),cmap(0.99)]
letters = 'ABCDEF'
axlabelsize = 6
axticklabelsize = 6
titlesize = 10
markersize=5
allow_swapping_to_unimodal_plots = False
for c in range(3):
        if c == 0:
            indexconversiondic = {'Seeing my thought content in some way':'visual', 'Hearing my thought content in some way':'auditory', 'There is no difference':'no difference', 'I do not see or hear my thought content in any way':'no experience'}
            thisdf = pd.DataFrame(index=list(compare2qdic.values()), columns=list(indexconversiondic.values()))
            for qid in compare2qdic.keys():
                qdata = data[qid]
                qdf = 100*qdata.value_counts(normalize=True).reindex(list(indexconversiondic.keys())) #reorders
                qdf.index = list(indexconversiondic.values()) #replaces
                aspect = compare2qdic[qid]
                thisdf.loc[aspect,['visual','auditory','no difference','no experience']] = qdf
            title = 'Visual vs. auditory imagery'
            subtitle = 'Which modality most meets the criterion?'
            xlabel = 'criteria'
        if c == 1:
            qiddic = {'see':1,'hear':2,'smell':4,'taste':3,'touch':5,'no experience':6} # Dictionary order controls plotting order
            thisdf = pd.DataFrame(index=list(compare5qdic.values()), columns=list(qiddic.keys()))
            for q in compare5qdic.keys():
                aspect = compare5qdic[q]
                for modality in qiddic.keys():
                    qid = qiddic[modality]
                    modalitydata = data[f'{q}_{qid}']
                    prop = (modalitydata.count() - (modalitydata == '0').sum(axis=0))/modalitydata.count()
                    thisdf.loc[aspect, modality] = 100*prop
            title = 'Imagery comparisons across five senses'
            subtitle = 'Which modality most meets the criterion?'
            xlabel = 'criteria'
        if c == 2:
            qiddic = {'see':1,'hear':2,'smell':4,'taste':5,'touch':3,'move':6,'speak':7,'feel emotion':8,'experience phenomenon, but abstractly':9,'experience phenomenon, but not sure how':10,'never experience phenomenon':11,"don't know what it means to experience the phenomenon":12} # Dictionary order controls plotting order
            thisdf = pd.DataFrame(index=list(compare3qdic.values()), columns=list(qiddic.keys()))
            for q in compare3qdic.keys():
                aspect = compare3qdic[q]
                for modality in qiddic.keys():
                    qid = qiddic[modality]
                    if aspect == 'imagination': qid = qid + 1
                    modalitydata = data[f'{q}_{qid}']
                    prop = modalitydata.value_counts(normalize=True)['Agree']
                    thisdf.loc[aspect, modality] = 100*prop
            title = 'Experiences during thoughts, imagination, and dreams'
            subtitle = 'What do you experience in each phenomena?'#\n(multiple responses allowed)'
            xlabel = 'how phenomenon is experienced'
            
        # Create plots
        if c == 0:
            thisdf.index = [textwrap.fill(text,10,break_long_words=False) for text in thisdf.index] # Wraps text
            ax = thisdf.plot(kind='bar', color=colours2, rot=0, ax=axes[c], width=0.65)
            ax.text(s=f'n = {len(data)}', y=0.88, x=0.5, horizontalalignment='center', size=axlabelsize, transform=ax.transAxes)
            ax.legend(fontsize=axticklabelsize, frameon=False, labelcolor=colours2, handlelength=0, handletextpad=0, bbox_to_anchor=(0,0.1,1,1), ncol=2)
        if c == 1:
            thisdf.index = [textwrap.fill(text,10,break_long_words=False) for text in thisdf.index] # Wraps text
            ax = thisdf.plot(kind='bar', color=colours5, rot=0, ax=axes[c], width=0.825)
            ax.text(s=f'n = {len(data)}', y=0.88, x=0.5, horizontalalignment='center', size=axlabelsize, transform=ax.transAxes)
            ax.legend(fontsize=axticklabelsize, frameon=False, labelcolor=colours5, handlelength=0, handletextpad=0, bbox_to_anchor=(0,0.1,1,1), ncol=2)
        if c == 2:
            thisdf = thisdf.swapaxes('index','columns')
            thisdf.index = [textwrap.fill(text,10,break_long_words=False) for text in thisdf.index] # Wraps text
            ax = thisdf.plot(kind='bar', color=colours3, rot=0, ax=axes[c], width=0.825)
            ax.text(s=f'n = {len(data)}', y=0.88, x=0.5, horizontalalignment='center', size=axlabelsize, transform=ax.transAxes)
            ax.legend(fontsize=axticklabelsize, frameon=False, labelcolor=colours3, handlelength=0, handletextpad=0, loc='upper right') 
            ax.set_xticklabels(thisdf.index, fontsize=5.5)
            [l.set_fontsize(4) for l in ax.get_xticklabels()[-4:]]
            for i in range(len(ax.containers)):
                ax.bar_label(ax.containers[i], label_type='edge', fontsize=axticklabelsize, fmt='%.0f')
        if c in [0,1]:
            ax.set_xticklabels(thisdf.index, fontsize=5.5)
            for i in range(len(ax.containers)):
                ax.bar_label(ax.containers[i], label_type='edge', fontsize=4, fmt='%.0f')
          
        ax.set_ylim((0,124))
        ax.set_xlabel(xlabel, fontsize=axlabelsize, labelpad=5)
        ax.set_title(title, y=1.1, x=0.5, size=titlesize, pad=0, fontweight='bold')
        ax.spines.right.set_visible(False)
        ax.spines.top.set_visible(False)
        ax.spines.left.set_bounds(0, 100)
        ax.text(s=subtitle, y=.98, x=0.5, horizontalalignment='center', size=axlabelsize, transform=ax.transAxes)
        ax.text(s=letters[c], y=1.1, x=0, size=titlesize, fontweight='bold', transform=ax.transAxes)#x=-0.12 to put label far left
        ax.tick_params(axis='y', which='major', labelsize=axticklabelsize)
        ax.set_ylabel('% respondents', fontsize=axlabelsize, labelpad=5)
        
fig.tight_layout()
fig.savefig('Figure 5.png')

# %% [markdown]
*Note.* **A.** Participants made a single choice for each criterion. **B.** Participants could choose multiple responses if each applied equally. **C.** Participants could select each experience that applied in each phenomenon. Statistical comparisons are detailed in the supplementary materials (Supplementary S1.2 & S1.3)

# %% [markdown]
# Confidence: visual vs. auditory imagery

# %% [code]
data = prehall_filtered.copy()
confqdic = {'presence':'3',
          'automaticity':'6',
          'onset latency (cat)':'8',
          'consistency':'10',
          'proportion':'12',
          'peak moment':'14',
          'onset latency (num)':'17',
          'duration (num)':'19',
          'movement req.':'22',
          'required movement of head':'23#2_1',
          'required movement of brow':'23#2_2',
          'required movement of eyelids':'23#2_3',
          'required movement of eyes':'23#2_4',
          'required movement of ears':'23#2_5',
          'required movement of cheeks':'23#2_6',
          'required movement of nose':'23#2_7',
          'required movement of mouth':'23#2_8',
          'required movement of tongue':'23#2_9',
          'required movement of throat':'23#2_10',
          'required movement of arms/hands':'23#2_11',
          'required movement of torso':'23#2_12',
          'required movement of legs/feet':'23#2_13',
          'required movement of other body part':'25',
          'location control (eyes open)':'29',
          'location/position (categorical; eyes open)':'31',
          'spatial location (sideview; eyes open)':'33',
          'spatial location (topview; eyes open)':'35',
          'location control (eyes closed)':'38',
          'location/position (categorical; eyes closed)':'40',
          'spatial location (sideview; eyes closed)':'42',
          'spatial location (topview; eyes closed)':'44',
          'typicality':'46',
          'relative ability':'48',
          'terminology':'50'
            }

allvisconfs = pd.Series(dtype='object')
allaudconfs = pd.Series(dtype='object')
conversiondic = {'Completely unsure':0,'Mostly unsure':1,'Somewhat unsure':2,'Somewhat sure':3,'Mostly sure':4,'Completely sure':5}
for c, key in enumerate(confqdic.keys()):
    visq = 'Q4.' + confqdic[key]
    audq = 'Q5.' + confqdic[key]
    allvisconfs = pd.concat([allvisconfs, data[visq]])
    allaudconfs = pd.concat([allaudconfs, data[audq]])
    data.loc[:,[visq,audq]] = data[[visq,audq]].replace(conversiondic)
    diff = data[visq] - data[audq]
    diff.name = f'{key}'
    if c == 0:
        diffdf = pd.DataFrame(diff)
    else:
        diffdf[f'{key}'] = diff       

if allvisconfs.mode()[0] != allaudconfs.mode()[0] or list(conversiondic.keys())[int(allvisconfs.replace(conversiondic).median())] != list(conversiondic.keys())[int(allaudconfs.replace(conversiondic).median())]:
    modemeantext = f"""For visual questions, the modal confidence rating was "{allvisconfs.mode()[0]}" and median rating was "{list(conversiondic.keys())[int(allvisconfs.replace(conversiondic).median())]}". For auditory questions, the modal confidence rating was "{allaudconfs.mode()[0]}" and median rating was "{list(conversiondic.keys())[int(allaudconfs.replace(conversiondic).median())]}"."""
else:
    modemeantext = f"""For both visual and auditory questions, the modal confidence rating was "{allvisconfs.mode()[0]}" and median rating was "{list(conversiondic.keys())[int(allvisconfs.replace(conversiondic).median())]}" in each modality."""

# %% [code]
text=(f"""For each imagery question, participants were asked to indicate how sure they were about their response on a six-point scale from "Completely unsure" (0) to "Completely sure" (5). Here we analysed the confidence of participants when reporting on visual imagery relative to when reporting on auditory imagery for all {len(confqdic.keys())} questions with visual and auditory equivalents. {len(diffdf)} participants were analysed (including non-imagers and those classifying sensory thoughts as hallucinations). {modemeantext} To numerically compare confidence between modalities, ratings were converted to a 0-5 score, and the difference in confidence between modalities (visual confidence score - auditory confidence score) was calculated for each of the {len(confqdic.keys())} questions. Hence, there were {len(diffdf)*len(confqdic.keys())} possible confidence differences between modalities in total ({len(diffdf)} participants X {len(confqdic.keys())} questions). However, as the availability of some questions was conditional on previous answers, some participants did not complete each question for each modality. Hence, only {len(diffdf.stack())} differences were analysed. A two-tailed Wilcoxon signed-rank test showed that participant confidence was slightly higher when reporting on auditory imagery compared to visual imagery (mean difference = {round(diffdf.stack().mean(),1)}, standard deviation = {round(diffdf.stack().std(),1)}; {reportWilcoxon(diffdf.stack())}). Note that this comparison is only a coarse measure of the confidence difference between modalities. Statistical comparisons of the average paired difference in confidence between modalities for each underlying modality-specific question asked are shown in Supplementary S1.4."""
     )
md(text)

# %% [markdown]
# Confidence: imagination vs. thoughts vs. dreaming

# %% [code]
data=prehall_filtered.copy()
labeldic = {'1':'see',
            '2':'hear',
            '3':'touch',
            '4':'smell',
            '5':'taste',
            '6':'move',
            '7':'speak',
            '8':'emotion',
            '9':'have phenomenon, \nbut experience unlisted',
            '10':'have phenomenon, \nbut experience unsure',
            '11':'never have phenomenon',
            '12':"don't know what \nphenomenon is"
           }
qdic = {'thoughts':'Q7.1','imagination':'Q7.2','dreams':'Q7.3'}
conversiondic = {'Completely unsure':0,'Mostly unsure':1,'Somewhat unsure':2,'Somewhat sure':3,'Mostly sure':4,'Completely sure':5}
medians = {}
modes = {}

include = ''
props = pd.DataFrame(index=list(qdic.keys()),columns=list(labeldic.values()))
for c, phenomenon in enumerate(list(qdic.keys())):
    parentq = qdic[phenomenon]
    allconfs = pd.Series(dtype='object')
    for i in range(1,13):
        # To account for the presence of an attention check in 'imagination' questions
        checkcorrection = 0
        if phenomenon == 'imagination': checkcorrection = 1
        confq = f'{parentq}#2_{i+checkcorrection}'
        respq = f'{parentq}#1_{i+checkcorrection}'
        
        # Filter to include only those who agreed/disagreed that they experienced the sense/feeling/action
        if include == 'Agree' or include == 'Disagree':
            inclusionmask = data[respq] == include
            confdf = data.loc[inclusionmask,confq]
        else:
            confdf = data.loc[:,confq]
            
        allconfs = pd.concat((allconfs,confdf))
        confdf.replace(conversiondic, inplace=True)
        mean = confdf.mean()
        props.loc[phenomenon,labeldic[str(i)]] = mean
    mode = allconfs.value_counts().sort_values().index[-1]
    median = list(conversiondic.keys())[int(allconfs.replace(conversiondic).median())]
    medians[phenomenon] = median
    modes[phenomenon] = mode

# To elucidate differences between pairs of phenomena
# For each pair of phenomena, store a dic entry containing a df with an index of every participant, and columns being the difference between every matched pair of questions
diffdic = {}
meandiffs = {}
for pair in ['dreams-imagination','dreams-thoughts','imagination-thoughts']:
    phena = pair.split('-')[0]
    phenb = pair.split('-')[-1]
    # Include corrections accounting for the attention check in 'imagination' questions
    correctiona = 0
    correctionb = 0
    if phena == 'imagination': correctiona = 1
    if phenb == 'imagination': correctionb = 1
    
    diffdf = pd.DataFrame(index=data.index)
    for i in range(1,13):
        phenaq = f'{qdic[phena]}#2_{i+correctiona}'
        phenbq = f'{qdic[phenb]}#2_{i+correctionb}'
        aspect = labeldic[str(i)]
        diffqdf = data.loc[:,phenaq] - data.loc[:,phenbq]
        diffdf[f'{labeldic[str(i)]}'] = diffqdf
    diffdic[pair] = diffdf

# Check stats at the time of initial writing are still true    
if modes['thoughts'] != modes['imagination']: raise Exception('Not equal')
if medians['thoughts'] != medians['imagination']: raise Exception('Not equal')
if medians['dreams'] != modes['dreams']: raise Exception('Not equal')

# %% [code]
text=(f"""As per visual and auditory imagery questions, we also recorded and analysed the confidence ratings of participants when reporting on what type of sensory or non-sensory experience they had during their thoughts, imagination, and dreams. There were {len(confdf)*12} confidence ratings in total for each phenomenon ({len(confdf)} participants X 12 different types of experience to rate per phenomenon). For thoughts and imagination, the modal rating was "{modes['thoughts']}" and median rating was "{medians['thoughts']}" for each modality. For dreams, the modal and median confidence ratings were both "{modes['dreams']}". Scoring confidence ratings from 0 (Completely unsure) to 5 (Completely sure), we investigated whether confidence was higher in one phenomenon relative to another with two-tailed Wilcoxon signed-rank tests for the {len(confdf)*12} paired differences in confidence between each pair of phenomena. Confidence was significantly higher when reporting on dreams relative to imagination (mean difference = {round(diffdic['dreams-imagination'].stack().mean(),1)}; {reportWilcoxon(diffdic['dreams-imagination'].stack())}, and dreams relative to thoughts (mean difference = {round(diffdic['dreams-thoughts'].stack().mean(),1)}; {reportWilcoxon(diffdic['dreams-thoughts'].stack())}). Confidence was significantly higher when reporting on imagination relative to thoughts (mean difference = {round(diffdic['imagination-thoughts'].stack().mean(),1)}; {reportWilcoxon(diffdic['imagination-thoughts'].stack())}). Hence, respondents were most confident about the experiences of dreaming, followed by imagination, then thinking. Mean confidences for each individual aspect of each phenomenon are shown in Supplementary S1.5. Detailed comparisons of confidence within each phenomenon, and between phenomena, are reported in Supplementary S1.6 & S1.7"""
     )
md(text)

# %% [markdown]
# SUPPLEMENTARY

# %% [markdown]
## Supplementary S1.2

# %% [markdown]
Within-phenomenon paired comparisons of the proportion with which each aspect was reported during thoughts, imagination, and dreams

# %% [code]
masterbonf = (66*3) + (12*3)

data = filtered.copy()
compare3qdic = {'thoughts':'Q7.1#1', 'imagination':'Q7.2#1', 'dreams':'Q7.3#1'}
qiddic = {'see':1,'hear':2,'smell':4,'taste':5,'touch':3,'move':6,'speak':7,'feel emotion':8,'experience phenomenon, but abstractly':9,'experience phenomenon, but not sure how':10,'never experience phenomenon':11,"don't know what it means to experience the phenomenon":12} # 

# This is somewhat elegant code for only retaining unique combinations
# It skips pairs until after we have found matching pairs, then it keeps the pairs after that
statsdic = {'aspect 1':[], 'aspect 2':[]}
for aspect1 in qiddic.keys():
    skip = True
    for aspect2 in qiddic.keys():
        if skip:
            if aspect2 == aspect1:
                skip = False
            continue
        statsdic['aspect 1'].append(aspect1)
        statsdic['aspect 2'].append(aspect2)

# Do stats on each unique pair of confidence ratings within each phenomenon separately
for phenomenon in compare3qdic.keys():
    pvalues = []
    annotations = []
    q = compare3qdic[phenomenon]
    for aspect1, aspect2 in zip(statsdic['aspect 1'], statsdic['aspect 2']):
        qid1 = qiddic[aspect1]
        qid2 = qiddic[aspect2]
        if phenomenon == 'imagination':
            qid1 = qid1 + 1
            qid2 = qid2 + 1
        p = testSquareTable(data[[f'{q}_{qid1}',f'{q}_{qid2}']])[1].pvalue
        if p <= (0.05/masterbonf):
            annotations.append('**')
        elif p <= 0.05:
            annotations.append('*')
        else:
            annotations.append('')
        if p < 0.000001:
            p = 'p < .000001'
        elif p > 0.9999995:
            p = 'p > .999999'
        else:
            p = 'p = ' + f'{p:.6f}'[1:]
        pvalues.append(p)
    statsdic[f'{phenomenon}, p-value'] = pvalues
    statsdic[f'{phenomenon}, annotations'] = annotations

shorthanddic = {'experience phenomenon, but abstractly':'exp. abstract','experience phenomenon, but not sure how':'exp. but unsure','never experience phenomenon':'never exp.',"don't know what it means to experience the phenomenon":"don't know"} 

# Stats table created here and copied to clipboard
statsdf = pd.DataFrame(statsdic, index=[1+i for i in range(len(statsdic['aspect 1']))])
statsdf = statsdf.replace(shorthanddic)
statsdf.to_clipboard()
print(f"Number of comparisons here: {3*len(statsdic['aspect 1'])}, overall comparisons: {masterbonf}, Bonf. corrected critical p-value: {0.05/masterbonf}")
statsdf

# %% [markdown]
## Supplementary S1.3

# %% [markdown]
Between-phenomenon paired comparisons of the proportion with which each aspect was reported during thoughts, imagination, and dreams

# %% [code]
masterbonf = (66*3) + (12*3)

data = filtered.copy()
compare3qdic = {'thoughts':'Q7.1#1', 'imagination':'Q7.2#1', 'dreams':'Q7.3#1'}
qiddic = {'see':1,'hear':2,'smell':4,'taste':5,'touch':3,'move':6,'speak':7,'feel emotion':8,'experience phenomenon, but abstractly':9,'experience phenomenon, but not sure how':10,'never experience phenomenon':11,"don't know what it means to experience the phenomenon":12} # 

# This is somewhat elegant code for only retaining unique combinations
# It skips pairs until after we have found matching pairs, then it keeps the pairs after that
statsdic = {'aspect':list(qiddic)}

# Do stats on each unique pair of confidence ratings within each phenomenon separately
for phenpair in [('thoughts','imagination'),('thoughts','dreams'),('imagination','dreams')]:
    pvalues = []
    annotations = []
    phen1, phen2 = phenpair
    for aspect in statsdic['aspect']:
        q1 = compare3qdic[phen1]
        q2 = compare3qdic[phen2]
        qid1 = qiddic[aspect]
        qid2 = qiddic[aspect]
        if phen1 == 'imagination': qid1 = qid1 + 1
        if phen2 == 'imagination': qid2 = qid2 + 1
        p = testSquareTable(data[[f'{q1}_{qid1}',f'{q2}_{qid2}']])[1].pvalue
        if p <= (0.05/masterbonf):
            annotations.append('**')
        elif p <= 0.05:
            annotations.append('*')
        else:
            annotations.append('')
        if p < 0.000001:
            p = 'p < .000001'
        elif p > 0.9999995:
            p = 'p > .999999'
        else:
            p = 'p = ' + f'{p:.6f}'[1:]
        pvalues.append(p)
    statsdic[f'{phen1} vs {phen2}, p-value'] = pvalues
    statsdic[f'{phen1} vs {phen2}, annotations'] = annotations

shorthanddic = {'experience phenomenon, but abstractly':'exp. abstract','experience phenomenon, but not sure how':'exp. but unsure','never experience phenomenon':'never exp.',"don't know what it means to experience the phenomenon":"don't know"} 

# Stats table created here and copied to clipboard
statsdf = pd.DataFrame(statsdic, index=[1+i for i in range(len(statsdic['aspect']))])
statsdf = statsdf.replace(shorthanddic)
statsdf.to_clipboard()
print(f"Number of comparisons here: {len(statsdic['aspect'])}, overall comparisons: {masterbonf}, Bonf. corrected critical p-value: {0.05/masterbonf}")
statsdf

# %% [markdown]
## Supplementary S1.4

# %% [markdown]
Mean confidence ratings for modality-specific questions

# %% [code]
# Supplementary for "Confidences (visual versus auditory)" 
data = prehall_filtered.copy()
confqdic = {'prevalence':'3',
          'intentionality':'6',
          'coarse onset latency':'8',
          'consistency':'10',
          'proportion':'12',
          'peak':'14',
          'precise onset latency':'17',
          'precise duration':'19',
          'movement-dependence':'22',
          'body-part required: head':'23#2_1',
          'body-part required: brow':'23#2_2',
          'body-part required: eyelids':'23#2_3',
          'body-part required: eyes':'23#2_4',
          'body-part required: ears':'23#2_5',
          'body-part required: cheeks':'23#2_6',
          'body-part required: nose':'23#2_7',
          'body-part required: mouth':'23#2_8',
          'body-part required: tongue':'23#2_9',
          'body-part required: throat':'23#2_10',
          'body-part required: arms/hands':'23#2_11',
          'body-part required: torso':'23#2_12',
          'body-part required: legs/feet':'23#2_13',
          'body-part required: other':'25',
          'location control (eyes open)':'29',
          'coarse position (eyes open)':'31',
          'precise spatial position (sideview; eyes open)':'33',
          'precise spatial position (topview; eyes open)':'35',
          'location control (eyes closed)':'38',
          'coarse position (eyes closed)':'40',
          'precise spatial position (sideview; eyes closed)':'42',
          'precise spatial position (topview; eyes closed)':'44',
          'typicality':'46',
          'ability':'48',
          'terminology':'50',
          'cross-modal':('Q6.4','Q6.2'),
          'dual-modal':('Q6.8','Q6.6')
            }

# Calculate differences between modalities
conversiondic = {'Completely unsure':0,'Mostly unsure':1,'Somewhat unsure':2,'Somewhat sure':3,'Mostly sure':4,'Completely sure':5}
for c, key in enumerate(confqdic.keys()):
    if key in ['cross-modal','dual-modal']:
        visq = confqdic[key][0]
        audq = confqdic[key][1]
    else:
        visq = 'Q4.' + confqdic[key]
        audq = 'Q5.' + confqdic[key]
    data.loc[:,[visq,audq]] = data[[visq,audq]].replace(conversiondic)
    thisdata = data.loc[:,[visq,audq]]
    thisdata = thisdata.dropna(how='any') # Remove nans for questions where they did one spatial map question but not another
    viscounts = thisdata[visq].value_counts(normalize=True)
    audcounts = thisdata[audq].value_counts(normalize=True)
    vismean = thisdata[visq].mean()
    audmean = thisdata[audq].mean()
    diffs = thisdata[visq] - thisdata[audq]
    diff = diffs.mean(axis=0)
    vismeanerror = thisdata[visq].std()/((len(thisdata[visq]))**0.5)
    audmeanerror = thisdata[audq].std()/((len(thisdata[audq]))**0.5)
    differror = diffs.std()/((len(diffs))**0.5)
    wilc = wilcoxon(diffs).pvalue
    p = round(wilc,3)
    if p < 0.001:
        ptext = 'p < .001'
    elif p > 0.9995:
        ptext = 'p > .999'
    else:
        ptext = 'p = ' + f'{p:.3f}'[1:]
    qdata = pd.Series({'name':f'{key}\n{visq}, {audq}','viscounts':viscounts,'vismean':vismean,'vismeanerror':vismeanerror,'audmean':audmean,'audmeanerror':audmeanerror,'audcounts':audcounts, 'pvalue':wilc, 'ptext':ptext, 'diff':diff,'differror':differror})
    if c==0:
        masterdf = qdata
    else:
        masterdf = pd.concat((masterdf,qdata), axis=1)
masterdf = masterdf.T
masterdf = masterdf.set_index('name')
masterdf = masterdf.sort_values(by='diff')

# Plot
fig, axes = plt.subplots(ncols=2,nrows=1, figsize=(12,12), dpi=600)
masterdf['diff'].plot(kind='barh', xlim=(-1,0.6), ax=axes[1], color='grey', zorder=2, xerr=masterdf['differror'], error_kw=dict(ecolor='black', capsize=0, capthick=0.5, lw=0.5))
masterdf[['audmean','vismean']].plot(kind='barh', ax=axes[0], xlim=(3,4.6), color=['orchid','cornflowerblue'], zorder=2, xerr=[masterdf['audmeanerror'],masterdf['vismeanerror']], error_kw=dict(ecolor='black', capsize=0, capthick=0.5, lw=0.5))

# Colour annotations
ylabelcols = []
for y in axes[1].get_yticks():
    barcol = 'grey'
    textcol = 'black'
    if masterdf.at[masterdf.index[y],'pvalue'] <= 0.05:
        barcol = 'darkorange'
        textcol = 'darkorange'
    if masterdf.at[masterdf.index[y],'pvalue'] <= (0.05/len(masterdf)):
        barcol = 'red'
        textcol = 'red'
    ylabelcols.append(textcol)
    #axes[1].patches[y].set_facecolor(col)

axes[0].grid(visible=True, which='major', axis='y', zorder=1, alpha=0.5)
axes[1].grid(visible=True, which='major', axis='y', zorder=1, alpha=0.5)
axes[0].tick_params(axis='y', length=0)
axes[1].tick_params(axis='y', length=0, labelright=True)
axes[0].set_yticklabels(masterdf.index, size=8)
axes[1].set_yticklabels(masterdf['ptext'])
axes[1].yaxis.tick_right()
axes[1].yaxis.set_label_position("right")
yticklabels0 = axes[0].get_yticklabels()
yticklabels1 = axes[1].get_yticklabels()
for i in range(len(yticklabels0)):
    yticklabels0[i].set_color(ylabelcols[i])
    yticklabels1[i].set_color(ylabelcols[i])
axes[0].set_yticklabels(yticklabels0)
axes[0].set_xlabel('mean confidence')
axes[1].set_xlabel('mean paired confidence diff. (vis. confidence - aud. confidence)\nmore confident in auditory <---> more confident in visual')
axes[0].set_ylabel('question')
axes[1].set_ylabel('')
axes[0].spines.right.set_visible(False)
axes[1].spines.right.set_visible(False)
axes[0].spines.top.set_visible(False)
axes[1].spines.top.set_visible(False)
axes[0].spines.left.set_visible(False)
axes[1].spines.left.set_visible(False)

handles, labels = axes[0].get_legend_handles_labels()
axes[0].legend(reversed(handles), ['visual','auditory'], loc='upper right', frameon=False)
fig.tight_layout()

# Wholeness (visual only) confidence mean:
wholenessmean = data.loc[:,["Q4.53"]].replace(conversiondic).mean()[0]
wholenesssem = ((data.loc[:,["Q4.53"]].replace(conversiondic).std())/len(data.loc[:,["Q4.53"]]))[0]
print(f'Number of Qs: {len(masterdf)}')
print(f'Wholeness (visual only) confidence; mean: {wholenessmean:.2f}, SEM: {wholenesssem:.4f}')

# %% [markdown]
## Supplementary S1.5

# %% [markdown]
Mean confidence ratings for thoughts, imagination, and dreams

# %% [code]
data = filtered.copy()
compare3qdic = {'thoughts':'Q7.1#2', 'imagination':'Q7.2#2', 'dreams':'Q7.3#2'}
conversiondic = {'Completely unsure':0,'Mostly unsure':1,'Somewhat unsure':2,'Somewhat sure':3,'Mostly sure':4,'Completely sure':5}
qiddic = {'see':1,'hear':2,'smell':4,'taste':5,'touch':3,'move':6,'speak':7,'feel emotion':8,'experience phenomenon, but abstractly':9,'experience phenomenon, but not sure how':10,'never experience phenomenon':11,"don't know what it means to experience the phenomenon":12} # 

# Make figure
fig = plt.figure(dpi=600, figsize=(6,4))
ax = fig.add_subplot()
cmap = mpc.LinearSegmentedColormap.from_list("", ["white","mediumslateblue"])
colours3 = [cmap(0.4),cmap(0.7),cmap(0.99)]
thisdf = pd.DataFrame(index=list(compare3qdic.keys()), columns=list(qiddic.keys()))
thiserrordf = pd.DataFrame(index=list(compare3qdic.keys()), columns=[entry + ' (error)' for entry in qiddic.keys()])

# Get values
for phenomenon in compare3qdic.keys():
    q = compare3qdic[phenomenon]
    for aspect in qiddic.keys():
        qid = qiddic[aspect]
        if phenomenon == 'imagination': qid = qid + 1
        aspectdata = data[f'{q}_{qid}'].replace(conversiondic)
        meanconf = aspectdata.mean()
        errorconf = aspectdata.std()/(len(aspectdata)**0.5)
        thisdf.loc[phenomenon, aspect] = meanconf
        thiserrordf.loc[phenomenon, aspect + ' (error)'] = errorconf

thisdf = thisdf.swapaxes('index','columns')
thisdf.index = [textwrap.fill(text,10,break_long_words=False) for text in thisdf.index] # Wraps text
ax = thisdf.plot(kind='bar', color=colours3, rot=0, ax=ax, width=0.825, yerr=thiserrordf.values, error_kw=dict(ecolor='black', capsize=0, capthick=0.5, lw=0.5))
ax.legend(fontsize=None, frameon=False, labelcolor=colours3, handlelength=0, handletextpad=0, loc='upper center', ncol=3) 
ax.set_xticklabels(thisdf.index, fontsize=6.5)
[l.set_fontsize(4) for l in ax.get_xticklabels()[-4:]]
ax.set_ylim((3.5,5.19))
ax.set_xlabel('aspect of experience', fontsize=None, labelpad=5)
ax.spines.right.set_visible(False)
ax.spines.top.set_visible(False)
ax.spines.left.set_bounds(3.5, 5)
ax.set_ylabel('mean confidence', fontsize=None) #labelpad=5
fig.tight_layout()

# %% [markdown]
## Supplementary S1.6

# %% [markdown]
Within-phenomenon paired confidence comparisons for thoughts, imagination, and dreams

# %% [code]
masterbonf = (66*3) + (12*3)

data = filtered.copy()
compare3qdic = {'thoughts':'Q7.1#2', 'imagination':'Q7.2#2', 'dreams':'Q7.3#2'}
conversiondic = {'Completely unsure':0,'Mostly unsure':1,'Somewhat unsure':2,'Somewhat sure':3,'Mostly sure':4,'Completely sure':5}
qiddic = {'see':1,'hear':2,'smell':4,'taste':5,'touch':3,'move':6,'speak':7,'feel emotion':8,'experience phenomenon, but abstractly':9,'experience phenomenon, but not sure how':10,'never experience phenomenon':11,"don't know what it means to experience the phenomenon":12} # 

# This is somewhat elegant code for only retaining unique combinations
# It skips pairs until after we have found matching pairs, then it keeps the pairs after that
statsdic = {'aspect 1':[], 'aspect 2':[]}
for aspect1 in qiddic.keys():
        skip = True
        for aspect2 in qiddic.keys():
            if skip:
                if aspect2 == aspect1:
                    skip = False
                continue
            statsdic['aspect 1'].append(aspect1)
            statsdic['aspect 2'].append(aspect2)

# Do stats on each unique pair of confidence ratings within each phenomenon separately
for phenomenon in compare3qdic.keys():
    pvalues = []
    annotations = []
    q = compare3qdic[phenomenon]
    for aspect1, aspect2 in zip(statsdic['aspect 1'], statsdic['aspect 2']):
        qid1 = qiddic[aspect1]
        qid2 = qiddic[aspect2]
        if phenomenon == 'imagination':
            qid1 = qid1 + 1
            qid2 = qid2 + 1
        p = wilcoxon(data[f'{q}_{qid1}'].replace(conversiondic), data[f'{q}_{qid2}'].replace(conversiondic)).pvalue
        if p <= (0.05/masterbonf):
            annotations.append('**')
        elif p <= 0.05:
            annotations.append('*')
        else:
            annotations.append('')
        if p < 0.000001:
            p = 'p < .000001'
        elif p > 0.9999995:
            p = 'p > .999999'
        else:
            p = 'p = ' + f'{p:.6f}'[1:]
        pvalues.append(p)
    statsdic[f'{phenomenon}, p-value'] = pvalues
    statsdic[f'{phenomenon}, annotations'] = annotations

shorthanddic = {'experience phenomenon, but abstractly':'exp. abstract','experience phenomenon, but not sure how':'exp. but unsure','never experience phenomenon':'never exp.',"don't know what it means to experience the phenomenon":"don't know"} 

# Stats table created here and copied to clipboard
statsdf = pd.DataFrame(statsdic, index=[1+i for i in range(len(statsdic['aspect 1']))])
statsdf = statsdf.replace(shorthanddic)
statsdf.to_clipboard()
print(f"Number of comparisons here: {3*len(statsdic['aspect 1'])}, overall comparisons: {masterbonf}, Bonf. corrected critical p-value: {0.05/masterbonf}")
statsdf

# %% [markdown]
## Supplementary S1.7

# %% [markdown]
Between-phenomenon paired confidence comparisons for thoughts, imagination, and dreams

# %% [code]
masterbonf = (66*3) + (12*3)

data = filtered.copy()
compare3qdic = {'thoughts':'Q7.1#2', 'imagination':'Q7.2#2', 'dreams':'Q7.3#2'}
conversiondic = {'Completely unsure':0,'Mostly unsure':1,'Somewhat unsure':2,'Somewhat sure':3,'Mostly sure':4,'Completely sure':5}
qiddic = {'see':1,'hear':2,'smell':4,'taste':5,'touch':3,'move':6,'speak':7,'feel emotion':8,'experience phenomenon, but abstractly':9,'experience phenomenon, but not sure how':10,'never experience phenomenon':11,"don't know what it means to experience the phenomenon":12} # 

# This is somewhat elegant code for only retaining unique combinations
# It skips pairs until after we have found matching pairs, then it keeps the pairs after that
statsdic = {'aspect':list(qiddic)}

# Do stats on each unique pair of confidence ratings within each phenomenon separately
for phenpair in [('thoughts','imagination'),('thoughts','dreams'),('imagination','dreams')]:
    pvalues = []
    annotations = []
    phen1, phen2 = phenpair
    for aspect in statsdic['aspect']:
        q1 = compare3qdic[phen1]
        q2 = compare3qdic[phen2]
        qid1 = qiddic[aspect]
        qid2 = qiddic[aspect]
        if phen1 == 'imagination': qid1 = qid1 + 1
        if phen2 == 'imagination': qid2 = qid2 + 1
        p = wilcoxon(data[f'{q1}_{qid1}'].replace(conversiondic), data[f'{q2}_{qid2}'].replace(conversiondic)).pvalue
        if p <= (0.05/masterbonf):
            annotations.append('**')
        elif p <= 0.05:
            annotations.append('*')
        else:
            annotations.append('')
        if p < 0.000001:
            p = 'p < .000001'
        elif p > 0.9999995:
            p = 'p > .999999'
        else:
            p = 'p = ' + f'{p:.6f}'[1:]
        pvalues.append(p)
    statsdic[f'{phen1} vs {phen2}, p-value'] = pvalues
    statsdic[f'{phen1} vs {phen2}, annotations'] = annotations

shorthanddic = {'experience phenomenon, but abstractly':'exp. abstract','experience phenomenon, but not sure how':'exp. but unsure','never experience phenomenon':'never exp.',"don't know what it means to experience the phenomenon":"don't know"} 

# Stats table created here and copied to clipboard
statsdf = pd.DataFrame(statsdic, index=[1+i for i in range(len(statsdic['aspect']))])
statsdf = statsdf.replace(shorthanddic)
statsdf.to_clipboard()
print(f"Number of comparisons here: {len(statsdic['aspect'])}, overall comparisons: {masterbonf}, Bonf. corrected critical p-value: {0.05/masterbonf}")
statsdf

# %% [markdown]
## Supplementary S1.8

# %% [markdown]
Open-text responses to select questions (including from respondents who labelled sensory thoughts as hallucinations)

# %% [code]
# Free text responses
data = prehall_filtered.copy()

def reportOpenText(qid, title):
    print(f'{title} ({qid})')
    c = 1
    for text in data[qid][data[qid] != '-99']:
        if not isinstance(text, str):
            continue
        if text.isspace():
            continue
        text = text.replace("\n", " ")
        print(f'{c}. ',text)
        c += 1
    print()

reportOpenText('Q4.24_3_TEXT', 'Other body parts required: visual')
reportOpenText('Q5.24_3_TEXT', 'Other body parts required: auditory')
reportOpenText('Q4.49_4_TEXT', 'Alternative terminologies: visual')
reportOpenText('Q5.49_4_TEXT', 'Alternative terminologies: auditory')
reportOpenText('Q4.54', 'Extra comments: visual')
reportOpenText('Q5.51', 'Extra comments: auditory')
reportOpenText('Q6.10', 'Extra comments: visual-audio interactions')
reportOpenText('Q7.4', 'Extra comments: thoughts, imaginings, dreams')
reportOpenText('Q8.24', 'Extra comments: direct multimodal comparisons')
reportOpenText('Q9.14', 'Thinking vs. imagining: is there a difference?')

# %% [markdown]
# Export notebook

# %% [code]
infile = "Supplementary S3.ipynb"
outfile = "Exported S3.html"
temp = "styletemplate.tpl"
!jupyter nbconvert --to html --no-input --no-prompt --template full "{infile}" --TagRemovePreprocessor.remove_cell_tags='hideme' --output-dir="./" --output "{outfile}"
```
