## Supplementary figures and images for "Comparing mental imagery experiences across visual, auditory, and other sensory modalities"

### Supplementary S4.png

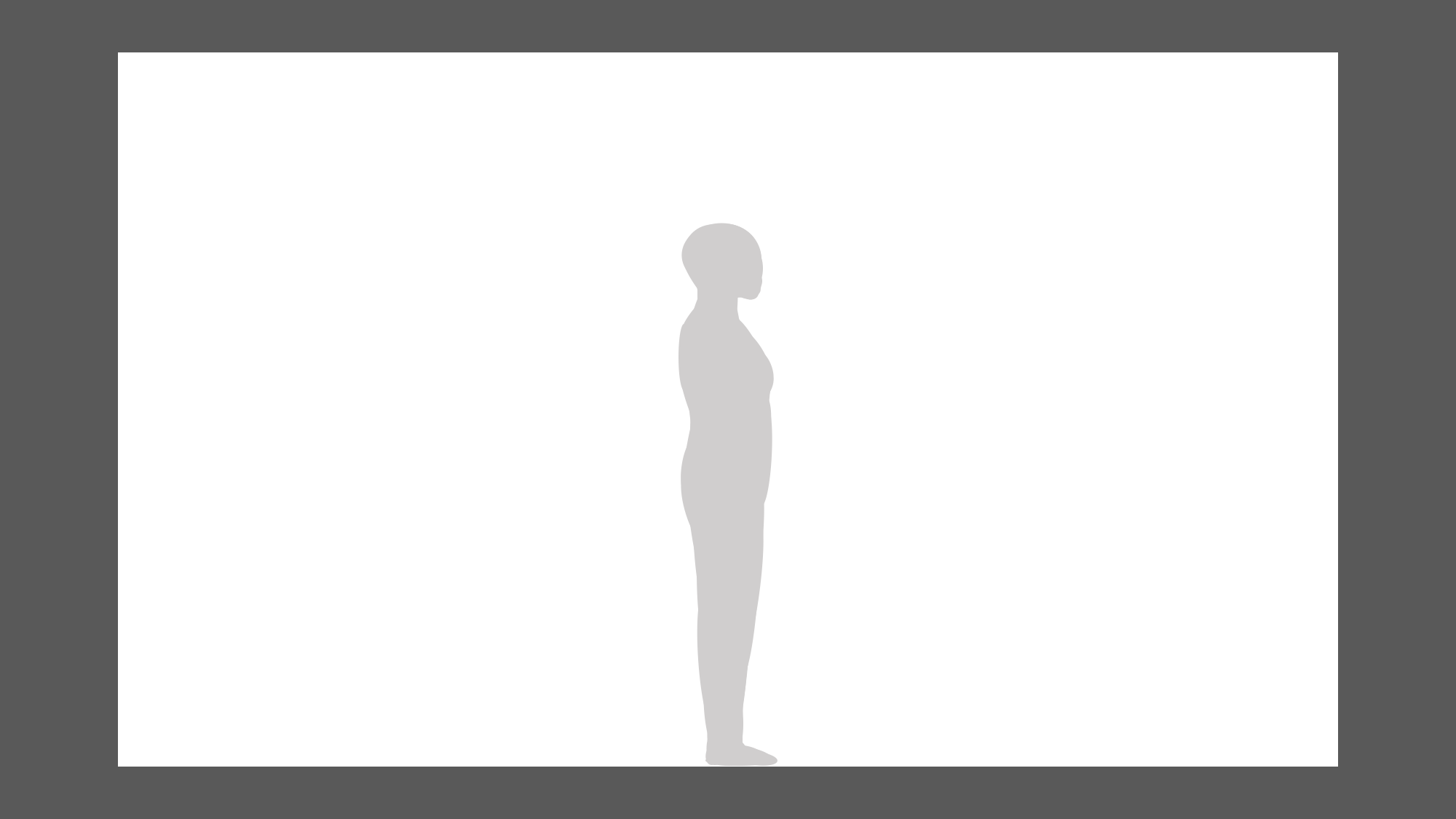

### Supplementary S5.png

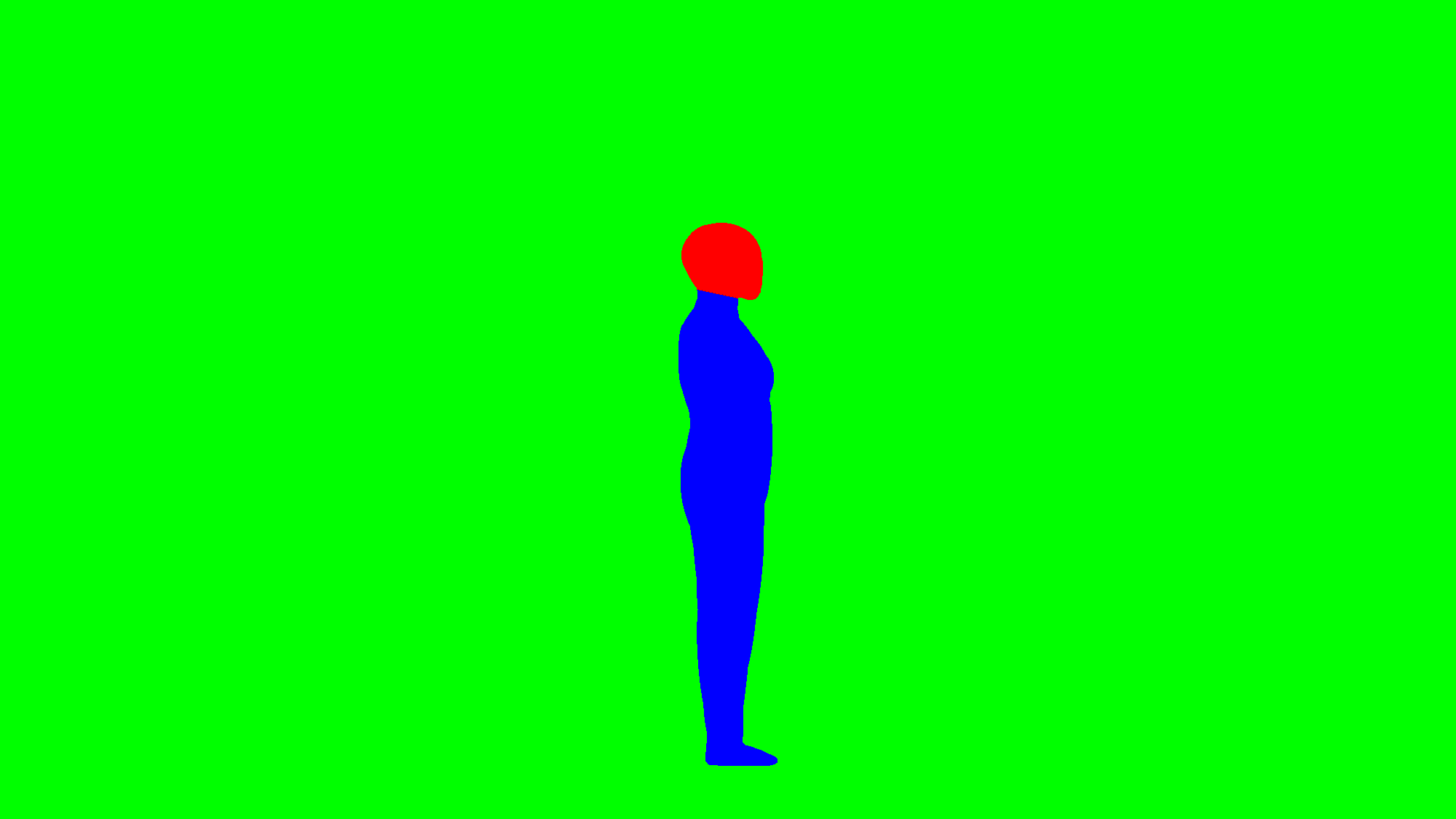

### Supplementary S6.png

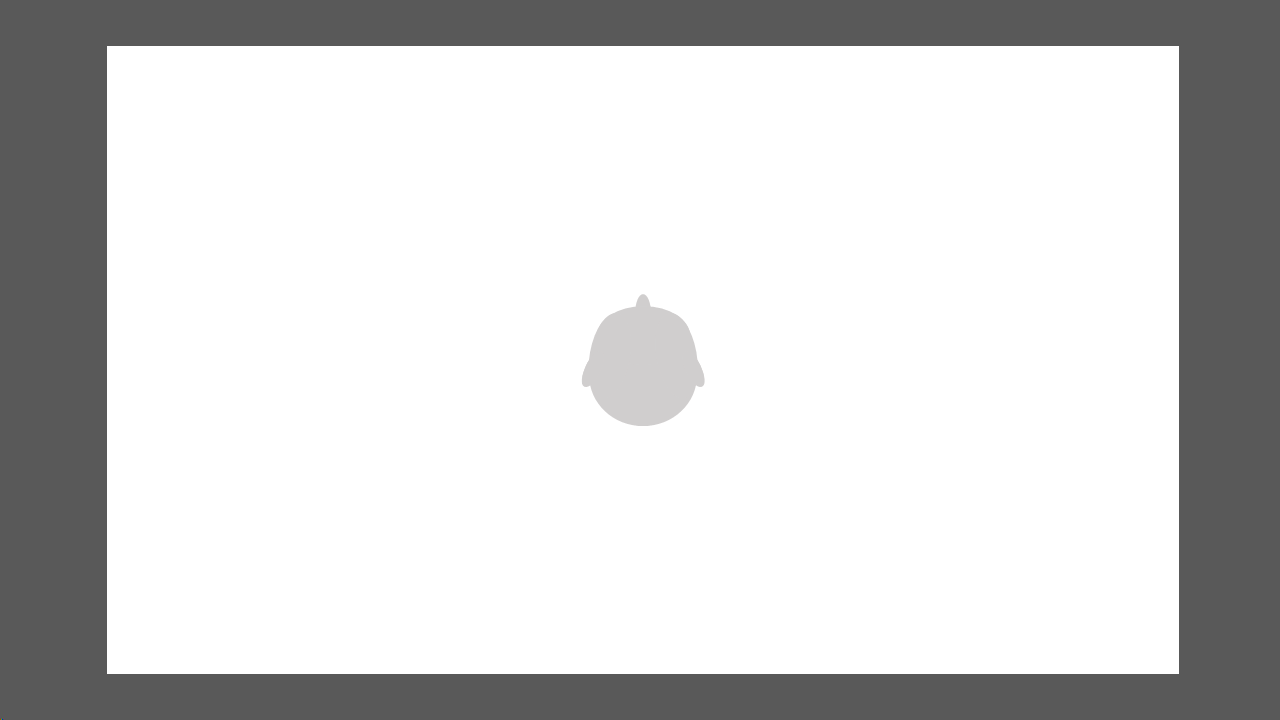

### Supplementary S7.png

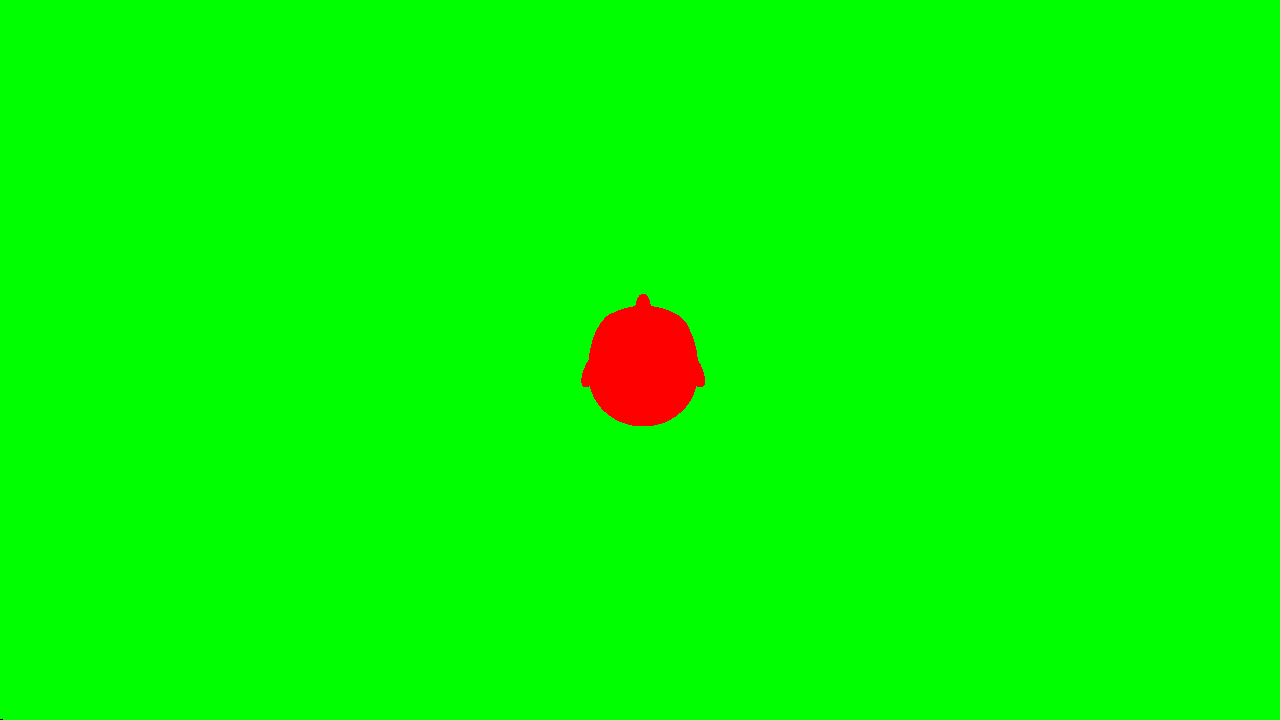
